# Imaging high-frequency voltage dynamics in multiple neuron classes of behaving mammals

**DOI:** 10.1101/2024.08.15.607428

**Authors:** Simon Haziza, Radosław Chrapkiewicz, Yanping Zhang, Vasily Kruzhilin, Jane Li, Jizhou Li, Geoffroy Delamare, Rachel Swanson, György Buzsáki, Madhuvanthi Kannan, Ganesh Vasan, Michael Z. Lin, Hongkui Zeng, Tanya L. Daigle, Mark J. Schnitzer

## Abstract

Fluorescent genetically encoded voltage indicators report transmembrane potentials of targeted cell-types. However, voltage-imaging instrumentation has lacked the sensitivity to track spontaneous or evoked high-frequency voltage oscillations in neural populations. Here we describe two complementary TEMPO voltage-sensing technologies that capture neural oscillations up to ∼100 Hz. Fiber-optic TEMPO achieves ∼10-fold greater sensitivity than prior photometry systems, allows hour-long recordings, and monitors two neuron-classes per fiber-optic probe in freely moving mice. With it, we uncovered cross-frequency-coupled theta- and gamma-range oscillations and characterized excitatory-inhibitory neural dynamics during hippocampal ripples and visual cortical processing. The TEMPO mesoscope images voltage activity in two cell-classes across a ∼8-mm-wide field-of-view in head-fixed animals. In awake mice, it revealed sensory-evoked excitatory-inhibitory neural interactions and traveling gamma and 3-7 Hz waves in the visual cortex, and previously unreported propagation directions for hippocampal theta and beta waves. These technologies have widespread applications probing diverse oscillations and neuron-type interactions in healthy and diseased brains.

## INTRODUCTION

Fluorescence imaging studies of neural activity using genetically encoded voltage indicators (GEVIs) have mainly focused on detecting the spiking dynamics of identified neuron classes at cellular resolution.^1–6^ However, complementary to such approaches, fiber-optic studies of subthreshold voltage activity across populations of genetically identified neuron-types can yield insights about neural circuit dynamics in freely behaving rodents.^7–9^ Neural population-level imaging studies have also used GEVIs to monitor the spatiotemporal dynamics of sensory-evoked potentials or slow voltage oscillations and activity in anesthetized or awake head-fixed animals.^10–13^ A key limitation, however, is that prior instruments for optical voltage-sensing in either freely moving or head-restrained mammals have lacked the sensitivity to detect rhythms at frequencies above the theta range (∼5–9 Hz) without substantial averaging across experimental trials.^7,9,13^ Due to this technology gap, how specific neuron-types shape the spatiotemporal dynamics of high-frequency oscillations (*i.e*., ≥10 Hz) remains poorly understood.

Addressing this challenge is crucial, because high-frequency oscillations appear to be integral to many important brain processes. Neural oscillations in the beta (∼15–30 Hz) and gamma (∼35–100 Hz) frequency bands are implicated in attention,^14,15^ motor control,^16,17^ memory^18,19,20,21^ and sensory processing.^22,23^ Impairments of high-frequency oscillations arise in brain disease, including cognitive disorders,^24^ schizophrenia,^25^ epilepsy,^26–28^ and Alzheimer’s^29–31^ and Parkinson’s diseases.^32–34^ Well-known hypotheses posit that high-frequency oscillations emerge via interactions between excitatory and inhibitory cells, or between coupled sets of mutually inhibitory interneurons.^9,35–37^ Available evidence points to inhibitory interneurons, especially fast-spiking interneurons,^22,38,39^ as being important for high-frequency rhythms, but far more work is needed to clarify how different neuron-types and both local and long-distance neural interactions may shape rhythmic synchronization.^9,14,21,35–37^

To enable studies that dissect how different neuron classes contribute to high-frequency oscillations and waves, this paper describes several new tools and techniques based on the TEMPO (**T**ransmembrane **E**lectrical **M**easurement **P**erformed **O**ptically) approach that we introduced previously in a fiber-optic format.^7^ TEMPO captures the aggregate, population-level voltage dynamics of specific neuron-types in behaving mammals. The motivation for such measurements is that, just as in electrophysiology, optical studies of neural population dynamics can reveal collective activity patterns that may not be discernible from single-cell activity traces.

TEMPO has 4 basic ingredients: (1) Use of one or more GEVIs to track voltage activity in chosen neuron-types, which may be targeted using viruses, transgenic animals, or other means of selective labeling; (2) Use of a reference fluor that is not voltage-sensitive to track hemodynamic, brain motion and other artifacts that impact the detection of voltage-sensitive fluorescence signals; (3) A dual-color fluorescence measurement apparatus; (4) Computational unmixing of the artifactual signals from the records of neural voltage activity.

Since both TEMPO and extracellular local electric field potential (LFP) recordings measure neural population voltage activity, it might be tempting to view the two as analogous; however, LFP recordings generally reflect the contributions of multiple unidentified cell-types, are influenced by electrode shape, orientation and composition, and comprise a time-varying, unknown mixture of signal sources across multiple length scales—from currents within ∼0.1 mm of the electrode to volume-conducted signals originating up to 1 cm away.^40^ By comparison, TEMPO probes the local intracellular potentials of user-selected cell-types and is free from volume-conduction effects.

Here, we present an ultrasensitive form of fiber-optic TEMPO for use in freely moving animals and a TEMPO mesoscope for voltage-imaging over tissue areas up to 8 mm in diameter in head-fixed behaving animals. Both instruments are unprecedented in that they can track high-frequency neural voltage oscillations (up to ∼100 Hz) without trial-averaging. In addition, we created a transgenic Cre-dependent reporter mouse line expressly designed for TEMPO studies. While most of our investigations relied on viral expression strategies, this reporter line provides a convenient option for uniform dual-fluor labeling and high reproducibility across animals without virus use. Finally, we describe a novel algorithm to unmix biological artifacts and noise from voltage signals while preserving the fidelity of photon shot noise-limited, high-frequency voltage dynamics.

Our fiber-optic TEMPO recordings rely on a photometry apparatus that we call uSMAART (**u**ltra-**S**ensitive **M**easurement of **A**ggregate **A**ctivity in **R**estricted cell-**T**ypes). We distinguish the TEMPO concept from the uSMAART apparatus, as the former need not be implemented in optical fiber, and the latter is an ultrasensitive fiber-optic system that does more than just voltage-sensing, including monitoring fluorescent [Ca^2+^] or neuromodulator reporters. The uSMAART apparatus has dual-channel fluorescence detection, which, when used for TEMPO, allows voltage-sensing of two targeted neuron classes per fiber-optic probe while also tracking the emissions of a reference fluor. uSMAART has ∼10-fold greater sensitivity than prior photometry systems for use in freely moving animals, and, when combined with the best available GEVI for sensing subthreshold voltages, allows a ∼100-fold improvement over prior fiber-optic studies of voltage dynamics.

With uSMAART, we measured population voltage activity at frequencies up to the gamma band (∼35–100 Hz) in freely behaving mice, in sparse and dense cell-types, without trial-averaging. Illustrating the importance of this new capability, we uncovered cross-frequency coupling (CFC) between pairs of voltage rhythms of different frequencies. In CFC, a lower frequency oscillation modulates the attributes of a higher frequency rhythm. CFC has previously been observed in electrical recordings in multiple animal species, and different forms of CFC associate with various cognitive states and diseases.^27,32,41–45^ uSMAART allowed the first observations of CFC within the transmembrane potentials of specific neuron-types, namely delta frequency (0.5–4 Hz) modulations of low- (30–60 Hz) and high-gamma (70–100 Hz) rhythms in parvalbumin-expressing (PV) interneurons in the visual cortex of anesthetized mice, as well as theta (5–9 Hz) modulations of gamma (40–70 Hz) rhythms in PV cells in the hippocampus of freely moving mice. By using uSMAART and two GEVIs, we monitored two neuron-types at once, which revealed the differential voltage dynamics of excitatory and inhibitory cell-types in visual cortex and hippocampus.

Since fiber photometry measurements are limited to signals from just below the fiber tip, we created a complementary TEMPO mesoscope to image the population voltage dynamics of two neuron-types at once in head-restrained mice. The mesoscope can image low-gamma (30–60 Hz) activity across a ∼50 mm^2^ field-of-view or take high-speed (300 Hz) snapshots of wave propagation over a ∼21 mm^2^ area, at a spatial resolution >10 times finer than those provided by the densest electrode arrays for electrocorticography (ECoG).^46,47^ With this system, we imaged traveling neocortical voltage waves with CFC synchronized across the delta and gamma bands in PV and layer 2/3 pyramidal cells of anesthetized mice. In awake mice, we found paired trains of visually evoked waves in these cell-types; visual stimulation elicited traveling gamma waves, followed by traveling 3-7 Hz waves at stimulus offset. In hippocampus, TEMPO imaging of PV interneurons provided the first evidence that hippocampal beta rhythms are actually traveling waves and can propagate in a pair of orthogonal anatomic directions. We also performed the first imaging studies of pathway-specific neuronal voltage oscillations by using axonal labeling to target hippocampal pyramidal neurons with a specific projection pattern. Further, our imaging studies of hippocampal theta waves revealed these waves can propagate bidirectionally along the CA1-CA3 axis, which was previously unknown. Finally, by using two GEVIs, we imaged the joint dynamics of visual cortical excitatory and inhibitory neural 3-7 Hz waves evoked in response to visual stimulation.

Overall, the instruments, reporter mouse and analytics introduced here allow a wide range of previously infeasible studies of high-frequency (*i.e.,* ∼10–100 Hz) transmembrane voltage dynamics in one or two neuron-classes at once, in freely moving or head-fixed behaving animals.

## RESULTS

### Ultra-sensitive fiber photometry system

To capture high-frequency oscillations up to ∼100 Hz in freely moving mammals, we created the ultra-sensitive fiber photometry system uSMAART. Prior fiber apparatus have been unable to capture optical voltage traces revealing high-frequency neural oscillations in freely behaving mice. This is challenging, because (i) fiber photometry measurements from GEVIs usually involve very weak fluorescence signals (∼50-250 pW), (ii) neural oscillations at higher frequencies afford fewer GEVI signal photons per oscillation cycle, (iii) GEVIs alter their emissions by only ∼0.1–1% per mV of voltage change, (iv) hemodynamics and tissue motion typically induce artifactual signals of this magnitude or greater, and (v) fiber-optic apparatus generally have multiple noise sources, including light sources, photodetectors, amplifiers and fiber autofluorescence.

Our own past work on fiber-optic TEMPO came nearest to the landmark achieved here but fell short in key respects.^7,9^ Specifically, our studies of high-frequency oscillations used head-fixed mice and averages over hundreds of trials to attain statistical conclusions.^7^ A follow-up study, which also involved substantial trial-averaging, used freely behaving mice but was confined to quantifications of cross-hemisphere gamma-band synchronization across a pair of fiber-optic probes, rather than capturing high-frequency voltage dynamics in unaveraged traces.^9^ To surpass this past work, we examined our prior TEMPO instrumentation and datasets with the goal of first identifying and then reducing the remnant noise sources.

We found that optical mode-hopping in the fiber conveying laser light to the brain was the main limiting noise source in our earlier studies and arose selectively during unconstrained animal locomotion and thus motion of the fiber (**Figure S1A–E**). Mode-hopping is a type of speckle noise occurring when coherent laser light propagates in multimode fiber, due to fluctuating interference between light in different modes when the fiber flexes.^48^ To prevent speckle, one might consider incoherent light-emitting diodes (LEDs) as alternative light sources. But solid-state lasers are generally more stable than LEDs and can be modulated at higher frequencies (∼50–100 kHz in this paper) at which detectors usually have much lower noise than at lower frequencies (*e.g.* ∼5 kHz for LEDs) (**Figure S1C**). By implication, even when LEDs are used for photometry with detectors of near-zero electronic noise, such as a photon-counting photomultiplier tube^49^ or a scientific-grade CMOS (sCMOS) camera^50–52^, the additional illumination noise (**Figures S1A,B**) prevents the system from attaining the low noise floor achievable in principle with laser illumination.

These considerations yielded a key engineering principle for building fiber-photometry systems for freely moving animals with greater sensitivity than prior LED- or laser-based versions. Namely, for illumination sources one should use high-frequency-modulated lasers for their superior stability, but this choice necessitates a means of breaking the illumination’s coherence for the sake of avoiding fiber-optic mode-hopping noise when the animal is actively behaving. By embodying this design principle, uSMAART surpasses the sensitivity of prior LED-based fiber photometry systems^9,50^ and of laser-based systems that lacked active suppression of speckle noise.^7,53^ In total, uSMAART has 4 modules, all designed to minimize noise: (i) a low-noise laser illumination module, enabling; (ii) a decoherence module to preclude mode-hopping noise; (iii) a high-efficiency fluorescence sensing module; (iv) a phase-sensitive detection module with digital lock-in amplifiers (**Figure 1A; STAR Methods**). The laser illumination module is about 10-times more stable than that of prior photometry systems (**Figure S1E**);^7,9,54,55^ when it is used together with the decoherence module, the r.m.s. laser illumination fluctuations at the specimen of only ∼0.004% are impervious to fiber and animal motion (**Figure S1D,E**). The fluorescence sensing module minimizes bleedthrough of emissions and photon shot noise from the reference fluor into the detection channel for the GEVI, and the phase-sensitive detection module unmixes emission signals that are excited by different lasers but captured on the same photodetector.

**Figure 1:**
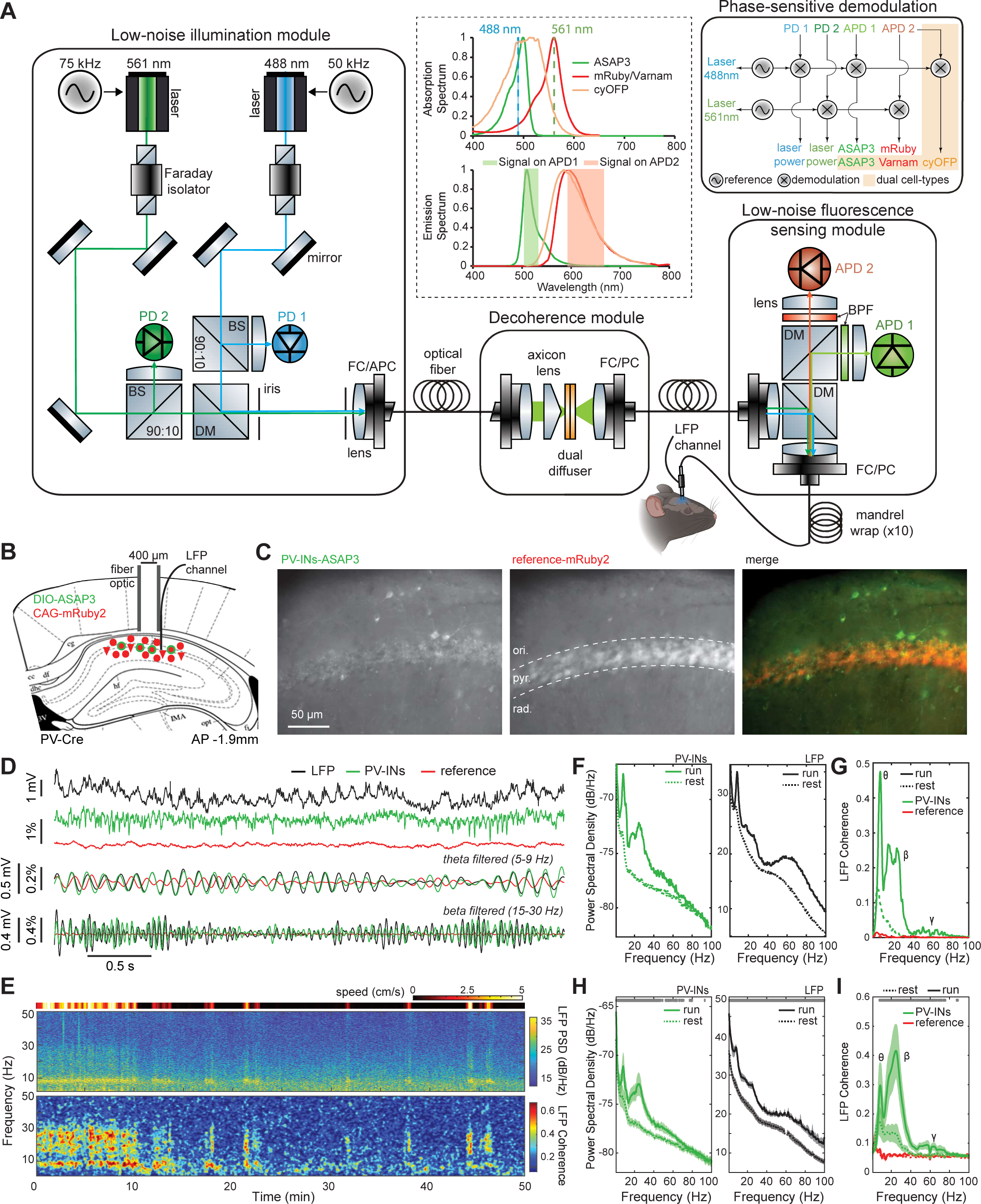
uSMAART fiber photometry captures population-level voltage dynamics, at frequencies up to 100 Hz, of genetically targeted neuron-types in freely behaving mice. (**A**) Schematic of the 4 modules in uSMAART. Within a low-noise illumination module (left box), the emissions of blue and green lasers are modulated at distinct frequencies (50 and 75 kHz, respectively) and jointly coupled into an optical fiber via an angled physical contact fiber connector (FC/APC). A portion of the light from each laser is split from the main pathway using a 90:10 beam splitter (BS) and monitored with a photodiode (PD). To achieve immunity to fiber motion artifacts, the illumination passes through a free-space, dual-stage optical diffuser within a decoherence module (middle box) and is then coupled back into optical fiber via a physical contact fiber connector (FC/PC). A low-noise fluorescence sensing module (lower right box) achieves high-efficiency, low-noise fluorescence detection using a pair of avalanche photodiodes (APDs) and a combination of dichroic mirrors (DMs) and bandpass filters (BPFs) that minimizes crosstalk between fluorescence channels. To attenuate noise from spatial mode-hopping, the fiber conveying light to and from the brain is wrapped ten times around a mandrel. Local field potential (LFP) signals are measured in the brain near the tip of the optical fiber. Electronics for phase-sensitive frequency demodulation (upper right box) isolate signals conveying the illumination and fluorescence emission powers. In the dual cell-type voltage-sensing configuration (*orange shading*), cyOFP emissions are captured on APD2 but demodulated at 50 kHz owing to their excitation by the 488-nm laser. *Inset* (dashed box): Absorption and emission fluorescence spectra for single and dual cell-type voltage-sensing configurations, for which an mRuby and cyOFP, respectively, are the reference fluors. (**B**) To track the aggregate transmembrane voltage activity of CA1 hippocampal parvalbumin-positive (PV) neurons (**C–I**), we virally expressed the ASAP3 voltage-indicator in PV-Cre mice, along with a reference fluor, mRuby2. As schematized in a coronal brain section, we implanted an optical fiber (400-μm diameter core) and an LFP electrode just atop CA1, to capture optical and electrical signals, respectively. AP: antero-posterior coordinate of the section relative to bregma. (**C**) Coronal hippocampal section from a PV-Cre mouse, imaged by fluorescence microscopy, showing expression of green fluorescent ASAP3 in PV interneurons (PV-INs; *left*), strong expression of red-fluorescent mRuby2 in the *stratum pyramidale* (pyr.) layer of CA1 and weaker expression in *stratum oriens* (ori.) and *stratum radiatum* (rad.) (*middle*), and the overlap of the two expression patterns (*right*). Scale bar: 50 µm. (**D**) Example traces showing concurrently acquired, LFP (*black trace*) and fluorescence signals from ASAP3 (*green trace*) and mRuby2 (*red trace*) in a freely behaving PV-Cre mouse, as well as overlaid, bandpass-filtered versions of the three traces in the theta (5-9 Hz) and beta (15–30 Hz) frequency ranges, revealing a strong concordance between the optical and electrical recordings. (The LFP is an extracellular measurement and thus is anti-correlated with the optical measurement of transmembrane voltage. Owing to the negative voltage-sensitivity of the indicators used, scale bars for all fluorescence indicator traces elsewhere in the paper are shown with negative units so that upward-sloped traces convey membrane depolarization. Whereas, here the scale bars for ASAP3 signals have positive units, for the sake of showing the phase alignment between oscillations in the LFP and ASAP3 recordings). (**E**) Color plots of locomotor speed (*top*), a time-dependent spectrogram of the LFP (*middle*), and the frequency-dependent coherence between the LFP and ASAP3 signals (*bottom*) across a 50-min continuous recording from the same mouse as in (**D**). Note the bouts of high theta- and beta-band coherence specifically when the mouse is running. (**F**) Power spectral density plots for the LFP (*right*) and ASAP3 fluorescence signals (*left*) determined from joint, continuous 50-min recordings from the same mouse as in (**D**), computed separately for periods when the mouse was running (solid curve) or resting (dashed curve). LFP and ASAP3 signals exhibit substantial power in the theta (5-9 Hz) and beta (15-30 Hz) frequency ranges during running bouts but not during rest. (**G**) Plots of the frequency-dependent coherence between LFP signals and either the ASAP3 fluorescence trace (green curves), or the mRuby2 trace (red curves), for time periods when the mouse of (**D**) was running (solid curves) or resting (dashed curves). Note the theta, beta and gamma activity specific to periods of locomotion. (**H**) Same as (**F**) but averaged across n=6 mice. Red dots near the top of the plots in **H** and **I** mark frequencies at which the *y*-axis values during running bouts differed significantly from those during resting periods (signed-rank tests; p<0.05; n=6 mice). (**I**) Same as (**G**) but averaged across n=6 mice. Coherence in the theta, beta and gamma frequency bands significantly increased during locomotion.

To quantitatively evaluate uSMAART against prior LED- or laser-based photometry apparatus that do not follow the uSMAART design principle,^7,9^ we generated artificial 50 Hz signal bursts using a movable fluorescent sample (**Figure S1F**) and verified uSMAART boosts sensitivity about ten-fold (**Figure S1G–I**). Further, for uSMAART studies in the live brain, we took LFP recordings at the same tissue sites to support our interpretations of the optical traces. We stipulated that, as in our prior studies of low-frequency rhythms,^7,9^ all high-frequency optical voltage events should be accompanied by a coherent rise in LFP signals in the same frequency band, not just by a concomitant rise in high-frequency LFP power over the event time-course^7,9^ (**Figure S1J–N**). This criterion might, in principle, lead one to overlook optical voltage signals from certain forms of neural activity that are not coherent with the LFP (**Discussion**), but ensuring coherence between the two modalities affords additional confidence in the tiny optical signals arising from high-frequency events. We applied this coherence criterion to every biological phenomenon studied in this paper but not every single recording, for in some mice we omitted LFP recordings for the sake of experimental simplicity.

In addition to unprecedented detection sensitivity and immunity to fiber motion, uSMAART allows concurrent recordings from two cell-types. This required an innovation beyond our prior TEMPO studies of one neuron-class, which combined a green GEVI to track voltage signals and a red reference fluor to track hemodynamic and other artifacts. For studies with two GEVIs, we tried using an infrared reference fluor, but, due to the wavelength-dependence of hemoglobin absorption and light scattering,^56^ an infrared fluor poorly tracked the artifacts affecting visible fluorescence. Instead, we combined spectrally distinct green and red GEVIs along with a long Stokes-shift fluorescent protein, cyOFP,^57^ of which the absorption spectrum overlaps that of the green GEVI and the emission spectrum overlaps that of the red GEVI (**Figure 1A**). This labeling approach, when applied along with two light sources modulated at distinct frequencies, enabled unambiguous detection of signals from the 3 fluors using only 2 fluorescence detection channels.

Our *in vivo* studies explored uSMAART’s ability to capture high-frequency neural dynamics, starting with densely labeled neurons of a single class in head-fixed mice (**Figure S2**) and then progressing to sparsely labeled cells in freely moving mice (**Figures 1,2**). We followed a similar progression for studies of two cell-types (**Figures S2, S3** and **3**). Throughout, we used various GEVIs, including the FRET-opsins Ace-mNeon1,^2^ Varnam1^8^ and Varnam2,^1^ and the GFP-based GEVI ASAP3.^58^ Given our interests in sub-threshold oscillations, ASAP3 proved superior due to its ∼10-fold greater sensitivity at subthreshold voltages (1% Δ*F*/*F* for 1 mV changes between –100 and –40 mV; **Figure S1O**). Altogether, using uSMAART and ASAP3, which each provided a ∼10-fold sensitivity gain (see **Figures S1I** and **S1O**, respectively), we achieved ∼100-fold greater sensitivity than prior TEMPO measurements.^7,9^ This ∼100-fold improvement was vital to observing the high-frequency neural phenomena described below.

**Figure 2:**
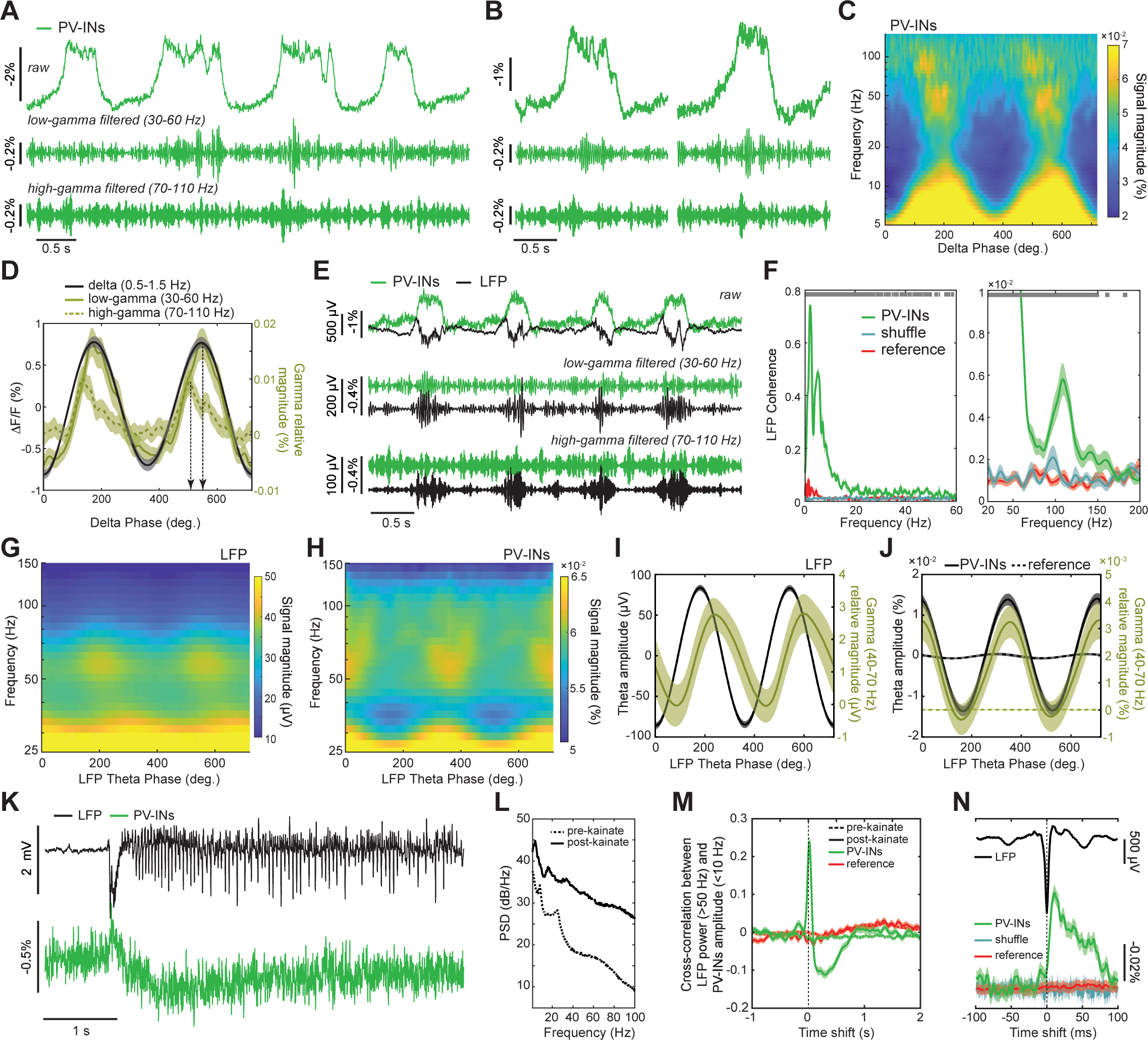
uSMAART captures cell-type specific cross-frequency coupling in anesthetized and freely behaving mice. (**A–F**) Delta-gamma frequency coupling in the membrane potential of ASAP3-labeled PV interneurons in the primary visual cortex (V1) of a ketamine-xylazine anesthetized mouse. (**A**) *Upper*: Example trace of PV cell voltage dynamics showing up- and down-state transitions. Filter versions of the trace illustrate that power in the low- (30-60 Hz; *Middle*) and high-gamma (70-–100 Hz; *Bottom*) frequency bands both increased during up-states. (**B**) Two example epochs of up- and down-state transitions, with raw and gamma filtered traces shown as in (**A**). (**C**) A wavelet spectrogram, with a logarithmic y-axis, averaged over 257 oscillation cycles in the delta frequency range (centered at 0.9 Hz, the frequency of peak delta power), reveals two distinct high-frequency power spectral peaks that arise at different phases of the delta rhythm. In panels **C, D,** the convention used for the phase of the delta oscillation is that 0-deg. refers to the trough of the oscillation, *i.e.*, the greatest hyperpolarization in the TEMPO signal. (**D**) Plot of the mean fluorescence signal in the delta frequency band (0.9 ± 0.25 Hz) (black trace; left axis), overlaid with plots of the delta phase-dependent oscillations of signal magnitude in the low (30–60 Hz; olive trace) and high (70–110 Hz; dashed olive trace) gamma ranges (right axis). As in (**A**, **B**), the plot shows distinct delta-phase shifts (dashed vertical lines; n = 122 delta events) for amplitude modulations of the low- and high-frequency gamma rhythms. The two arrows indicate the two delta phases at which the magnitudes of the low and high gamma oscillations are the greatest. Shading: s.e.m. (**E**) Raw (*top*), low-gamma (30–60 Hz;*middle*) and high-gamma (70–110 Hz;*bottom*) filtered traces acquired in joint LFP (black) and PV-cell TEMPO (green) recordings in cortical area V1 of a ketamine-xylazine anesthetized mouse. (**F**) Plots of frequency-dependent coherence for the same LFP and TEMPO recordings as in (**E**), using either a 2-s (*left*) or a 200-ms (*right*) temporal window to compute coherence values. The green, teal, and red curves respectively show coherence values between the LFP trace and traces for the ASAP3-labeled PV cells, a temporally shuffled version of the ASAP3 trace, and the red reference fluor. Gray dots at the top of the plots mark frequencies at which the coherence of the ASAP3 and LFP recordings differed significantly from the coherence of the LFP and the reference fluor (rank sum test; p<0.05), highlighting significant coherence in the delta, low- and high-gamma bands, as in (**A–D**). Shading: s.e.m. (**G–J**) Theta-gamma frequency coupling in the voltage dynamics of PV interneurons in the dorsal CA1 hippocampal area of a freely behaving mouse (see **Figure 1B** for labeling strategy). (**G, H**) Mean wavelet spectrograms for the LFP (**G**) and PV population voltage (**H**) recordings, plotted as a function of theta-phase over two theta cycles (with theta-phase extracted from the LFP trace at the frequency (7.5 Hz) of peak theta power). In **G–J**, the convention used for the phase is that 0-deg. refers to the trough of the theta oscillation in the LFP, *i.e.*, the greatest hyperpolarization in the extracellular electric field recording. (**I, J**) Mean signal amplitudes at the peak theta frequency (black traces; left axes) and signal magnitudes in the gamma (40–70 Hz) frequency range (olive traces; right axes) for LFP (**I**) and fluorescence (**J**) signals. Solid and dashed curves in **J** are for ASAP3 and reference fluor recordings, respectively. We made similar findings as in **G–J** in n = 4 mice. Shading: 95% C.I.. (**K–N**) PV cell-population voltage signals, as reported by ASAP3, revealed a ∼100 ms transmembrane depolarization, followed by a prolonged hyperpolarization, during a kainate-induced epileptic seizure (typified by high-frequency LFP dynamics) in the dorsal CA1 hippocampal area of a freely behaving mouse. (**K**) Example traces of LFP (black trace; top) and PV membrane voltage (green trace; bottom) signals, illustrating that the appearance of epileptic spikes in the LFP channel correlated with a ∼100 ms depolarization of PV interneurons (see also **N**). (**L**) Power spectral density plots for the LFP, recorded either during pre- (dashed curve) or after (solid curve) kainate injection. (**M**) Plots of the mean temporal cross-correlation between high-frequency (>50 Hz) power in the LFP and the amplitude (at frequencies <10 Hz) of PV-cell membrane voltage signals (green traces) or reference fluor signals (red traces). Solid traces: Averages over n=21 seizure events (each sampled for 5 s; **STAR Methods**) that occurred after kainate injection. Dashed curves: Averages over 5-s-intervals taken from the period before kainate injection. Shading in panels **M, N**: 95% C.I. (**N**) Plots of the epileptic, ictal spike-triggered average activity in the LFP and PV cell traces, and in the fluorescence reference channel and temporally shuffled versions of the PV cell traces. Note that PV interneurons depolarized during ictal spikes.

### Fiber-optic TEMPO captures high-frequency voltage dynamics in behaving animals

We first examined if uSMAART could sense the high-frequency dynamics of primary visual cortical (V1) neurons of awake head-fixed mice during passive viewing (**Figure S2A–F**). We virally expressed Varnam1 in pyramidal cells and the reference fluor GFP non-selectively (**Figure S2A**), and we recorded TEMPO and LFP signals while presenting drifting grating visual stimuli (**Figure S2B**). In accord with past work,^59–61^ visually evoked 3–7 Hz oscillations arose in both recordings (**Figure S2C–E**). Crucially, during visual stimulation the unmixed GEVI but not the reference fluor channel underwent a rise in coherence with the LFP over frequencies up to 30 Hz (**Figure S2F**), validating that TEMPO had captured cell-type-specific high-frequency dynamics.

Next, we sought to record high-frequency voltage signals from a sparse cell-type in head-fixed but active mice (**Figure S2G–L**). In hippocampal area CA1, we virally expressed ASAP3 in PV interneurons and the reference fluor mRuby2 non-selectively (**Figure S2G**). To elicit rest-to-run state changes, we delivered an airpuff to the mouse’s back as it rested (**Figure S2H**). The airpuff triggered locomotor-evoked ASAP3 and LFP signals in the beta frequency range (15–30 Hz). As in our prior work,^7,8^ reference fluor signals reflected a slow (∼1 Hz) hemodynamic response and the heartbeat (∼10-12 Hz) (**Figure S2I–K**), the absence of which in the ASAP3 traces showed the success of our unmixing strategy. A locomotor-triggered rise in beta-band coherence between the LFP and ASAP3 signals but not with mRuby2 signals confirmed the veracity of the PV cells’ beta-frequency dynamics (**Figure S2L**).

We next studied CA1 PV cells in freely behaving mice and tracked TEMPO and LFP signals up to ∼100 Hz as mice explored an open arena (**Figure 1B–I**). Like the LFP, TEMPO signals from PV neurons exhibited a clear dependence on the mouse’s locomotor state (**Figure 1E,F**). During locomotion, these TEMPO signals increased their power (**Figure 1H**) and coherence with the LFP (**Figure 1I**) in the theta (5–9 Hz), beta (15–30 Hz) and gamma (30-100 Hz) frequency bands (p<0.05; signed-rank tests; n=6 mice), whereas reference fluorescence signals did not.

### Cross-frequency coupling between low- and high-frequency voltage oscillations

With our newfound ability to capture cell-type specific high-frequency voltage dynamics, we decided to explore whether cross-frequency coupling (CFC), which is associated with a variety of human cognitive processes, could be observed in an individual neuron class. *A priori*, it was not clear this would be feasible, as CFC might be a collective electrophysiological phenomenon that emerges from the non-stationary contributions of diverse cell-types (**Figure 2**).^62–65^

We first studied cortical delta-gamma coupling in anesthetized mice (**Figure 2A–F**), previously characterized with electrical recordings.^66^ We recorded TEMPO signals in visual cortical PV cells expressing ASAP3 and found prominent delta rhythms,^7^ along with high-frequency gamma oscillations during up-states^66^ (**Figure 2A,B**). Strikingly, these up-state specific bursts of gamma activity were concentrated in two distinct spectral bands, low (30–60 Hz) and high (70–110 Hz) gamma bands, which were modulated at two different phases of delta (**Figure 2C,D**). The apex of high gamma activity preceded the peak delta depolarization (–56 ± 14 ms; mean ± s.e.m; p<10^-4^; n=122 delta oscillations; signed rank test), whereas the apex of low gamma followed it (17 ± 6 ms; p <0.005). Both gamma oscillations nested in the delta rhythm were coherent with the LFP at levels well above chance expectations or those for the reference channel (**Figure 2E,F**).

We next looked for CFC in freely behaving mice. In mice expressing ASAP3 in hippocampal PV cells, both LFP and TEMPO signals displayed theta-gamma CFC. Notably, the amplitude of PV cell gamma activity peaked at the apex of the LFP theta oscillation, whereas the peak amplitude of LFP gamma activity was phase-shifted relative to the LFP theta oscillation (**Figure 2G–J**); this fits with reports that PV cells exert a hyperpolarizing influence at the peak of the theta rhythm.^67^

### Bi-phasic dynamics of PV interneurons in an epilepsy model

We next explored neural voltage dynamics in a disease model. Dysfunctional PV cells are implicated in several neuropsychiatric disorders, including epilepsy,^68,69^ and activity imbalances between excitatory and inhibitory cells can induce hyperexcitability and seizures.^70^ Notably, alterations in the membrane voltage dynamics of PV cells can impact their ability to regulate pyramidal cell activity and maintain a balance between excitation and inhibition. To study PV cell voltage dynamics during drug-induced seizures, we utilized uSMAART in mice expressing ASAP3 in PV cells in area CA1 of the dorsal hippocampus.

We induced seizures with kainic acid and jointly recorded LFP and TEMPO signals in freely moving mice (**Figure 2K,L**). During epileptiform events, PV cell voltage dynamics exhibited two successive characteristic changes. First, they depolarized for ∼100 ms, during which the LFP revealed an extracellular hyperpolarization (*i.e.* depolarized intracellular voltages inducing a local extracellular current sink) (**Figure 2M,N**). Then, PV cells hyperpolarized for ∼1 s as the tissue became hyperexcitable and exhibited ictal spikes, brief high-frequency bursts of activity thought to result from a positive feedback loop between excitatory and inhibitory cells.^71^ During this second phase of seizure activity, another interneuron-type might limit the PV cell discharge initiated in the first phase. The ictal spikes observed here in PV cells’ dynamics fit with prior recordings^72^, support the idea that there is a shift between excitatory and inhibitory activity during epileptiform events,^73^ and show that TEMPO can help reveal how different neuron-types influence epilepsy generation.

### Dual neuron-class recordings of visually evoked activity in awake mice

To validate its capacity for dual cell-type recordings, we assessed if uSMAART can report the concurrent dynamics of glutamatergic and GABAergic neural populations. We first studied awake head-fixed mice expressing cyOFP non-selectively, and Ace-mNeon1 and Varnam1 in pyramidal cells and interneurons, respectively, during passive viewing of visual stimuli (**Figure S2M–P**). In line with our studies of individual cell-types (**Figure S2**), these older GEVIs reported stimulus-evoked oscillations in both cell-types in the 3-7 Hz frequency band but not gamma frequencies (**Figure S3O,P**). To capture excitatory/inhibitory interactions in more detail, we prepared mice expressing ASAP3 in PV cells and Varnam2 in pyramidal cells (**Figure 3A**). Visual stimuli consistently evoked post-stimulus 3-7 Hz oscillations in PV and pyramidal cells (**Figure 3B–E**). The superior dynamic range of ASAP3 also enabled the detection of stimulus-evoked increases in gamma (30–70 Hz) activity in PV interneurons (**Figure 3F**). To verify that the lack of observed gamma activation in pyramidal cells was due to the indicator assignments, rather than a *bona fide* difference between the two cell-types, we prepared mice in which we reversed the indicator assignments between pyramidal and PV cells and performed joint LFP and TEMPO recordings (**Figure 3G-I**). In these mice, both cell-types and the LFP exhibited the post-stimulus 3-7 Hz band activation (**Figure 3H**), and pyramidal cells showed the expected increases in gamma activity during visual stimulation, as did the LFP signals (**Figure 3I**).

**Figure 3:**
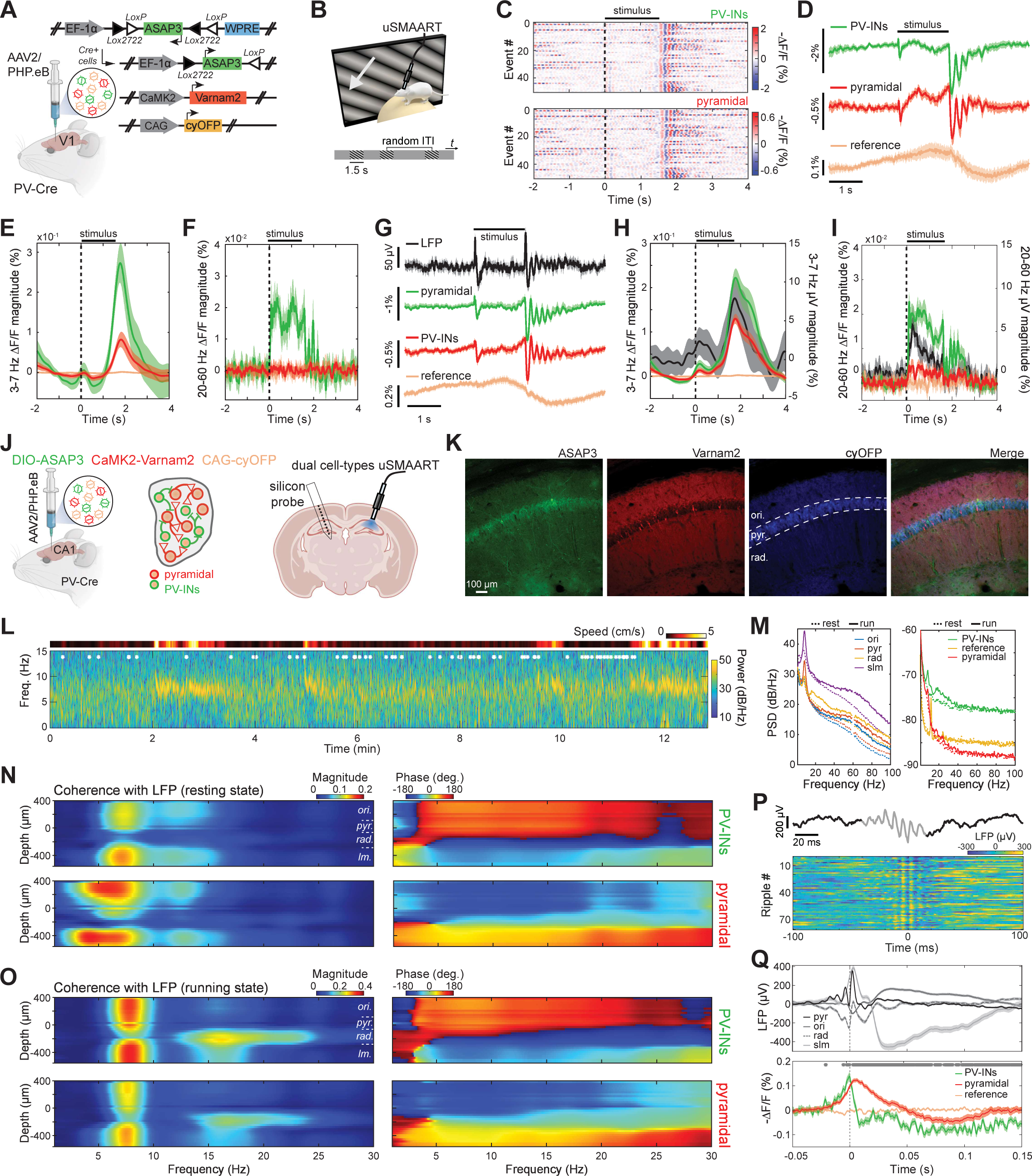
uSMAART tracks the concurrent voltage dynamics of two genetically identified neuron classes in behaving mice. (**A–I**) Visual stimulation evoked gamma oscillations in brain area V1 of awake mice during stimulus presentation, followed by 3-7 Hz oscillations after stimulus offset. (**A**) Retro-orbital injection of three PHP.eB adeno-associated viruses into PV-Cre mice enabled expression of Cre-dependent ASAP3 in PV interneurons, Varnam2 in pyramidal cells, and cyOFP (a reference fluorophore) in all neuron-types. (**B**) Schematic of the visual stimulation paradigm used for dual cell-type voltage-sensing studies. Head-fixed mice viewed a computer monitor placed in front of one eye, as we recorded fluorescence voltage activity in the contralateral primary visual cortex (V1). Drifting grating visual stimuli (each 1.5 s in duration) swept across the monitor. Intervals between stimulation trials were randomized between 2–5 s. (**C**) Visual stimuli consistently evoked post-stimulus 3-7 Hz oscillations in ASAP3-labeled PV interneurons (top) and Varnam2-labeled pyramidal (bottom) cells. Each row has data from one of 50 different trials in the same mouse. (**D**) Mean time-dependent fluorescence traces, obtained by averaging the signals from all 3 fluors across all 50 trials of (**C**). Shading in (**D**–**I**): 95% C.I. (**E, F**) Mean time-dependent fluorescence signal magnitudes in the 3-7 Hz (**E**) and gamma (30–70 Hz; **F**) frequency bands for all 3 fluors used in (**C, D**), computed using wavelet transforms. (**G**) Mean time-dependent fluorescence traces from studies in which we reversed the GEVI assignments to PV and pyramidal cells from those of (**C**), and in which we concurrently performed LFP and TEMPO recordings (averages are over 100 trials). (**H, I**) Same as (**E, F**) but for the studies of **G**, including the LFP recordings. (**J–Q**) We studied the dynamical inter-relationships between excitatory and inhibitory activity in the hippocampus of an active mouse, across different behavioral states. (**J**) We performed dual cell-type fluorescence recordings with uSMAART, concurrently with electrophysiological recordings in the contralateral CA1 area (linear silicon probe; 32 recording sites) that allowed us to detect ripple (120–200 Hz) events. The fluorescence labeling strategy using 3 different viruses was the same as in panel (**A**). (**K**) Fluorescence confocal images of a brain slice of a mouse expressing ASAP3 in PV interneurons, Varnam2 in pyramidal cells, and cyOFP in all cell-types. The far right image shows an overlay of the three preceding images, highlighting the different targeting of all three fluorescent proteins. (**L**) *Top*: Plot of mouse speed. *Bottom*: Time-dependent spectrogram of LFP signals recorded in CA1 *stratum pyramidale*. White asterisks mark occurrences of ripple events. The LFP exhibited power increases in the theta (5–9 Hz) band during locomotion, whereas ripples occurred only during rest, consistent with past studies. (**M**) Power spectral densities for (*left*) LFP signals recorded across all 4 canonical hippocampal layers (ori.: *stratum oriens*, pyr.: *stratum pyramidale*, rad.: *stratum radiatum*, slm.: *stratum lacunosum moleculare*) and for (*right*) the three fluorescent signals from the TEMPO recordings, for periods of rest (dashed curves) or running (solid curves). (**N, O**) Plots of the coherence magnitude (*left plots*) and phase (*right plots*) between PV (*top rows*) and pyramidal cell (*bottom rows*) fluorescence voltage signals and the LFP measured across the 32 recording sites of the silicon probe (plotted as a function of depth relative to the center of *stratum pyramidale*; tissue layers abbreviated as in **M**) during periods of resting (**N**) or running (**O**). During rest, PV and pyramidal cell voltage dynamics were both coherent with the LFP but with distinct frequency signatures, extending up to the alpha frequency range (∼10–15 Hz). During locomotion, both cell-types became highly coherent with the LFP at theta frequencies (∼5–9 Hz) in nearly all layers of CA1; however, the LFP signal in *stratum radiatum* (depth: ∼200 μm) had higher coherence with each cell-type in the beta band (∼15–30 Hz) than at theta frequencies. Note the sharp changes in coherence phase that occur near the boundaries of different tissue layers. (**P**) *Top*: Example LFP trace showing a hippocampal ripple event (light gray part of the trace) recorded in *stratum pyramidale*. *Bottom*: LFP traces shown for 80 different ripple events from the same mouse, temporally aligned to the time on each trial at which the LFP signal was at its minimum. (Q) Mean time-dependent traces of the LFP signals across all 4 hippocampal layers (*top*) and fluorescence voltage signals for PV and pyramidal cells (*bottom*) during ripples, obtained by averaging over 273 ripples. At ripple peak magnitude (vertical dashed line), both pyramidal cells and PV depolarized. PV cells then hyperpolarized more sharply, whereas pyramidal cell voltages more gradually hyperpolarized below baseline voltages. Gray dots at the top of the bottom plot mark times at which the fluorescence changes in the two neuron classes were significantly different (two-tailed signed-rank test; p<0.05). Shaded areas: s.e.m.

### Dual neuron-class recordings in the hippocampus of freely moving mice

We next studied the relationships between excitatory and inhibitory activity in the hippocampus by performing dual cell-type uSMAART recordings in mice expressing cyOFP non-selectively, ASAP3 in PV cells and Varnam2 in pyramidal cells (**Figure 3J–Q**). We also recorded electrical dynamics in the contralateral hippocampus with a linear silicon probe, which allowed us to identify well-known physiological signatures of the different hippocampal laminae (**Figure S3A,B**).^74–76^ As in prior studies, LFP power in theta (5-9 Hz) frequencies rose across the laminae during locomotion, but hippocampal sharp wave ripples (120–200 Hz) occurred mainly during rest (**Figure 3L,M**). Locomotor-evoked increases in beta (15–30 Hz) activity were more limited in the LFP recordings to *stratum radiatum*, whereas the evoked increases in gamma activity (30–100 Hz) occurred across laminae (**Figure 3M**). Especially in ASAP3-labeled PV cells, locomotion triggered prominent increases in theta and beta activity (**Figure 3M**), consistent with our prior results (**Figure 1**).

During rest, the dynamics of both PV and pyramidal cells were coherent with the LFP, but with distinct frequency signatures (**Figures 3N** and **S3C–E**). During running, both cell-types had theta-band (5-9 Hz) activity of increased coherence with the LFP in nearly all CA1 layers, as well as beta (15–30 Hz) and gamma (30-70 Hz) dynamics that were more selectively coherent with the LFP in *stratum radiatum* (**Figures 3O** and **S3C–E**).

During ripples (**Figure 3P**), both PV and pyramidal cells depolarized at ripple onset, consistent with reports that the peak of pyramidal cell activity aligns with the ripple’s envelope peak.^77,78^ This was followed by a sharp hyperpolarization of PV cells preceding the depolarization peak of the pyramidal cells, and then by a more gradual hyperpolarization of pyramidal cells (**Figures 3Q** and **S3F**). The steepness of the PV cell hyperpolarization is suggestive of a time-gating mechanism for precisely releasing synchronized pyramidal cell firing during SWRs, in accord with the idea that inhibitory control by PV interneurons shape the output of pyramidal cells during SWRs.^75,79^ Future studies should investigate the mechanisms underlying these interactions and explore their consequences for information encoding and memory consolidation. Altogether, these findings show that, when combined with our triple fluor expression strategy, uSMAART can reveal the joint dynamics of two neuron-classes in freely moving mice.

### Concurrent uSMAART recordings of two cell-types in two brain areas

An ability to track the concurrent dynamics of multiple identified cell classes across more than one brain area in freely behaving mice would empower mechanistic studies of inter-area oscillatory phenomena. uSMAART provides this capability, in that it allows joint recordings of two cell-types in each of two brain areas (**Figure S3G–M**). In ketamine-xylazine (KX) anesthetized mice expressing ASAP3 in PV cells and Varnam2 in glutamatergic neurons across cortex (**Figure S3H**), interneurons and pyramidal cells in area V1 and primary motor cortex (M1) displayed up-and-down state transitions in a delta rhythm with gamma oscillations nested into the up-state (**Figure S3I**). In line with our prior results (**Figure 2**), PV cells in V1 and M1 exhibited delta-gamma CFC (**Figure S3J–L**). We assessed the time delays between the delta oscillations in the two brain areas using the TEMPO signals from either the PV or pyramidal cells (**Figure S3M**). Using PV cells to estimate the delay, the delta rhythms in M1 had a temporal lead of –103 ± 41 ms (mean ± s.d.) ahead of those in V1. The same calculation for pyramidal cells led to an estimated temporal lead in M1 of –117 ± 35 ms. These results are suggestive of a voltage wave traveling at 10–15 mm⋅s^-1^ from anterior to posterior, which prompted us to do follow-up imaging studies (**Figure 4**).

**Figure 4:**
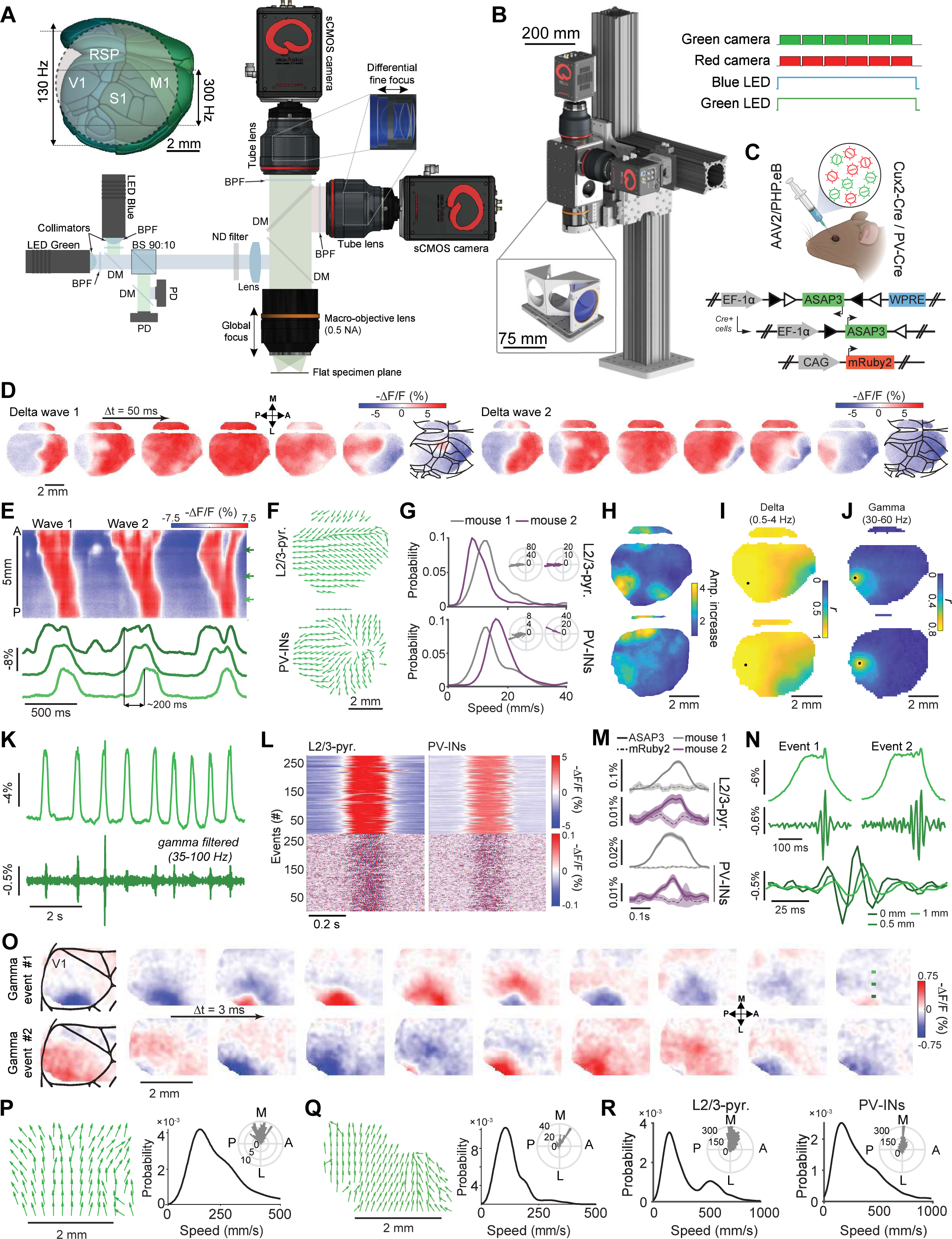
TEMPO imaging in anesthetized mice reveals traveling neocortical waves in the delta and gamma frequency bands that exhibit cross-frequency coupling. (**A**) Schematic of the TEMPO mesoscope. A pair of low-noise light-emitting diodes provides two-color illumination; a corresponding pair of photodiodes monitors their emission powers. The illumination reflects off a dual-band dichroic mirror (70 mm × 100 mm) and is focused onto the specimen by a 0.5 NA macro objective lens providing a 8-mm-diameter maximum field-of-view (FOV) when used in our system. Fluorescence returns through the objective lens and dual-band dichroic mirror. A short-pass dichroic mirror (70 mm × 100 mm) splits the fluorescence into two detection channels. After passing through a bandpass filter, fluorescence in each channel is focused onto a fast sCMOS camera by a tube lens (85 mm effective focal length). *Inset*: A view from the Allen Brain Atlas, showing the location of the glass cranial windows (dotted black circle; 7–8 mm diameter) used for **Figures 4,5**,**7**. Across the full area visible through the cranial window, the cameras acquired images at 130 Hz. In some studies, to increase the imaging speed to 300 Hz, we sampled a region-of-interest on the camera chip that covered the brain areas between the two black horizontal lines. *Abbreviations*: RSP (retrosplenial cortex), M1 (primary motor cortex), V1 (primary visual cortex), S1 (primary somatosensory cortex), BPF (bandpass filter), BS (beamsplitter), DM (dichroic mirror), LED (light-emitting diode), ND (neutral density), PD (photodiode). (**B**) Computer-assisted design mechanical drawing of the mesoscope. *Lower inset,* magnified view of the large custom fluorescence filter set. *Upper inset*: Timing protocol for voltage imaging using a single green GEVI plus the mRuby2 reference fluor. Illumination from both LEDs is continuous (bottom two traces). Image acquisition by the two cameras is initiated by an external trigger, ensuring that the image pairs are temporally aligned (top two traces; 800 Mbytes ⋅ s^-1^ data rate per camera). (**C**) *Top,* We retro-orbitally co-injected a pair of AAV2/PHP.eB viruses to co-express red fluorescent mRuby2 and a green fluorescent GEVI, ASAP3. *Bottom,* One virus expresses mRuby2 via the CAG promoter; the other allows Cre-dependent expression of ASAP3 via the EF-1α promoter. By using Cux2-Cre^ERT2^ or PV-Cre mice, we performed voltage-imaging studies of neocortical layer 2/3 pyramidal (L2/3) or PV cells, respectively. (**D**) Two examples of traveling voltage waves in the delta frequency band, shown in ASAP3 image sequences (50 ms between images) taken at 130 Hz in ketamine-xylazine-anesthetized Cux2-Cre^ERT2^ mice. Images underwent unmixing (**Figure S5**) to remove hemodynamic changes captured in the mRuby2 channel but were otherwise unfiltered. Brain area boundaries (see (**A**) inset), are superposed on the last image in each sequence. In each case, a depolarization (denoted by red hues) sweeps across cortex in the anterior to posterior (A-P) direction. Data in (**E–R**) are also from ketamine-xylazine-anesthetized mice and were acquired at 130 Hz in (**D–J**) and 300 Hz in (**K–R**). (**E**) Color plot (*top*) showing the anterior to posterior propagation of the two traveling waves in (**D**). At each time point (*x*-axis) and for each A-P coordinate (*y*-axis), we averaged fluorescence values along the medio-lateral direction. Arrows in 3 different shades of green mark 3 different positions along the A-P axis for which voltage-dependent fluorescence traces are plotted (*bottom*) in corresponding colors. (**F**) Flow maps showing local propagation directions of voltage depolarization for a pair of individual delta waves observed in example Cux2-Cre^ERT2^ (*top*) and PV-Cre (*bottom*) mice. Flow vectors are all normalized to have the same length. (**G**) Distributions of delta wave propagation speed across all delta events seen in two Cux2-Cre^ERT2^ (*top*) and two (PV-Cre) (*bottom*) mice, computed near the center of area V1 (marked by black dots in (**I**)), where there was consistent anterior to posterior propagation. *Insets*: Polar histograms showing distributions of wave propagation direction for the same 4 mice at the center of V1, revealing the approximate alignment of wave propagation with the A-P axis in all 4 mice (n = 313, 320, 405 and 200 delta events in the individual mice). (**H**) During the peaks of the delta waves, we found enhanced activity in the gamma (30–60 Hz) frequency band. In two example Cux2-Cre^ERT2^ (*top*) and PV-Cre mice (*bottom*), the amplitudes of gamma oscillations increased during delta wave depolarizations up to ∼4-fold over baseline values in brain areas V1 and RSP and to a lesser extent in other areas. (**I**) Maps of peak correlation coefficients, *r*, for the same mice as in (**H**), computed for each spatial point by calculating the temporal correlation function between the local fluorescence trace and that at the center of V1 (black dots) and then finding this function’s maximum value. (**J**) Maps of peak correlation coefficients, computed as in (**I**) but using gamma bandpass-filtered fluorescence traces, show that the gamma oscillations were less spatially coherent than the delta waves. (**K**) To study gamma activity in greater detail, we acquired a subset of the video data at 300 Hz (**K–R**) over a more limited FOV (see inset of (**A**)). *Top*: Example fluorescence trace of voltage activity at the center of V1 in the same Cux2-Cre^ERT2^ mouse as in (**H-J**), showing ongoing delta waves. *Bottom*: Gamma-band filtered (35–100 Hz) version of the top trace, revealing increases in gamma-band activity during the peaks of the delta oscillations. (**L**) *Top*: Example fluorescence traces of voltage activity at the center of V1, from each of the two mice in (**H-J**), temporally aligned to the peak of delta wave depolarization to reveal the consistent waveform of delta activity (n = 260 waves shown per mouse). *Bottom*: Gamma-band filtered (35–100 Hz) versions of the same traces reveal delta-gamma coupling as in (**K**). (**M**) Mean time-dependent amplitudes of gamma-band (35–100 Hz) activity in the voltage (solid lines) and reference (dashed lines) signals, for each of the 4 mice in (**G**), computed by averaging over all delta events in each mouse, identified as in (**L**) and aligned to the peak of the delta oscillation using the wavelet spectrogram. Voltage but not reference channel signals showed increased gamma band activity during the depolarization phase of delta oscillations. Shading: 95% C.I. (**N**) *Top*: Magnified views of two example delta depolarization events at the center of V1 in the same mouse as in (**K**). *Middle*: Gamma-filtered (35–100 Hz) versions of the same traces, highlighting gamma events near the end of each delta depolarization. *Bottom*: Gamma-filtered traces of activity during Event 1, from the color-corresponding locations marked with green-shaded dots in the upper rightmost image of panel (**O**). (**O**) Sequences of gamma-band (35-100 Hz) filtered images (3 ms between successive frames), showing the spatiotemporal dynamics in V1 of the same two delta depolarization events as in (**N**). Green dots mark the anatomic locations for the voltage traces in the bottom panel of (**N**). For display purposes only, the images shown were spatially low-pass filtered using a Gaussian filter (156 μm FWHM). (**P, Q**) *Left panels*: Flow maps showing the local wave propagation directions during the same two individual gamma wave events as in (**N**). As in (**F**), all flow vectors are normalized to have the same length. *Right panels*: Histograms showing the distributions of propagation speed across the brain region shown in (**O**), for the same two gamma events as in the left panels. Unlike delta waves, which traveled along the A-P axis, the gamma waves illustrated here traveled more aligned to the medio-lateral (M-L) axis and had much faster speeds than the accompanying delta waves (compare to (**G**)). *Insets*: Polar histograms showing the distributions of propagation direction for the two gamma events, computed across the flow maps of the left panels. (**R**) Histograms of propagation speed, aggregated across n = 20 events and all spatial bins (62.5 µm wide) in V1, in the same two mice as in (**H-J**) (n = 200 and 300 waves, respectively, in the Cux2-Cre^ERT2^ and PV-Cre mice). *Insets*: Polar histograms of the directions of gamma wave propagation, showing that in both L2/3 pyramidal and PV cells the gamma waves traveled approximately in the M-L direction, roughly orthogonal to the propagation directions of their carrier delta waves (**E–G**).

### A TEMPO mesoscope

As fiber photometry recordings are inherently local and limited to signals arising beneath the fiber tip, we created a complementary instrument, the TEMPO mesoscope (**Figures 4A,B** and **S4A–K**), to image the spatiotemporal dynamics of population voltage activity in identified cell-types across the cortical surface of head-restrained mice. The instrument design embodies the TEMPO approach and has 3 main features: (1) a pair of low-noise LEDs for dual-color fluorescence excitation of two GEVIs plus a reference fluor; (2) a low-aberration, high numerical aperture (NA) macro-objective lens to image a field-of-view (FOV) up to 8 mm wide (∼50 mm^2^); and (3) two fluorescence detection pathways with a pair of sCMOS cameras to track GEVI and reference fluor emissions concurrently. We acquired full-frame images on the sCMOS cameras at 130 Hz, above the Nyquist frequency for neural activity in the low-gamma band (30–60 Hz). When we sought to detect higher frequency activity or capture fine temporal snapshots of voltage wave propagation, our cameras allowed imaging at 300 Hz albeit over a narrower FOV (2.7 mm ✕ 8 mm) (**Figure 4A**, *inset*). In either imaging mode, in 10 min of operation the system generates about 1 TB of raw data, from which we unmix the voltage signals from biological and instrumentation artifacts.

### Frequency-dependent unmixing of voltage and artifact signals

In our current and prior studies,^7–9^ unmixing biological and instrumentation noise from the voltage channel is a key part of the TEMPO approach. We initially used independent component analysis to unmix artifacts captured in the reference channel from the voltage channel^7^ and have since explored the use of linear regression. However, these algorithms are suboptimal in that they lack the flexibility to account for the variable extents to which different artifacts impact the two channels in different frequency bands (**Figure S5A**). To boost the sensitivity with which TEMPO can detect small-amplitude neural oscillations, we sought an improved unmixing method.

The main biological artifacts reflect hemodynamic processes, which in general are spatially heterogeneous across the field-of-view.^56,80^ Heartbeat-related signals and changes in tissue’s optical properties due to variations in blood volume and hemoglobin oxygenation have distinct temporal frequency signatures and are generally coherent between the GEVI and reference fluor channels, but with non-zero phase shifts (**Figure S5B**). Prior methods to unmix hemodynamic contributions to fluorescence signals used either frequency-independent or zero-phase lag transformations that cannot capture frequency-dependent, non-zero phase lags.^7,8,13,56,81,82^

To address this deficiency, we created a convolutional filtering approach^83^ to remove biological and instrumentation artifacts from neural voltage signals in a frequency-dependent way (**Figure S5C–M, STAR Methods**). The algorithm first estimates a linear filter that describes the frequency-dependent manner in which artifacts in the reference channel affect the GEVI channel. It then convolves this filter with the reference channel trace to obtain an estimate of the non-voltage signals in the GEVI channel. Lastly, it subtracts this estimate from the GEVI channel trace to estimate the true voltage signals. This unmixing method outperformed a simple (*i.e.*, frequency independent) linear regression in that it better removed broad- and narrow-band artifacts and did not transfer noise from reference to GEVI channel (**Figure S5E–I**), which was crucial for detecting high-frequency activity with high sensitivity (**Figure S5J**). When applied to individual or groups of pixels in the TEMPO video data, our convolutional approach captured the heterogeneity of hemodynamic signals across different blood vessels (**Figure S5K,L**). Altogether, unlike prior methods, convolutional unmixing preserves high-frequency voltage signals, is applicable to both fiber-optic and imaging TEMPO data, and is an important facet of the toolbox and all analyses presented in this paper.

### A transgenic reporter mouse for expressing the TEMPO fluorophores

To co-express the FRET-opsin voltage sensor AcemNeon1^2^ and the reference fluor mRuby3^84^ in a chosen cell-type, we created a transgenic Cre-dependent reporter mouse line expressly designed for TEMPO studies (**Figure S6A–S**). We used our published Flp-in approach to make a TIGRE 2.0-based reporter mouse (**Figure S6A, STAR Methods**).^85–87^ By enabling dual fluor expression without virus injection, this mouse (termed Ai218) gives TEMPO users a convenient option for uniform indicator labeling in a way that is designed to be highly reproducible across mice.

To evaluate the Ai218 mouse line for TEMPO imaging, we crossed it with PV-Cre or Cux2-Cre^ERT2^ driver mice, to express Ace-mNeon1 and mRuby3 in PV or layer 2/3 (L2/3) pyramidal cells, respectively (**Figure S6B,C**). For optical access to the majority of a cortical hemisphere (∼20 brain areas), we created a preparation in which a circular glass window of 7–8 mm diameter is installed in the cranium, allowing us to repeatedly image individual mice over several months.

We first assessed the ability of the TEMPO mesoscope to track voltage rhythms in laminar tissues, where LFP recordings have long been fruitful^88,89^ and fiber-optic TEMPO was validated (**Figures 3, S2, S3**).^7,8^ In KX-anesthetized Cux2-Cre^ERT2^ × Ai218 mice (**Figure S6B**), we found prominent up-down state transitions traveling anterior to posterior (**Figure S6D–G**). These delta (0.5-4 Hz) waves had a high spatial coherence across most of the imaged brain areas (**Figure S6H**). But, unlike past electrical measurements^66^ and our uSMAART studies (**Figures 2, S3**), we did not see gamma range (30–100 Hz) dynamics (**Figure S6I–K**), very likely due to the 10-fold poorer subthreshold dynamic range of Ace-mNeon1 as compared to ASAP3 (**Figure S1O**).

Next, we studied cell-type specific voltage dynamics in awake PV-Cre × Ai218 mice (**Figure S6C**) as they viewed moving grating visual stimuli (**FIgure S6L**). Strong 3-7 Hz oscillations arose at visual stimulus offset in V1 but not other brain areas (**Figure S6M–R**). Visually evoked activity in the gamma range (20–50 Hz) arose in visual cortex during visual stimulation (**Figure S6Q,O,S**), in line with our uSMAART data (**Figures 3, S2**). Overall, the Ai218 reporter mice enabled explorations of cell-type specific voltage oscillations up to the low gamma range.

### Traveling cortical voltage waves exhibit delta-gamma cross-frequency coupling

While the Ai218 mouse offers a convenient means of expressing TEMPO fluors, we also sought to benefit from newer GEVIs.^1,58^ Thus, we created viral tools, based on the PHP.eB serotype^90^ of adeno-associated viruses (AAVs), for brain-wide expression of GEVI and reference fluors. For instance, to study cortical PV or L2/3 pyramidal cells, we retro-orbitally co-injected into PV-Cre or Cux2-Cre^ERT2^ mice, respectively, a pair of AAV2/PHP.eB viruses to drive Cre-dependent expression of ASAP3 and non-selective expression of mRuby2 (**Figure 4C, S6T,U**).

With this labeling method, we found delta wave depolarizations that swept across cortex in KX-anesthetized mice (**Figure 4D–H**). These waves traveled anterior-to-posterior (A-P) in excitatory and inhibitory cell-types (**Figure 4F,G**; n = 200–405 delta events in individual mice, speeds: 12.0 ± 0.4 mm/s for Cux2-Cre^ERT2^ and 15.7 ± 0.9 mm/s for PV-Cre mice; angle from the anterior direction: 176 ±1 deg for Cux2-Cre^ERT2^ and 164 ± 1 deg. for PV-Cre mice; mean ± s.e.m.). We also found localized increases in gamma (30–60 Hz) activity, which arose during the peaks of delta waves and, unlike delta waves, were confined to a ∼2 mm wide area (**Figure 4I,J**).

To study gamma activity in more detail, we took a subset of the video data at 300 Hz over a more limited field-of-view (**Figure 4K–R**). This revealed delta-gamma coupling in L2/3 pyramidal and PV cell-types, with gamma arising at the delta wave peaks (**Figure 4K–N**). We quantified the speed and direction of each individual delta-nested gamma event (**Figure 4O–R**). Unlike delta waves, which traveled approximately along the A-P axis, gamma waves traveled approximately along the medio-lateral (M-L) axis (72 ± 3 and 88 ± 5 deg. from the A-P axis in Cux2-Cre^ERT2^ and PV-Cre mice, respectively; mean ± s.e.m.) and were much faster than the accompanying delta waves (225 ± 28 and 279 ± 20 mm⋅s^-1^ in Cux2-Cre^ERT2^ and PV-Cre mice, respectively; **Figure 4R**). This is an important observation, because it shows that coupled oscillations need not co-propagate in the same anatomic direction. Overall, TEMPO imaging was crucial for uncovering the complex spatiotemporal patterns of the nested delta and gamma waves.

### Visually evoked sequences of traveling gamma and 3-7 Hz waves in awake mice

We next studied visual cortical processing in awake mice. Mice with virally-mediated expression of ASAP3 in PV or cortical L2/3 pyramidal cells viewed drifting grating stimuli with the eye contralateral to the cortical hemisphere in which we performed TEMPO imaging (**Figure 5**, **Figure S7A**). In these mice, gamma waves (30–60 Hz) consistently arose in the visual cortex during stimulus presentation, followed by waves of ∼3–7 Hz frequency at stimulus offset^59,61^ (**Figure 5A–L**). Here, both PV and L2/3 pyramidal cells showed significant rises in gamma power in area V1 during visual stimulation over baseline (p<10^-9^ for each of 2 PV-Cre and p<10^-3^ for each of 3 Cux2-Cre^ERT2^ mice, n=50 trials each, rank sum test, **Figure 5H,L**). By comparison, reference fluor signals were stationary (p>0.27 in all mice). Visually evoked 3–7 Hz rhythms were previously found in rodents in electrophysiological studies and likely involve thalamocortical interactions akin to those of the human alpha rhythm.^59–61^ We also observed 3-7 Hz oscillations in areas other than V1, such as in M1, but these were incoherent with the visually evoked waves in V1 (**Figure 5I–K**).

**Figure 5:**
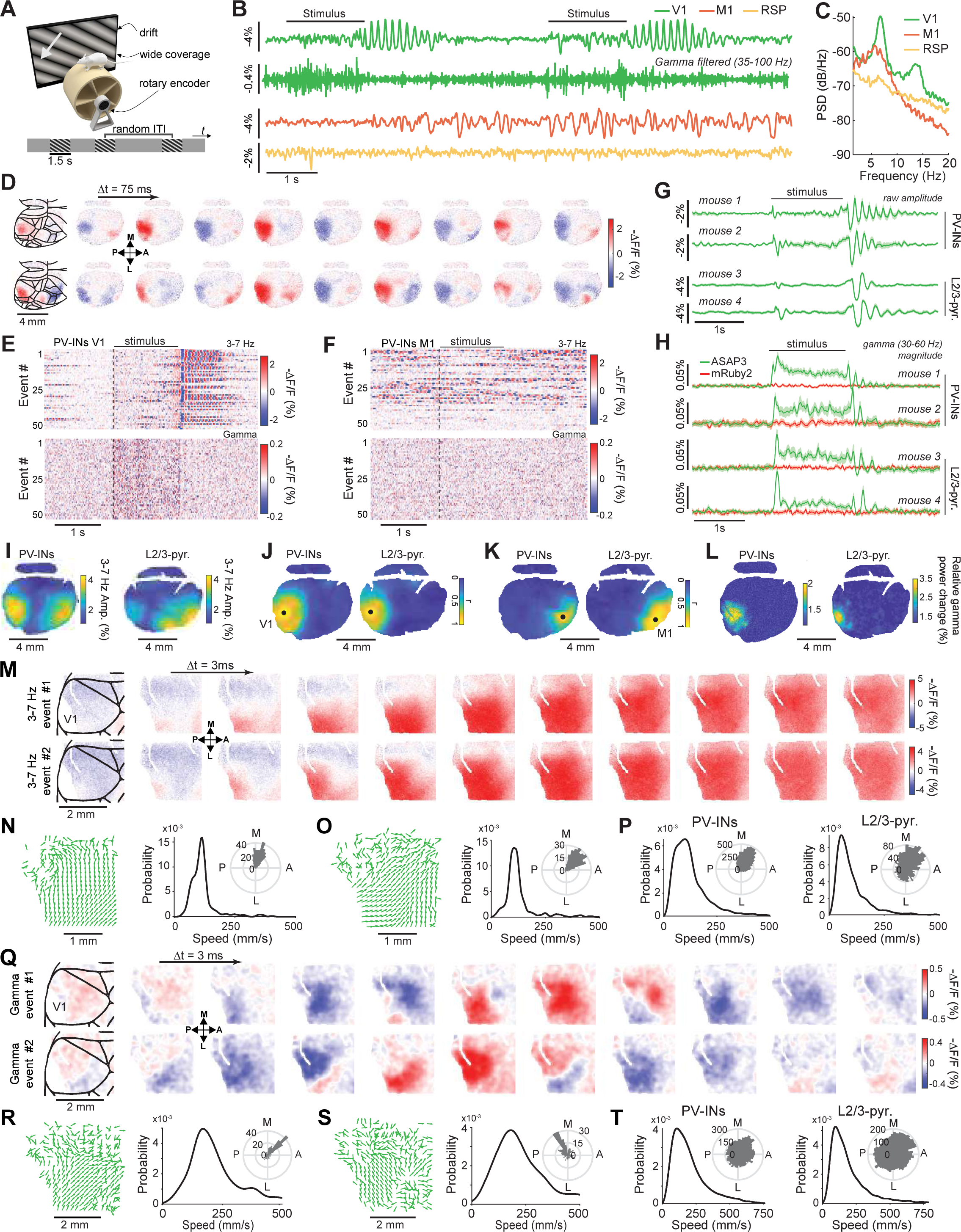
TEMPO imaging reveals visually evoked sequences of traveling gamma and then 3-7 Hz waves in area V1 of awake head-fixed mice. (**A**) For all studies in this figure, we used the same visual stimulation approach as in **Figure 3B**. We expressed the GEVI (ASAP3) and reference fluor (mRuby2) as in **Figure 4C**. (**B**) Example traces of visual stimulus-evoked voltage activity from a PV-Cre mouse, averaged over primary visual cortex (V1), primary motor cortex (M1) and retrosplenial cortex (RSP). In V1, visual stimuli evoked gamma oscillations (as seen in the gamma-filtered version of the raw trace) and 3-7 Hz oscillations after stimulus offset. (**C**) Power spectral densities (PSDs) of voltage activity determined from fluorescence traces that were averaged over V1, M1 or RSP in the same mouse as in (**B**) across the duration (276 s) of the recording session. Both V1 and M1 exhibited notable oscillations with peak power around 6-7 Hz, and a second-harmonic was also apparent in V1. (**D**) Two example sequences of fluorescence images showcasing the spatiotemporal dynamics of visually evoked 3-7-Hz waves in PV cells of area V1 (successive image frames are 75 ms apart). Brain area boundaries, based on the Allen Brain Atlas (**Figure 4A** inset), are superposed onto the first image in each sequence. Oscillations also arose in M1 but were not time-locked to stimulus presentation (see (**F**)). (**E, F**) Visual stimuli consistently evoked 3-7 Hz and gamma (30–60 Hz) band oscillations in V1 but not M1 in the same mouse as (**D**). *Top* plots: Raster color plots showing fluorescence voltage signals, spatially averaged over V1 (**E**) or over M1 (**F**), revealing 3-7 Hz oscillations in both areas that were locked to stimulus-offset in V1 but not M1 (50 stimulus trials shown). *Bottom plots*: Raster plots of gamma-band filtered (30-60 Hz) PV cell voltage activity in V1 (**E**) and M1 (**F**), showing that gamma-band activity was evoked during stimulus presentation in V1 but not M1. (**G**) *Top trace*: Mean time-dependent activity trace (mouse 1), obtained by averaging the raw signals of (**E**) over all 50 trials. *Bottom 3 traces*: Analogous traces for 3 additional mice, one a PV-Cre mouse and the other two Cux2-Cre^ERT2^ mice. Shading: 95% C.I. (**H**) Mean time-dependent fluorescence signal magnitudes in the gamma-band (30-60 Hz) for all 4 mice in (**G**), determined by a wavelet spectrogram as in **Figure 4M**. All 4 mice showed significant increases in gamma band power during visual stimulation as compared to baseline values (p = 4 × 10^-19^, 6 × 10^-10^, 1.5 × 10^-16^ and 1 × 10^-17^ in mice 1–4 for the 100-ms-interval after stimulus onset, and 2.6 × 10^-18^, 3 × 10^-16^, 6.7 × 10^-16^ and 7 × 10^-15^ for the 0.5-s-interval at the middle of the stimulus period, for n = 50 stimulus trials, Wilcoxon sum rank test). Shading: 95% C.I. (**I**) Spatial maps of 3-7 Hz power, averaged across the entire recording sessions for mice 1 and 3 of panel (**H**) (recording durations of 275 s and 266 s, respectively). See (**D**) for brain area boundaries. (**J**) Maps of peak correlation coefficients, *r*, for the Cux2-Cre^ERT2^ and PV-Cre mice of (**H**), computed for each point in space by calculating the temporal correlation function between the local fluorescence trace and that at the center of V1 (black dots) and then finding this function’s peak value. (**K**) Maps of peak correlation coefficients, *r*, computed as in (**J**), except that correlations were computed relative to a point in M1 (black dots) not V1. Unlike in (**J**), in which coherent 3-7-Hz-band activity is largely restricted to V1, here the coherent activity is confined to near M1. Thus, the 3-7-Hz-band oscillations in M1 and V1 appear to be incoherent with each other. (**L**) Maps of visually evoked increases in the amplitude of neural activity in the gamma-band (35-60 Hz), computed by wavelet transform as in (**H**). Both L2/3-pyramidal and PV cell-types underwent increases in gamma activity that were localized to V1. (**M**) Two example sequences of fluorescence images from mouse 1 of (**H**), from videos taken at 300 Hz (3 ms between frames) to reveal the detailed progression of 3-7 Hz waves in V1, for 2 different wave events. (**N, O**) *Left panels*: Flow maps showing the local wave propagation directions (normalized to have the same amplitudes) for the same two 3-7-Hz wave events as in (**M**). *Right panels*: Histograms showing the distributions of propagation speed across the region in (**M**), for the same two events as in the left panels. *Insets*: Polar histograms of propagation direction for the two 3-7-Hz events, computed across the flow maps of the left panels, showing that the two different events had distinct directions of propagation across V1. The number values on each polar graph refer to the counts in each bin of the histogram. (**P**) Histograms of propagation speed, aggregated across spatial bins (62.5 µm wide) in V1 and 30 different 3-7 Hz waves in each of the same two mice as in (**L**). *Insets*: Polar histograms of the directions of 3-7 Hz wave propagation direction, showing that while individual waves had a clear direction of propagation (*e.g.* as in (**N, O**)), the distribution of these propagation directions across all 3-7 Hz waves was less uniform. (**Q**) Sequences of gamma-band (35-100 Hz) filtered images (3 ms between frames), from the same mouse as in (**M**), showing the spatiotemporal dynamics in V1 of two different gamma oscillation events. For display purposes only, the images shown were spatially low-pass filtered using a Gaussian filter (156 μm FWHM). (**R, S**) *Left panels*: Flow maps showing the local wave propagation directions during the same two gamma events as in (**Q**). *Right panels*: Histograms showing the distributions of propagation speed across the region shown in (**Q**), for the same two gamma events as in the left panels. *Insets*: Polar histograms showing the distributions of propagation direction for the two gamma events, computed across the flow maps of the left panels. (**T**) Histograms of propagation speed, aggregated across all spatial bins (62.5 µm wide) in V1 and 30 gamma waves in each of the same two mice as in (**L**). For both L2/3 pyramidal and PV cells, modal speeds of gamma waves were higher than those of 3-7 Hz waves, (**P**). *Insets*: Polar histograms of gamma wave propagation directions across all observed events, showing that while individual gamma waves had a clear propagation direction (*e.g.,* **R, S**), the distribution of these propagation directions across all gamma waves was far more isotropic.

To examine if other V1 neuron-types would exhibit similar dynamics, we characterized visually-evoked 3-7 Hz and gamma activity in somatostatin (SST) interneurons, layer 4 (L4) and layer 5 (L5) pyramidal cells (**Figure S7A**). The 3-7 Hz oscillations arose at visual stimulus offset in all these cell-types, albeit with slightly varying time-courses, but stimulus-evoked gamma activity in L4 pyramids was far weaker than in either L5 or L2/3 pyramids or the two interneuron classes and seemed temporally confined to stimulus onset and offset. These characterizations of sensory-evoked voltage dynamics in 5 V1 neuron-types should help refine models of visual cortical processing.

We next characterized the spatiotemporal patterns of the two types of waves in V1. The 3-7 Hz waves had speeds (108 ± 16 mm⋅s^-1^ for PV and 138 ± 16 mm⋅s^-1^ for L2/3 cells) and directions relative to the A-P axis (66 ± 8 deg for PV and 63 ± 10 deg for L2/3 cells) that were similar for the two cell-types (30 wave events each; mean ± s.e.m.; **Figure 5M–P**). Individual gamma waves generally traveled in a well-defined direction (**Figure 5R,S**), but the distributions of directions across all gamma waves were far more isotropic for both cell-types (**Figure 5T**). Gamma waves in PV and L2/3 pyramidal cells had much higher modal speeds (178 ± 85 and 145 ± 79 mm⋅s^-1^ respectively, mean ± s.e.m., 50 gamma events per cell-type) than 3-7 Hz waves (**Figures 5P,T**). By comparison, a subset of gamma-band activity arose concomitantly with and propagated similarly to 3-7 Hz wave activity, suggesting that, unlike the faster gamma waves arising in isolation, this slower subset of gamma-band activation was merely harmonics of 3-7 Hz activity (**Figure S6X,Y**).

### Locomotion-evoked theta and beta waves in hippocampus

To apply TEMPO imaging in a subcortical area, we studied the hippocampus during rest-to-run transitions. Past studies found locomotor-associated theta waves in hippocampus, but the use of electrode arrays or multi-site silicon probes led to relatively poor spatial resolution, the interpretative limitations of electric field potential recordings, and a lack of cell-type or neural pathway specificity.^74,91^

To overcome these limitations, we imaged ASAP3-expressing PV cells across a 3-mm-wide area of dorsal CA1. We took concurrent LFP recordings and directed an airpuff to the mouse’s back to elicit rest-to-run transitions (**Figures 6A–F, S2**). During running, ASAP3 and LFP signals evidenced a substantial rise in beta band power, accompanied by increased coherence between these two signal-types across most of the optical window (**Figures 6B–F, S7B-C**). The mRuby2 channel showed neither of these. Strikingly, we discovered that hippocampal beta oscillations are in fact traveling waves; we observed two orthogonal modes of beta wave propagation, in the CA3-to-CA1 and septo-to-temporal directions, with indistinguishable speeds (220 ± 4 *vs.* 220 ± 6 mm/s; median ± s.e.m; for n=428 and 199 waves propagating in the CA3-CA1 and septo-temporal directions, respectively, p=0.6; rank-sum test; **Figure 6G–J**).

**Figure 6:**
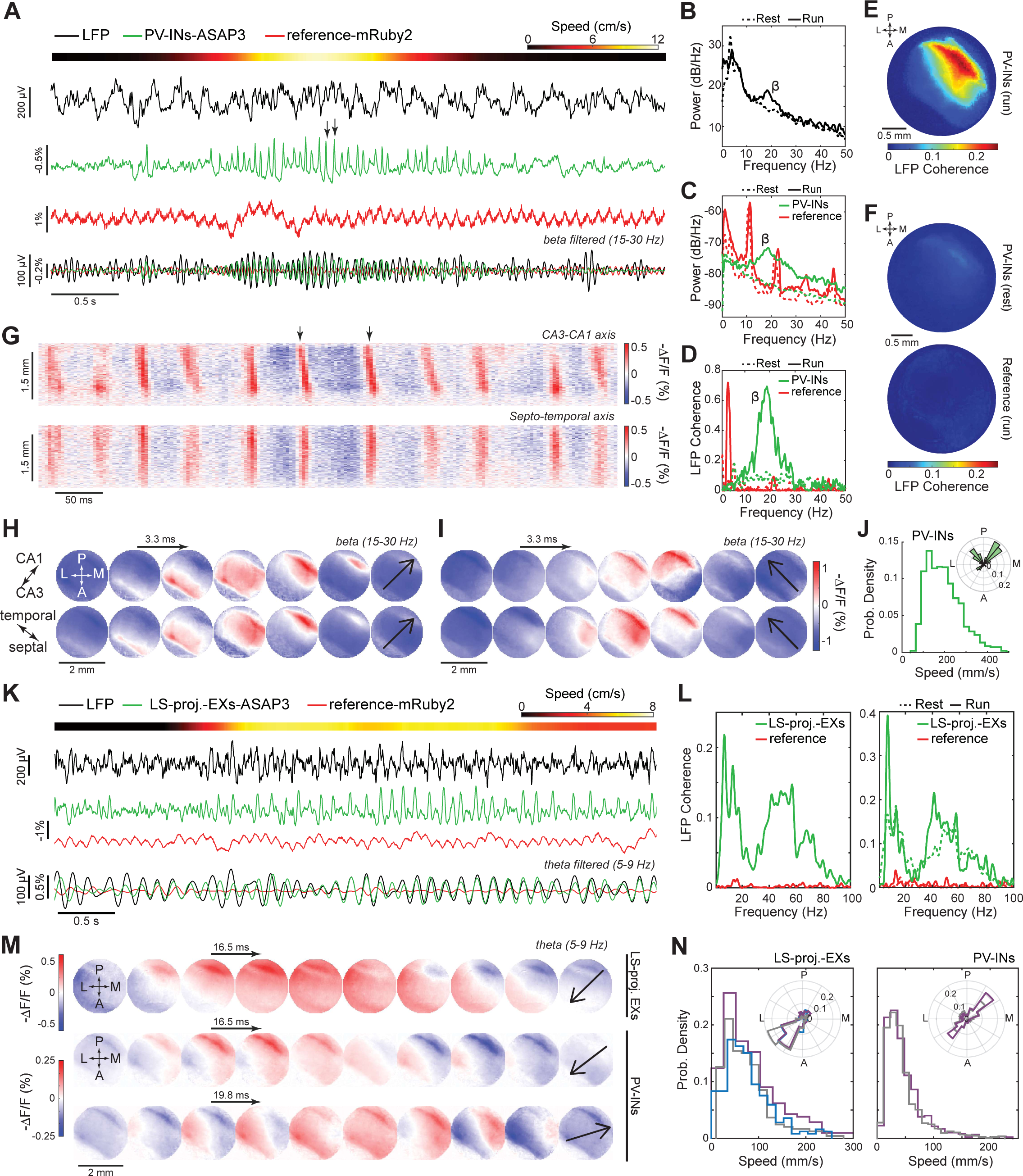
TEMPO imaging reveals locomotor-evoked bi-directional, theta- and beta-band traveling voltage waves in hippocampal PV interneurons and lateral-septum-projecting pyramidal neurons. (**A**) Plot of running speed in an example mouse (top), plus a set of concurrently acquired time traces (bottom 6 traces) illustrating locomotor-evoked hippocampal oscillations in the beta-band (15–30 Hz). Shown are broadband (top 3 traces) and bandpass-filtered versions (bottom 3 (overlaid) traces) of the hippocampal LFP (black traces), fluorescence voltage signals from ASAP3-expressing parvalbumin (PV) cells (green traces), and signals from the mRuby2 reference fluor (red traces). Fluorescence signals are averages across the entire visible portion of the hippocampus. During running, power in the beta band increased in both the LFP and ASAP3 traces but not the mRuby2 trace. Hemodynamic and motion artifacts are visible in the reference trace but not the ASAP3 and LFP traces. Two black arrows on the PV cell voltage trace mark events further characterized in (**G**) and (**H**). (**B, C**) Power spectral densities (PSDs) for the LFP (**B**) and fluorescence signals (**C**), for the same mouse as in (**A**) during 5 min of continuous recording, during resting (dashed curves) and running (solid curves) conditions. During running (defined as >2 cm/s speed), ASAP3 and LFP signals underwent power increases across the beta band. The mRuby2 signals had prominent spectral peaks arising from the heartbeat and its harmonics. Moreover, the mRuby2 PSD differs between resting and running states across nearly all frequencies, emphasizing the importance of correcting the GEVI signals for the changes in the reference channel. (**D**) Plots of the frequency-dependent coherence between the LFP and either the ASAP3 (green curves) or reference fluorescence signals (red curves) from the same mouse as in (**A**), during resting (dashed curves) and running (solid curves) conditions. ASAP3 but not mRuby2 signals underwent running-evoked increases in coherence with the LFP. The coherence peak near 0 Hz for mRuby2 during running reflects shared motion artifacts affecting the LFP and mRuby2 measurements but not the processed ASAP signals. Also see **Figure S7B-C** for differences along the CA1–CA3 axis in theta and beta power and coherence. (**E, F**) Spatial map of coherence (**E**) between the LFP and ASAP3 signals during running epochs across the hippocampal surface, averaged across the beta-band (15–25 Hz). The LFP electrode was located in CA1 (upper right portion of the map), which was, as expected, where coherence between the two measurements was highest. However, coherence values were significant across the full field-of-view, unlike values observed during rest (**F**; *top plot*) or for the fluorescence reference (**F**; *bottom plot*). (**G**) Space-time representation of PV voltage activity during beta waves, projected onto the CA3–CA1 (*top plot*) and septal-temporal (*bottom plot*) axes. Black arrows mark the same two voltage events marked in (**A**). The slope of each wave’s representation in the plots gives the reciprocal of that wave’s speed. Based on these slopes, one can see that the waves shown progress from CA3 to CA1 but exhibit almost no propagation in the septal-temporal direction. (**H, I**) Movie frames showing beta-frequency voltage waves traveling along the CA3-CA1, (**H**), or septo-temporal, (**I**), axes. Each row corresponds to one event; successive frames are 3.3 ms apart. The two events shown in (**H**) are marked with arrows in (**A**). Black arrows in the final frame of each plot show the direction of wave propagation, as determined computationally (**STAR Methods**). (**J**) Histogram of waves’ speeds and orientations (*inset*) for traveling beta waves (756 total events from n=2 mice, **STAR Methods**). Note that there are two main modes of wave propagation, along two orthogonal axes, the CA3-CA1 axis and the septo-temporal axes of the hippocampus. Beta waves traveled in these two directions with indistinguishable speeds (220 ± 4 mm/s and 220 ± 6 mm/s, median ± s.e.m, for n = 428 and 199 waves propagating along the CA3-CA1 and septo-temporal axes, respectively; p=0.64, rank-sum test). (**K**) Plots in the same format as those in (**A**), but with theta-band (5–9 Hz) filtered traces at bottom, for an experiment in which we used viral retrograde targeting to express ASAP3 in CA1 pyramidal neurons that had axonal projections to the lateral septum (LS). (**L**) Plots of coherence between the LFP and fluorescence signals from either LS-projecting pyramidal neurons (green curves) or the red reference fluor (red curves), computed across the entire recording from the same mouse as in **K** (*left*) or computed separately for resting and running epochs (*right*). Note that, unlike for PV-INs, for LS-projecting pyramidal cells the beta-band power is comparatively weak and is unmodulated by running; instead there is strong power in the theta- and gamma-frequency bands that is locomotor-dependent. (**M**) Movie frames showing traveling theta-frequency voltage waves for LS-projecting pyramidal (*top plot*) and PV inhibitory neurons (*middle and bottom*), as measured with ASAP3. Each row displays an individual theta wave event. Both neuron-types exhibited wave propagation in the CA1 to CA3 direction. Only PV neurons exhibited waves traveling in the CA3 to CA1 direction. Black arrows in the final frames of each plot show the direction of propagation, as computed computationally. (**N**) Histograms of speed and direction (*inset*) for traveling theta waves for LS-projecting pyramidal neurons (*left*; n=3 mice) and PV inhibitory neurons (*right*; n=2 mice). Note that only PV cells showed bidirectional wave propagation and that theta waves propagate with different speed between PV (40 ± 2 mm/s and 33 ± 1 mm/s, median ± s.e.m, for n = 930 and 1094 theta waves, for each 2 mice with PV cells targeting, respectively) and LS-projecting cells (90 ± 3 mm/s, 80 ± 3 mm/s and 84 ± 4 mm/s, median ± s.e.m, for n = 508, 465 and 441 theta waves, for each 3 mice with LS-projection targeting, respectively). Moreover, unlike beta waves, the two classes of propagating theta waves traveled with significantly different speeds in PV interneurons (35 ± 2 mm/s and 42 ± 2 mm/s, median ± s.e.m, for n = 777 CA3-to-CA1 and 637 CA1-to-CA3 theta waves, respectively, n=2 mice, p=0.0027).

To demonstrate a means of monitoring oscillations in a specific neural pathway, we virally expressed ASAP3 in CA1 pyramidal cells with axons in lateral septum (LS) (**Figure 6K**). In joint TEMPO and LFP recordings, LS-projecting pyramidal neurons exhibited broadband coherence with the LFP, which rose substantially in the theta (5-9 Hz) band during locomotion (**Figure 6L**). Beta-band coherence for LS-projecting pyramidal cells was weaker than for PV cells and was unchanged during locomotion.

We next compared the spatiotemporal dynamics of hippocampal theta waves in mice in which either PV interneurons or LS-projecting pyramidal cells expressed ASAP3 (**Figure 6M,N**). Both cell classes exhibited wave propagation in the CA1-to-CA3 direction, but only PV cells exhibited waves traveling in the reverse, CA3-to-CA1 direction. Interestingly, theta waves had different speeds in PV (36 ± 1 mm/s, median ± s.e.m; 2024 theta waves; 2 mice) and LS-projecting cells (87 ± 2 mm/s; 1414 waves; 3 mice). Moreover, unlike beta waves, for which speeds were statistically indistinguishable for the two directions of beta propagation, the two classes of theta waves in PV cells had statistically differentiable speeds (35 ± 2 mm/s and 42 ± 2 mm/s, median ± s.e.m, for n=777 CA3-to-CA1 and 637 CA1-to-CA3 waves, respectively; p=0.003; rank sum test).

Altogether, the discoveries that beta oscillations are actually traveling waves with two orthogonal modes of propagation, and of a previously unreported direction of travel for hippocampal theta waves, illustrates how TEMPO imaging can reveal the spatiotemporal patterns of voltage activity in specific neuron-types or projection pathways.

### Concurrently imaged, visually evoked spatiotemporal dynamics of excitatory and inhibitory cells in awake mice

To study visually evoked activity in ASAP3-expressing PV cells and Varnam2-expressing pyramidal cells in awake mice, we performed dual cell-type imaging with the TEMPO mesoscope across the right cortical hemisphere as mice viewed drifting grating stimuli with their left eye (**Figure 7A–F**). In V1, visual stimuli consistently evoked 3-7 Hz oscillations in both cell classes after stimulus offset (**Figure 7G–J**). In line with our earlier studies (**Figures 3,5,S2**), we observed stimulus-evoked gamma band increases only in the ASAP3-labeled cell class, here the PV cells (**Figure 7K**).

**Figure 7:**
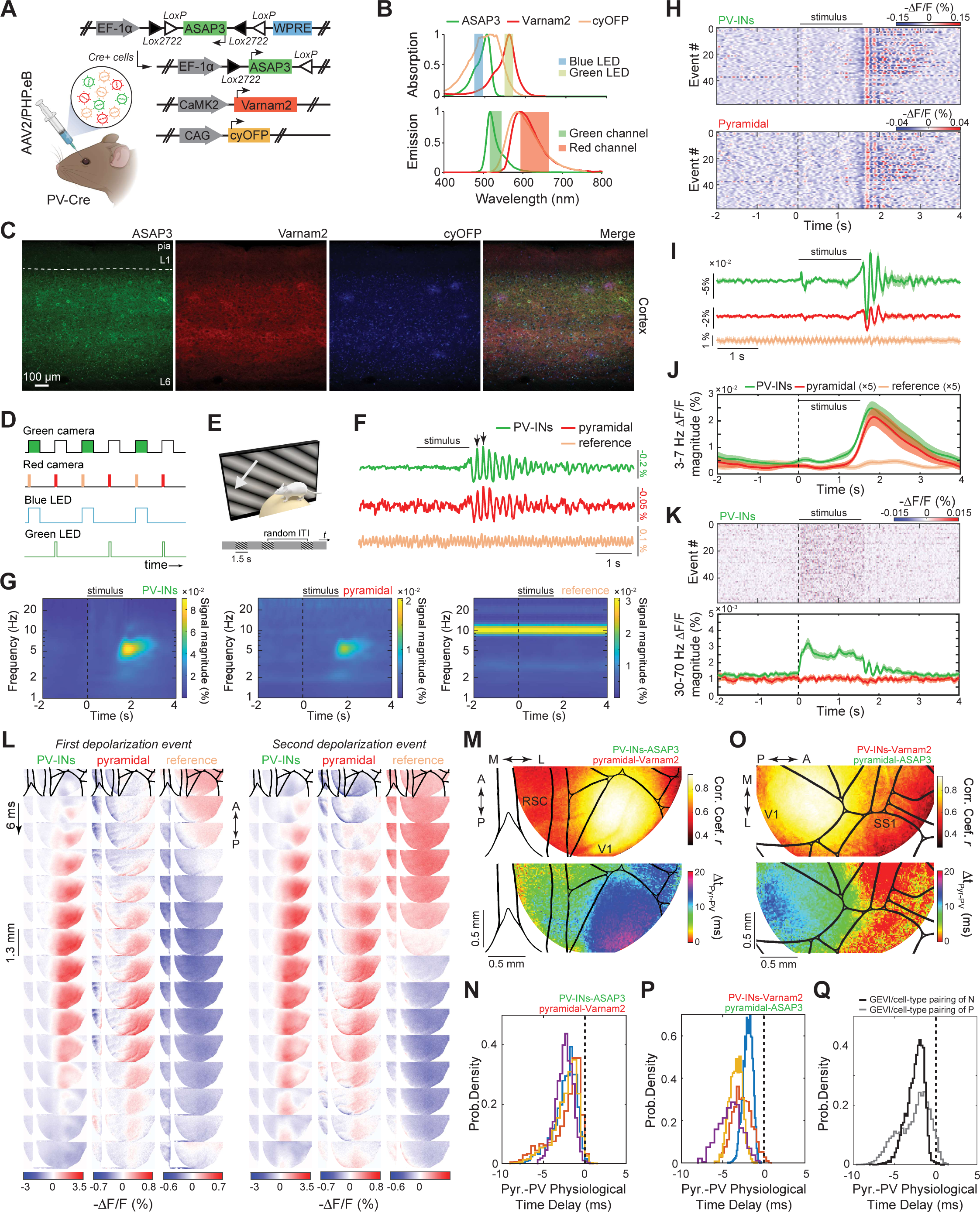
Concurrent voltage dynamics of two neuron-types observed with TEMPO imaging in behaving mice. (**A**) To image the voltage dynamics of two neuron-types at once, we expressed the 3 fluorophores using the same 3 viruses as for uSMAART sensing in 2 cell classes (**Figure 3A**). (**B**) Absorption and emission fluorescence spectra for the 3 fluorophores, shown along with emission spectra of the two light-emitting diodes (LEDs) and passbands of the emission filters in the TEMPO mesoscope (**Figure 4A**). (**C**) Fluorescence confocal images of a brain slice from a mouse expressing ASAP3 in PV cells, Varnam2 in pyramidal cells and cyOFP in all cell-types. The far right image is an overlay of the preceding images, highlighting the different targeting of all three proteins. L1: layer1, L6: layer 6. (**D**) Schematic of the timing protocol used for dual neuron-type recordings. The durations of image frame acquisitions by the two cameras are depicted via digital voltage pulses in the top two traces. Colors within each voltage pulse denote the colors of the acquired fluorescence signals. Periods of LED illumination are depicted via the voltage pulses in the bottom two traces. When the blue LED is illuminated, one camera captures ASAP3 fluorescence and the other camera captures cyOFP fluorescence. When the green LED is on, the second camera captures Varnam2 fluorescence. This scheme allows the 3 different fluorescence signals to be captured unambiguously using only 2 illumination and 2 detection pathways. (**E**) The visual stimulation paradigm was the same as in **Figure 3B**. (**F**) Example time traces of visually evoked membrane voltage dynamics in ASAP3-expressing PV interneurons and Varnam2-expressing pyramidal cells, spatially averaged over primary visual cortex (V1). The visual stimulus was presented during the interval marked with the black horizontal bar. Whereas both neuron-types exhibited 3-7 Hz voltage oscillations after stimulus offset, only heartbeat artifacts were observed in the reference cyOFP channel. Depolarization events marked with arrows are further characterized in (**L**). (**G**) Event-related wavelet spectrograms for PV interneurons, pyramidal cells and reference signals for the mouse in (**F**), averaged over the same 60 trials as in (**H**). 3-7 Hz oscillations arose at stimulus offset in both cell-types; the reference plot shows strong heartbeat artifacts (∼10 Hz) that were successfully unmixed from both voltage channels. (**H**) Raster plots of visual stimulus-evoked voltage dynamics in V1. Visual stimuli consistently evoked 3-7 Hz oscillations in PV (top) and pyramidal cells (bottom) after stimulus offset. Each row shows data from one of 60 different trials in the same mouse as in (**F**). (**I**) Mean time-dependent fluorescence traces, obtained by averaging the signals from all 3 fluors across all 60 trials of (**H**), showing the increase in 3-7 Hz power at stimulus offset. Shading on traces in (**I–K**) denotes 95% C.I. (**J**) Mean time-dependent fluorescence signal magnitudes in the 3-7 Hz band, as obtained from a Hilbert transform, showing the rise in oscillatory power after stimulus offset. (**K**) *Top*: Raster plots of PV cell voltage dynamics in the gamma band (30–70 Hz) across the same 60 trials as in (**H**). *Bottom*: Mean time-dependent fluorescence signal magnitudes in the gamma band for all 3 fluors, as determined from a Hilbert transform. PV cells exhibited a clear increase in gamma-band power during visual stimulation, in contrast to the increase in 3-7-Hz-band power observed after stimulus offset. (**L**) Movie frames of fluorescence emissions showing the voltage depolarization events marked with arrows in (**F**), for each of the 3 fluors. Brain area boundaries, based on the Allen Brain Map, are above each column and delineate areas labeled in (**M**). Both PV and pyramidal cells underwent depolarizations, but the reference cyOFP signals appear unrelated to the voltage dynamics of either neuron class. (**M, N**) Spatial maps (**M**) of the correlation coefficient (*top plot*) and time delay (*bottom plot*) between excitatory and inhibitory neuronal activity in the 3-7-Hz-band for every wave event detected (n=98 visual stimulation trials). Correlations in stimulus-evoked, 3-7 Hz activity between the two neuron-types were highest in visual cortex and accompanied by a greater temporal shift between the oscillations of the excitatory and inhibitory cells. Histograms in (**N**) show distributions of the physiological time delays between the two neuron classes, estimated in visual cortex based on 4 imaging sessions (denoted by curves in 4 separate colors) from 3 mice, as corrected computationally for the time-lags induced by the indicators’ kinetics. To estimate these time-lags, we parametrically fit their values using the measured time-delays in (**M**) and 6 other complementary experiments with various GEVI and cell-type assignments (**Figure S7D-F**). We repeated this procedure for each of the 4 recordings to generate the 4 histograms in (**N**). (**O, P**) Same as (**M**) and (**N**), respectively, but for the reversed labeling strategy in which pyramidal neurons express ASAP3 and PV interneurons express Varnam2. Each colored histogram in (**P**) is for one of 4 different recordings from 4 different mice. (**Q**) Histograms aggregating the data from each of (**N**) and (**P**), showing the estimated physiological time delays were statistically indistinguishable for the two labeling strategies [time delay: 2.27 ± 0.02 ms (median ± s.e.m.), averaged over V1 and 332 wave events in (**N**); 2.32 ± 0.01 ms, averaged over 230 wave events in V1 in (**P**); rank-sum test p = 0.08].

Since 3-7 Hz waves appeared in both cell classes concurrently (**Figure 7L**), we computed spatial maps of the empirically determined correlation coefficients and temporal delays between the waves in each cell class (**Figure 7M,O**). As expected, correlation coefficients were highest in visual cortex, but the measured temporal delays differed when we reversed the assignments of the two GEVIs to the two neuron classes (**Figure 7M,O**). This dependence on indicator assignments implied that one or both GEVIs likely induced an apparent temporal delay due to fluorescence voltage-signaling kinetics that were insufficiently fast to report the wave’s instantaneous phase. Motivated by this inference, we characterized the temporal delays induced in visual cortical PV and pyramidal cells by 3 GEVIs used in this paper, which indeed were nonzero for Varnam2 and Ace-mNeon2 but near zero for ASAP3 (**Figure S7D–F**). Having determined the delays induced by the GEVIs, we then estimated the true physiological delays between visual cortical PV and pyramidal cells during 3-7 Hz waves (**Figure 7N–Q; STAR Methods**). Satisfyingly, the values of these estimates were statistically indistinguishable irrespective of the assignments of ASAP3 and Varnam2 to PV and pyramidal cells (2.27 ± 0.02 ms *vs*. 2.32 ± 0.01 ms for n=332 events in **Figure 7N** and 230 events in **7P,** respectively; median ± s.e.m., 3 mice each; p = 0.08; rank-sum test) and show that pyramidal cell depolarization preceded that of the PV cells. This result is consistent with those from *in vivo* patch clamp studies of visually evoked 3-7 Hz oscillations in awake mice^59^ and provides a constraint on candidate mechanisms for wave propagation.

## DISCUSSION

TEMPO measurements involve cell-type specific expression of one or more GEVIs, a reference fluor to track biological and instrumentation artifacts, a dual-channel fluorescence sensing apparatus, and analytics to unmix artifact and GEVI signals. We made advances in each of these aspects, as discussed below. Together, these strides enabled the first voltage-sensing and -imaging systems capable of detecting high-frequency voltage dynamics of targeted neural populations in freely moving and head-fixed behaving animals, and in real-time without trial-averaging. This yielded the first recordings of beta and gamma rhythms in genetically targeted neuron-types, opening the door to mechanistic dissections of different cell-types’ contributions to the generation of these oscillations.

Further, the sensitivity of TEMPO imaging allowed observations of delta, theta, beta and gamma traveling waves at the single-trial level; this showcases the technique’s fine spatial resolution and is a key advance, since averaging the dynamics of many waves traveling in distinct directions will yield a very different, potentially misleading, picture of brain dynamics.^92,93^ To this point, we characterized the statistical distributions of wave propagation directions to an extent and accuracy level that has not been feasible with electrode arrays. We also uncovered previously undocumented traveling waves, hippocampal beta waves, and, unexpectedly, that propagation directions for both hippocampal theta and beta waves are bidirectional. The speeds of the waves we observed in awake mice ranged from ∼0.1–0.5 m⋅s ^-1^, across cell-types and brain areas. These values are ∼10–100 fold slower than conduction speeds of myelinated axons but consistent with those of unmyelinated axons and the delays incurred during generation of synaptic potentials.^94,95^ While each of the traveling wave forms seen here needs to be studied further, our TEMPO methods will facilitate studies of both their physiological mechanisms and functional roles.

Another key facet of our advances is the capability for dual cell-type recordings. Studies of excitatory (E) and inhibitory (I) neurons via electrical recordings often rely on classifying cell-types by their distinct spike rates, action potential waveforms, and refractory periods. However, differentiating neural subclasses (*e.g.* subtypes of inhibitory cells) is usually infeasible with electrical recordings alone. Although subtype identification during extracellular electrode recordings can be done via optogenetic activation, this method neither generalizes easily to more than one cell-type at once nor reports subthreshold voltage dynamics.^96,97^ TEMPO measurements offer these benefits and can thereby yield unique insights. To illustrate, we studied how locomotion alters the coherence of hippocampal PV and pyramidal cells with the LFP across different layers of CA1, and how sharp wave ripples engage these two cell-types (**Figure 3N–Q**). We also studied the inter-relationships between E and I activity in the visual cortex of awake mice (**Figures 3A–I, 7**).

### Fluorescence labeling

To express the TEMPO fluorescent proteins, we created complementary transgenic and viral labeling methods. The Ai218 TEMPO reporter mouse line is suited for labs that seek an easy labeling method or that lack access to or wish to avoid viral techniques. Ai218 can be crossed with Cre-driver mouse lines to achieve cell-type specific, homogeneous expression of Ace-mNeon1 and mRuby3, free from the intrinsic variability of viral labeling. However, Ace-mNeon1 is no longer the best GEVI, and with Ai218 it was more challenging to detect high-frequency activity (**Figure S6M–S**). For brain-wide (or local) expression of more recent GEVIs, PHP.eB-mediated viral labeling enables TEMPO measurements with higher sensitivity than feasible with Ai218 mice.

### Reference channel

In early forms of fluorescence fiber photometry, invented ∼50 years ago to track biological signals via optical fiber,^98^ the use of a reference channel to account for signal artifacts was commonplace. More recent studies have often^53,54,99^ but not always^55^ omitted the reference channel. Similarly, brain-imaging studies in head-fixed rodents have sometimes used an indicator isosbestic point or multiple excitation wavelengths to track and remove hemodynamic artifacts.^100,56,101^ Overall, the absence of a reference channel can be deeply problematic when the signal artifacts are near in magnitude to the signals of interest. As most GEVIs lack a convenient isosbestic point, TEMPO uses a spectrally separable, non-functional fluorophore as a reference.^7,9,82^ For dual cell-type TEMPO, we created a new reference approach using a long Stokes-shift fluorophore. Consistent with our and other prior reports^8,101,102^, we observed here that the reference channel captures brain motion artifacts (*e.g.* at running onset, **Figure 6A**), seconds-scale hemodynamic responses (**Figure 3D**), and heartbeat (**Figure S5B,E**).

To further aid removal of artifacts from the GEVI traces, we created a convolutional filtering method to unmix artifacts in a frequency-dependent manner; unlike prior methods, this approach accounts for the variable impact of different noise sources on the GEVI channel and thereby avoids adding reference channel noise into the GEVI traces (**Figure S5J**). The algorithm is as integral to the TEMPO toolbox as the new instrumentation and expression strategies introduced here and, moreover, is broadly applicable to other fluorescence sensing modalities in biomedicine.

### Comparison of TEMPO methods to prior voltage-mapping techniques

uSMAART is a fiber-optic embodiment of TEMPO that achieves a ∼10-fold improvement in instrumentation sensitivity, which in turn enables the first optical studies of high-frequency voltage activity, in moving animals and without trial-averaging. The central engineering insight underlying this ultrasensitive form of fiber photometry is the recognition that, to capitalize on the superior stability of high-frequency-modulated laser illumination over LED illumination, the apparatus must include a means of breaking the illumination coherence to prevent fiber-optic mode-hopping fluctuations when the animal actively behaves. When the ∼10-fold improvement provided by uSMAART is combined with the ∼10-fold superiority of ASAP3 over prior GEVIs for tracking subthreshold dynamics, we realized an overall sensitivity improvement of ∼100-fold over prior fiber-optic voltage-sensing studies.^7,9^

To map electric field potentials across the brain surface, prior work used voltage-sensitive dyes (VSDs), GEVIs, or ECoG electrode arrays.^88,89^ ECoG recordings usually have better time-resolution but much poorer spatial resolution than imaging methods.^40^ ECoG signals are also difficult to interpret in terms of neurophysiological signal sources owing to volume conduction and lack of cell-type specificity. Imaging studies with VSDs or GEVIs can map neural voltage signals across ∼100 mm^2^ of tissue at a spatial resolution more than 10-fold finer than that of state-of-the-art ECoG electrode arrays.^46,47^ However, VSD imaging also lacks cell-type specificity and often is highly prone to photobleaching and phototoxicity that sorely limit recording durations.

Our mesoscope is the first imaging instrument to embody the TEMPO approach and incorporates several innovations to visualize population voltage dynamics in genetically targeted cell-types across areas up to 8 mm wide in head-fixed animals (**Figure 4A,B**). The mesoscope reveals traveling voltage waves in a way that has never been possible before, either with electrode arrays, which sample discrete sets of points, or with prior voltage-imaging methods, which lack the sensitivity to observe the brain’s intrinsic high-frequency rhythms. Although one recent GEVI imaging study used large-amplitude sensory stimulation to evoke a high-frequency neural voltage oscillation at the same frequency,^82^ this differs markedly from the capability to observe the brain’s self-generated beta and gamma waves, as shown here. Showcasing its capabilities, our initial studies with the TEMPO mesoscope provided the first movies of cross-frequency coupling, enabled the first pathway-specific recordings of neural voltage oscillations, revealed that hippocampal beta oscillations are in fact traveling waves, and uncovered bidirectional propagation of hippocampal waves in both the theta and beta frequency bands.

Whereas high-speed (∼1 kHz) voltage-imaging studies of individual neural spikes are typically limited in mammals to a few minutes per imaging session due to photobleaching,^1,4–6^ both TEMPO modalities use far lower illumination intensities (∼2 mW/mm^2^) and allow hour-long studies with minimal bleaching and phototoxicity. Unlike fluorescence Ca^2+^ signals, TEMPO reports depolarizations and hyperpolarizations equally well; this aspect should be especially useful for studies of non-laminar brain areas, which often have multiple, interspersed cell-types that make LFP signals hard to interpret.^40^ Further, both uSMAART and the TEMPO mesoscope are compatible with fluorescent reporters beyond GEVIs, such as Ca^2+^, neuromodulator, neuropeptide or neurotransmitter indicators. While Ca^2+^-related fluorescence signals usually have high dynamic range, neuromodulator and neuropeptide indicators often produce smaller signals;^53,99^ thus, studies with the latter should benefit greatly from the instruments and unmixing methods introduced here.

### Cross-frequency coupling in targeted neuron-types

Using TEMPO’s ability to detect oscillations up to 100 Hz, we found the first instances of cross-frequency coupling (CFC) in the transmembrane potentials of genetically identified cell-types. These include interactions in PV cells between neocortical delta and gamma rhythms in anesthetized mice, and between hippocampal theta and gamma during locomotion. Notably, the former gamma waves traveled medio-laterally, orthogonal to the concomitant delta waves traveling anterior-to-posterior, showing the different rhythms engaged in CFC may travel in distinct directions. Our ability to resolve CFC in specific cell-types allows mechanistic studies of such phenomena.

Many past studies of CFC, including in human subjects, used field potential recordings, which likely captured contributions from multiple cell-types underlying the phenomenon. Other studies assessed CFC through joint LFP and extracellular electrical recordings of spiking by identified neuron-types.^103–107^ Here, the CFC signatures found with TEMPO reflect the transmembrane potential dynamics of individual cell classes, in which CFC may arise due to complex interactions between excitatory and inhibitory input signals and spiking outputs.

In all studies of CFC, it is crucial to differentiate true CFC from coupling between the different harmonics of an individual oscillation.^108^ The latter is a mathematical effect intrinsic to non-sinusoidal oscillations, whereas the latter reflects *bona fide* interactions between two physiologically distinct oscillations. The two phenomena can be distinguished, as coupled harmonics should co-propagate with the same speed and direction. For instance, we observed many instances of gamma band activity locked to a 3-7 Hz wave, traveling with the same speed and direction as its 3-7 Hz carrier (**Figure S6X,Y**). By comparison, isolated gamma waves were faster and more isotropic in their travel (**Figure 5T**). Identifying these features may require the spatial resolution of TEMPO imaging.

### Traveling voltage waves in visual cortex

We studied anesthesia- and sensory-induced gamma waves in visual cortex and characterized their spatiotemporal features across brain states. Individual gamma waves traveled mainly unidirectionally (**Figures 4P,Q** and **5R,S**), whereas the distribution of these propagation directions was nearly isotropic in awake but not anesthetized mice (**Figures 4R, 5T**). This is suggestive of different propagation mechanisms^109^ and supports the idea that the repertoire of accessible neural dynamics varies with brain state.^110,111^ Visually evoked gamma waves have been seen in electrical recordings in primate visual cortex with substantial heterogeneity of propagation directions.^112,113^ Here, the extent of visually-evoked gamma activity varied between neuron classes, with layer 4 pyramids showing far less activation than either layer 5 or layer 2/3 pyramids or PV or SST interneurons (**Figure S7A**).

In addition to gamma waves that were evoked during visual stimulation, we observed visual cortical 3-7 Hz waves at the offset of visual stimulation. This 3–7 Hz activity is thought to require thalamocortical interactions^61^ and might be a precursor of the human alpha rhythm that supports attentional processing.^60,61^ Prior electrical studies of this sensory-evoked oscillation examined different neuron classes, one cell at a time,^59^ or used ECoG electrode arrays^114^, which yielded similar wave speed estimates as those here but lacked cell-type specificity.

Notably, our dual cell-type imaging studies revealed that visual stimulation offset evokes simultaneous 3-7 Hz rhythms in pyramidal and PV cells, but with pyramidal cell activation preceding that in PV cells by ∼2 ms (**Figure 7N–Q**). This time delay estimate was unchanged when we swapped GEVI assignments between the two neuron-types; however, it was important to account for delays in fluorescence signaling of wave activity due to voltage-dependent kinetics of the two FRET-opsin GEVIs, Varnam2 and Ace-mNeon2, which were not fast enough to report the wave’s instantaneous phase (**Figure S7D–F**). By comparison, ASAP3 had near-zero signaling delays. The delays resulting from FRET-opsin kinetics differed between cell-types, reflecting that the GEVI time-constants themselves vary with voltage^1,2,8,58^ and thus any differences in resting potentials or oscillatory amplitudes between the neuron-types will lead to different values of the time-constants. While we expect additional fast GEVIs with near-zero signaling delays, like ASAP3, will emerge with further GEVI engineering, users seeking to explore the fine timing relationships between different neuron-types should take care as we did to characterize any GEVI-induced temporal shifts.

### Hippocampal theta and beta oscillations are bidirectional traveling waves

We studied locomotor-evoked voltage activity in PV and pyramidal neurons in the CA1 subfield of dorsal hippocampus. Using both TEMPO instruments, we found locomotor-evoked theta (5-9 Hz; **Figures 1D–I**, **3M–O, 6K–N**, **S3C–E**), beta (15-30 Hz; **Figures 1D–I**, **3M–O, 6A–J**, **S2J–L, S3C–E**), and gamma (30–70 Hz; **Figures 1E–I**, **2G–J**, **6L**, **S3E**) activity in freely moving and head-fixed behaving mice. In both neuron-types, theta activation was coherent with the LFP across all hippocampal laminae. Whereas, beta and gamma activation cohered most strongly with LFP signals in *stratum radiatum* for both PV and pyramidal cells. Within the theta rhythm, PV cells exhibited nested gamma (40–70 Hz) activity that arose at the peak of the LFP theta oscillation (**Figure 2G–J**). This form of hippocampal theta-gamma coupling is well known from electrical studies.^115^ PV cells also had especially prominent, locomotor-evoked beta activity, which appeared in the LFP recordings (**Figures 1E,F,H**; **3M–O**; **S2J,K**) but not as strongly as in the PV TEMPO signals. A combination of physiological and experimental factors likely explain this. Reflecting their fast-spiking characteristics, PV interneurons may exhibit locomotor-evoked beta and theta oscillations in different relative proportions than other neuron-types, such as pyramidal cells, that make prominent contributions to the LFP signal. PV cells are also relatively sparse and have modestly sized dendritic trees and thus modest dipole moments, implying that LFP signals may be less sensitive to their activity patterns than those of denser neurons with larger dendrites. Further, unlike for theta activity, detection of beta activity was more sensitive to location along the CA1-to-CA3 axis (**Figure S7B,C**) and an electrode’s laminar position (**Figure 3M**). Unlike gamma activity, PV cell beta activity did not appear to be nested within theta activity and, while theta and beta rhythms were both generally locomotor-evoked, their moment-by-moment amplitudes seemed to vary unrelatedly (**Figure 1D**), in line with past reports.^116,117^

While hippocampal beta oscillations have been reported before^116,117,118^, our imaging studies provide the first evidence these rhythms are in fact traveling waves. Strikingly, beta waves traveled along a pair of orthogonal directions, namely the CA1-to-CA3 and septo-to-temporal directions (**Figure 6J**). The presence of hippocampal beta activity^118^ during exploration of novel environments^117^ and olfactory associative learning^118–120^ argues for its behavioral relevance.

We also imaged the propagation dynamics of theta waves in CA1, in both PV-interneurons and LS-projecting pyramidal neurons. Our studies of the latter neuron class constitute the first recordings of pathway-specific neuronal voltage oscillations. Theta waves in LS-projecting neurons traveled approximately aligned to the CA1-to-CA3 direction, consistent with electrophysiological analyses of theta wave propagation.^74,91^ Unexpectedly, in PV cells, we observed bidirectional propagation of theta waves in both this and the reverse direction (**Figure 6N**). These results reveal a new class of theta waves that travel in the CA3-to-CA1 direction and suggest the LS-projecting subset of CA1 pyramidal cells might only participate in theta waves traveling CA1-to-CA3.

Overall, our findings of previously undocumented directions of travel for both hippocampal theta and beta waves raise important questions for future work about the underlying mechanisms and likely distinct functional roles of waves traveling in each anatomic direction, such as has been suggested for various neocortical waves and oscillations.^114,121^

### Limitations of the study

TEMPO offers insights that complement those from LFP, ECoG or EEG recordings. TEMPO and field potential signals have distinct physiological sources and thus, by design, will reveal different aspects of the brain’s electrical dynamics. Unlike LFP measurements, which probe extracellular potentials and thus are influenced by volume conduction as well as the composition or laminar organization of nearby tissue, TEMPO reports local transmembrane voltages and so is unaffected by the above tissue factors that can complicate interpretations of LFP recordings.^40^ Nonetheless, to be conservative in our interpretations of TEMPO signals, we applied the criterion that they should be coherent with LFP signals, above levels observed for the reference fluor.

In principle, this criterion could lead us to overlook *bona fide* TEMPO voltage signals that might be incoherent with the LFP. For instance, some forms of neural activity, perhaps in sparse neuron-types or reflecting asynchronous synaptic dynamics, might not be synchronized with the larger-amplitude network dynamics captured by the LFP recording. Alternatively, some weak oscillatory signals might only attain statistical significance after averaging spectral power increases over multiple experimental trials, and this averaging could mask coherence with the LFP.

However, there are good reasons to be cautious when interpreting TEMPO signals that are incoherent with the LFP. While our convolutional filtering approach is highly effective at unmixing physiological artifacts and other noise sources from the voltage traces, including artifacts with non-stationary amplitudes, it is conceivable that a physiological artifact could additionally be non-stationary regarding the frequency spectrum over which it appears in the voltage channel. The convolutional filter, once determined from the data, has a stationary form and will be unable to account fully for physiological artifacts that appear with different frequency signatures at different epochs in the recording, especially if their frequency spectrum overlaps that of other artifacts. This would leave some artifacts remaining in the unmixed voltage trace, and these residual artifacts would very likely be incoherent with the LFP.

Another interpretative issue for TEMPO studies is that, as with field potentials, there are complexities relating to where signals originate within neurons. The TEMPO signal is a weighted sum across all compartments of all GEVI-labeled cells in the recording volume, with the weightings influenced by the relative amounts of GEVI-labeled cell membrane area and proximities to the objective lens or fiber-optic tip. Dendrites and axons typically make the largest contributions to the net area of GEVI-labeled membranes. However, this does depend on patterns of GEVI trafficking, and whether a GEVI preferentially labels the soma, axon or dendrites will impact the relative labeling densities and contributions to the TEMPO signal from these compartments. Further, fluorescence signals from neural compartments further away from the optical collection apparatus will be attenuated in comparison to signals from closer compartments. But, for the fluorescence photons that are collected, the absence of optical sectioning in our instruments obscures the precise spatial origins of the optical signal. Without cellular resolution, TEMPO measurements report aggregate voltage changes across a targeted cell population; this differs from LFP recordings, in that with TEMPO the cell-type is user-selected, but is similar to LFP recordings in that one cannot distinguish large voltage changes in a modest set of cells from modest changes in a large cell set. GEVIs also vary in their capabilities to report subthreshold *vs.* spiking activity. In our work, we favored ASAP3 due to its ∼10-fold better sensitivity to subthreshold activity (**Figure S1**), but this does not exclude the possibility of spiking contributions to the TEMPO signal. Finally, since GEVIs are incorporated into the cell membrane, it should be noted that, especially at high levels of GEVI expression, alterations to transmembrane potential dynamics are conceivable.

### Technological Outlook

TEMPO will benefit from technical advances of several kinds. While most protein engineering efforts to improve GEVI signaling have focused on neural spike detection, creating separate GEVIs with superior voltage-sensitivity and speed at subthreshold voltages will be important to further enhance TEMPO’s sensitivity to subthreshold activity and to reduce temporal delays arising from GEVI time-constants. Further, given the amelioration of optical mode-hopping noise achieved here, the dominant remaining noise in our uSMAART apparatus appears to be from autofluorescence. Future studies could benefit from autofluorescence-free optical fibers and fluorescence lifetime-based separation of autofluorescence from GEVI signals.^49,122^ For TEMPO imaging studies, scientific-grade image sensor chips with more pixels and larger well capacities will enable wider fields-of-view and better sensitivity for detecting high-frequency voltage waves.

Other improvements could allow TEMPO studies in additional optical formats. Future versions of uSMAART might use high-density multi-fiber probes to study multiple brain areas at once.^51^ To access cells in deep cortical layers or brain regions, TEMPO imaging could be performed via optical microprisms^123^ or gradient index (GRIN) microendoscopes.^124^ Lastly, new viruses with engineered capsids,^106,107^ genetic tagging methods, and cell-type specific enhancers^125^ should expand TEMPO’s utility for studies of precisely defined cell classes and diverse species.

### Scientific Outlook

TEMPO is well poised to have a major impact on basic and disease-related neuroscience. The brain’s electric field dynamics remain poorly understood at a mechanistic level, and, for many types of oscillations, the neuron classes that drive the rhythm remain unknown or under debate.^35–37^ This has stymied interpretations of clinical electrophysiological recordings in terms of the activity of specific cell-types. Combined TEMPO and electrophysiological recordings (of field potentials and/or spiking activity) in normal and disease model animals should help break this impasse by revealing the neuron classes underlying specific EEG, ECoG or LFP signatures. Specifically, TEMPO can provide insights regarding the generation of electric field potentials by revealing which neuron-types are active in different neurophysiological states, the frequency bands in which each cell-type exhibits its activity, and the levels of coherence between TEMPO and LFP signals. Such joint optical and electrical studies may benefit from the use of transparent, graphene-based electrodes^126,127^, which can be placed within the optical field-of-view without occluding image collection. With distinct GEVI classes that signal supra- or sub-threshold activity, respectively, neuroscientists could dissect how the spiking and subthreshold dynamics of each cell-type relate to extracellular potentials. Dual cell-class TEMPO recordings will also enable studies of the real-time interactions between neuron-types. These efforts, in turn, should lead to fundamental advances in understanding the biophysics of electric field generation and to new electrophysiological biomarkers of brain disease.

As showcased by our discovery of new forms of traveling voltage waves (**Figure 6**), TEMPO also has the capacity to reveal previously unknown electrophysiological phenomena and offers new opportunities to examine the roles of propagating waves in neural computation. Many traditional neural network models emphasize network topology, as expressed in a matrix of synaptic weights.^128,129^ However, the possible role of traveling waves in neural computation^88,130^ suggests the geometric aspects of network connectivity may also shape information processing. We found that visually evoked cortical 3-7 Hz waves had a consistent propagation direction, but the directions of gamma waves were isotropically distributed (**Figure 5**). These results are suggestive of a mechanism for routing information through different pathways,^131^ a concept that merits experimental and theoretical exploration. By reporting neural oscillations up to ∼100 Hz, TEMPO will reveal contributions of different neuron-types to the brain’s rich set of high-frequency oscillations and waves and their roles in diverse forms of behavior, in health and disease.

## ACKNOWLEDGEMENTS

This study was supported by NIH BRAIN grants U01NS120822 (M.J.S. and G.V.), UF1NS107610 (M.J.S. and H. Z.), U19NS104590 (M.J.S. and I. Soltesz), 5U01NS10346403 (M.Z.L.), NSF NeuroNex grant DBI-1707261 (M.J.S. and K. Deisseroth), and by NCRR grant 1S10OD010580-01A1 to the Stanford Cell Sciences Imaging Center (RRID:SCR_017787), which provided training and usage support. We thank G.Y. Yin, B. Conrad, O. Hazon, T. Tasci, and J. Thompson for technical support. We thank X. Sun and M. Redinbaugh for their comments on the manuscript.

## AUTHOR CONTRIBUTIONS

Y.Z. performed viral vector subcloning, DNA preparations, virus testing and commissioning. Ja.L. bred and genotyped transgenic mice. T.D. and H.Z. created the Ai218 mouse line. Y.Z. and S.H. designed and performed mesoscope TEMPO surgeries in cortex and hippocampus, respectively. S.H. acquired and analyzed mouse behavioral data. Ja.L. and S.H. performed brain slices and confocal imaging, respectively. S.H. and R.C. designed, built and did the instrumentation programming for uSMAART and the TEMPO mesoscope, respectively. V.K. developed the convolutional unmixing algorithm. S.H. designed, performed and analyzed fiber-optic TEMPO and LFP experiments. S.H. designed mesoscope TEMPO experiments. S.H. and R.C. acquired mesoscope TEMPO data. S.H., R.C., V.K., Ji.L. and G.D. analyzed mesoscope TEMPO data. M.Z.L., M.K., G.V. contributed new reagents. R.S. and G.B. contributed analytic tools and neurophysiology expertise. S.H. and M.J.S wrote and edited the manuscript with inputs from all authors. M.J.S. conceived and supervised the project.

## STAR METHODS

### EXPERIMENTAL MODEL AND SUBJECT DETAILS

#### Mice

The Stanford APLAC approved all animal procedures. We used 3- to 12-week-old male and female C57BL/6, Thy1-GFP, PV-Cre, SST-Cre, Cux2-Cre^ERT2^, Nr5a1-Cre and Rbp4-Cre driver lines from Jackson Laboratory (C57BL/6: #000664, Thy1-GFP: #007788, PV-IRES-Cre: #008069, Sst-IRES-Cre: #013044, and FVB-Tg(Nr5a1-cre)2Lowl/J: #006364) and the Allen Institute (Cux2-Cre^ERT2^; MMRRC stock #032779-MU^132^ and Rbp4-Cre KL100; MMRRC stock #037128-UCD). We housed mice in normal light cycle conditions, 2–5 per cage before surgery, and 1 per cage afterward.

#### Transgenic mouse creation

To co-express the FRET-opsin voltage sensor Ace-mNeon1^2^ and the reference fluor mRuby3^84^ in genetically targeted cell-types, we generated a Cre-dependent Ai218 reporter mouse line expressly designed for TEMPO studies (**Figure S6**).^102^ We used our published Flp-in approach^85,87^ and constructed a targeting vector via gene synthesis and standard molecular cloning methods. We used an engineered mouse embryonic stem cell line containing Flp-recombinase sites paired with the constructed TIGRE target vector containing the following component^85–87^: FRT3 – 2X HS4 chicken beta globin insulators – TRE_tight_ promoter – LoxP – ORF-3X stops -Synthetic pA– hGH poly(A), PGK polyA) – LoxP – Ace2N-4AA-mNeonGreen-TS/ER – WPRE-bGH poly(A) – 2X HS4 chicken beta globin insulators – CAG promoter – Lox2272 – ORF-3X stops -Synthetic pA – hGH poly(A), TK poly(A) – Lox2272 – mRuby3– IRES2– tTA2 – WPRE – bGH poly(A) – AttB – PGK promoter – Hygro1-SD.1 – FRT5– origin– Amp (**Figure S6A**). Here, TS/ER denotes the Golgi export trafficking signal (TS) and the endoplasmic reticulum (ER) export sequence, as described.^2^

To express Ace-mNeon1 and mRuby3 in targeted neuron-types, we crossed Ai218 reporter mice with cell-type specific Cre-driver lines. To verify genotypes, we performed the polymerase chain reaction (PCR) on mice tail tissue samples. All mice used for optical experiments were Cre heterozygous and heterozygous Ai218 reporter mice, *i.e.*, *Pvalb-IRES-Cre/wt; Ai218/wt* or Cux2-Cre^ERT2^*/wt; Ai218/wt*. To activate the Cre-recombinase in crosses with the Cux2-Cre^ERT2^ mouse line, we intraperitoneally injected tamoxifen (T5648, Sigma), dissolved in corn oil (C8267, Sigma) at 20 mg/mL, when mice were 5–8 weeks old. Each mouse received 75 mg/kg of tamoxifen daily for 5 consecutive days. Ai218 is available at Jackson Laboratory (https://www.jax.org/strain/037940).

#### Viral vectors

We obtained the following plasmids: pcDNA3-mRuby2^133^ (#40260, Addgene), pNCS-cyOFP^57^ (#74278, Addgene), pAAV-CaMKII-Ace2N-4AA-mNeon^2^ (hereafter termed Ace-mNeon1), pAAV-Syn-VARNAM^8^ (hereafter termed Varnam1; #115554, Addgene), pAAV-CaMKII-Varnam2^1^ (#195527, Addgene), pAAV-CaMKII-Ace-mNeon2^1^ (#195526, Addgene) and pAAV-EF1α-DIO-ASAP3-WPRE^58^ (#132318, Addgene).

We subcloned the genetic sequence encoding the reference fluor, cyOFP, flanked by BamHI and HindIII targeting sequences for the associated restriction enzymes, into an AAV backbone with the CAG promoter followed by the WPRE-hGH PolyA sequence. We used the same strategy for the reference fluor mRuby2. We cloned the genetic sequence encoding the fluorescent GEVI proteins Varnam1, Varnam2 and Ace-mNeon2 into the AAV backbone pAAV-EF1α-DIO-Ace-mNeon1 by inserting the Varnam1, Varnam2 and Ace-mNeon2 protein sequences in place of Ace-mNeon1 between NcoI and AscI targeting sequences, respectively. We cloned the ASAP3 flanked with NheI and AscI into an AAV backbone with CaMKII promoter followed by the WPRE-hGH PolyA sequence. We confirmed that vectors had accurate sequences using Sanger sequencing (Quintara Biosciences). We used the QIAGEN Plasmid Plus Maxi Kit (#12963, QIAGEN) to amplify all plasmids prior to viral packaging.

The Viral Tools facility at the HHMI Janelia Research Campus manufactured all viruses used in our work. For fiber-optic studies (**Figures 1–3**,**S2,S3,S7**), we expressed green GEVIs using AAV2/PHP.eB-EF1α-DIO-ASAP3-WPRE (1.3E13 GC/mL), AAV2/PHP.eB-CaMKII-Ace-mNeon2-WPRE (7.48E13 GC/ml), AAV2/PHP.eB-Ef1a-DIO-Ace-mNeon2-WPRE (2.8E13 GC/ml) and AAV2/PHP.eB-CaMKII-Ace-mNeon1 (3.6E13 GC/mL). We expressed red GEVIs using AAV2/9-CaMKII-Varnam1-WPRE (1.8E13 GC/mL), AAV2/PHP.eB-EF1α-DIO-Varnam1-WPRE (1.3E13 GC/mL) or AAV2/PHP.eB-CaMKII-Varnam2 (3.6E13 GC/mL). We expressed reference fluors via AAV2/PHP.B-CAG-mRuby2 (2E14 GC/mL) and AAV2/PHP.eB-CAG-cyOFP (5.6E13 GC/mL).

For TEMPO microscopy studies (**Figures 4–7**,**S6,S7**), we expressed a green GEVI with AAV2/PHP.eB-EF1α-DIO-ASAP3-WPRE (1.3E13 GC/mL), AAV2-Retro-CaMKII-ASAP3-WPRE (4.6E13 GC/ml), AAV2/PHP.eB-CaMKII-ASAP3-WPRE (4.5E13 GC/ml), and/or a red GEVI with AAV2/PHP.eB-CaMKII-Varnam2 (3.6E13 GC/mL), AAV2/PHP.eB-Ef1a-DIO-Varnam2-WPRE (1.18E13 GC/ml). To express reference fluors, we used AAV2/PHP.B-CAG-mRuby2 (2E14 GC/mL) or AAV2/PHP.eB-CAG-cyOFP (5.6E13 GC/mL). For control studies in which neither fluor was a GEVI (**Figure S7A**), we used AAV2/PHP.eB-CAG-GFP (7.95E13 GC/ml) and AAV2/PHP.B-CAG-mRuby2.

As described previously,^2^ all sequences for red and green FRET-opsin GEVIs, but not for ASAP3, included the endoplasmic reticulum (ER) export sequence and Golgi export trafficking signal (TS), as in the Ai218 mouse reporter line described above.

#### Pharmacology

For studies of ketamine-xylazine-anesthetized mice (**Figures 2,4,S3,S6**), we used ketamine (#50989-996-06, VEDCO), xylazine (#59399-110-20, AnaSed). We dissolved ketamine-xylazine in phosphate buffered saline (PBS; 10 mg/mL ketamine; 1 mg/mL xylazine), yielding final dosages of 100 mg/kg and 10 mg/kg, respectively. For studies of epileptic mice (**Figure 2**), we used kainic acid monohydrate (#K0250-10MG, Sigma-Aldrich). We dissolved kainic acid in NaCl (5 mg/mL), yielding a final dosage of 10 mg/kg. We injected these drugs intraperitoneally (i.p.) with a 30-gauge needle at least 20 min before data acquisition, unless noted otherwise.

### METHOD DETAILS

#### Instrumentation for fiber-optic TEMPO measurements

The uSMAART (**u**ltra-**S**ensitive **M**easurement of **A**ggregated **A**ctivity in **R**estricted cell-**T**ypes) fiber photometry system has 4 main modules: the (1) low-noise illumination module; (2) decoherence module; (3) low-noise fluorescence sensing module; and (4) phase-sensitive demodulation module.

##### Low-noise laser illumination module

Our laser illumination module is ∼10-fold more stable than that of prior photometry systems (**Figure S1E**).^7,9,54,55^ To achieve stable illumination (<0.005% r.m.s. noise fluctuation), we selected solid-state laser sources, instead of light-emitting diodes (LEDs), since the former generally have ∼10-fold lower noise levels than the latter (**Figure S1A-B**). Further, commercial solid-state lasers commonly have an analog modulation bandwidth that extends to the ∼100 kHz range, much beyond the modulation bandwidths of typical LEDs. We modulated our lasers at 50 or 75 kHz, to perform measurements in a part of the frequency spectrum where both lasers and detectors have reduced instrumentation (1/*f*) noise (**Figure S1C**). We used a purely sinusoidal analogue modulation scheme to ensure that no spectral harmonics propagated to adjacent recording channels.

To attain low-noise illumination from a laser source, it is important to minimize optical back-reflections into the laser cavity, which can lead to unstable emission output levels. To prevent back-reflections, we placed a Faraday isolator (–40 dB attenuation) immediately succeeding each laser source in its illumination pathway (**Figure 1A**). Given these considerations, we used continuous wave laser light sources with 488 nm and 561 nm wavelengths (488-20LS and 561-50LS OBIS Lasers, Coherent) to excite green and red fluorescence, respectively. We used a fixed free-space Faraday isolator (IO-3-488-HP, Thorlabs) and a tunable free-space isolator (FI-500/820-5SV, Linos) to prevent back-reflections into the two respective laser cavities. To further reduce back-reflections, we coupled all laser beams into an 8°-angled fiber-optic patch-cord (FC/APC patch cord in **Figure 1A**).

To align the polarization of the 561-nm-emissions with the polarization axis of the accompanying isolator, we aligned an achromatic half-wave plate (AHWP05M-600, Thorlabs) in front of the isolator’s entrance face. We aligned the two laser beams onto a common optical path using steering mirrors (BB1-E02, Thorlabs) and iris diaphragms (CP20D, Throlabs). To direct 10% of the light from each laser beam to a photodiode (PDA100A, Thorlabs) for continuous monitoring of the emission power, we used a pair of 90/10 beamsplitters (BS028, Thorlabs). We combined the other 90% of the light power from each of the two beams using a dichroic mirror (FF511-Di01, Semrock). We coupled the pair of collimated beams into a multi-mode, 8°-angled fiber (105 µm core, 0.2 NA; FG105LCA, Thorlabs) with an FC/APC fiber collimator (F240APC-532, Thorlabs). Overall, the r.m.s. level of intensity fluctuations of the blue illumination was ∼0.004%, much lower than the nominal 0.25% r.m.s. value of fluctuations in the illumination intensity output from the OBIS lasers.

##### Decoherence module

The decoherence module breaks the coherence between light in different spatial (transverse) modes within the optical fiber (**Figure 1A**), rendering the illumination pattern and output power insensitive to motion of the multimode fiber connected to the animal. Without this coherence reduction, fiber motion would induce optical mode-hopping fluctuations, a form of optical speckle (**Figure S1D-E**).^48^ To break the illumination coherence, we built a dual-stage optical diffuser, the second stage of which is dynamic and reduces the spatial coherence of the beam by inducing rapid variations of the optical path length across each beam’s spatial cross-section. Within the decoherence module, after exiting the FC/APC optical fiber, each laser beam passes through an axicon lens, which converts each beam’s Gaussian cross-sectional profile to an approximate flat-top intensity profile (**Figure 1A**). This increases the number of micron-scale grains within the diffuser that interact with and scatter the laser light.

To implement this approach, we first collimated the laser light exiting the FC/APC fiber module using a fiber-optic collimator (F240APC-532; Thorlabs). Next, we positioned onto the light path a Ø1” plano-convex ring grade axicon lens (140° apex angle; #83-790, Edmund optics). The axicon focused each beam onto the dual static-dynamic diffuser (LSR-3005-24D-VIS, Optotune), which had an average grain size of 3 μm. The second diffuser in the pair circularly oscillated at 300 Hz in the two lateral dimensions orthogonal to the longitudinal optical axis, actuated by four independent electro-active polymer electrodes. To collimate the diffused laser light, we used an aspheric lens (A240TM-A, f = 8.00 mm, NA = 0.5, Thorlabs). Finally, to maximize the photon collection efficiency, we coupled the beams into a FC/PC multimode fiber Ø1000 µm / 0.39 NA (FP1000URT, Thorlabs) using a fiber collimator (F240PC-532, Thorlabs). To enable concurrent recordings in two brain areas (**Figure S3**), we coupled the illumination into a FC/PC bifurcated multimode fiber bundle, Ø400 µm / 0.39 NA, (BFY400HF2, Thorlabs), using a FC/PC to FC/PC mating sleeve (ADAFC2, Thorlabs).

In sum, owing to the decoherence module, the noise reductions achieved in the low-noise illumination module (r.m.s. fluctuations of ∼0.004% at the specimen plane) were impervious to fiber and animal motion (**Figure S1D,E**).

##### Fluorescence collection module

The collection module maximizes fluorescence signals while minimizing noise (**Figure 1A**). We computed, for multiple commercial filter sets, the expected collection efficiencies and levels of crosstalk between fluorescence detection channels, based on the spectral properties of all fluorophores used in our experiments. We thereby selected filter sets for which bleedthrough of emissions from the reference fluor into the detection channel for the GEVI was <0.01%, minimizing the addition of photon shot noise from reference fluor emissions into neural voltage measurements.

To maximize signal-to-noise levels of fluorescence detection, we selected an avalanche photodetector with a noise-equivalent power of 2.5 fW/√Hz and a signal detection bandwidth (100 kHz) wide enough to accommodate the two lasers’ analogue modulation frequencies (50 kHz and 75 kHz).

To deliver light to and from the brain, we chose an optical fiber of low intrinsic autofluorescence. Minimizing autofluorescence is vital, as otherwise it will diminish a detector’s effective dynamic range; further, autofluorescence photon shot noise can mask the weak signals associated with high-frequency voltage dynamics. To further minimize fiber autofluorescence, we photobleached the fiber conveying light to and from the brain at least 1 day before each experiment using the same illumination wavelengths, yielding a ∼5-10-fold reduction in autofluorescence (∼1–10 pW residual fiber fluorescence induced by ∼100 µW of 488 nm illumination).^134^ We removed residual light in the cladding modes of the fiber delivering light to and from the brain by winding the fiber around a 10x mandrel wrap (**Figure 1A**).

In our specific implementation of these approaches, the fluorescence collection pathway had two spectral detection channels, and a single-band dichroic mirror and two bandpass filters to separate and steer green and red signals to separate photodetectors. The excitation laser beams from one branch of the bifurcated multimode fiber bundle were collimated using a fiber collimator (F240PC-532, Thorlabs), reflected off a dual-edge dichroic mirror (Di01-R488/561, Semrock), and focused into a multimode, low autofluorescence, pre-photobleached, fiber-optic patch cord (1.5-m-long, Ø400 µm-core, 0.50 NA; FT400URT, Thorlabs) using a fiber collimator (F240PC-532, Thorlabs). The patch cord fiber was tightly wrapped ten times around a Ø1” pedestal post (RS3P8E, Thorlabs), placed just after the fiber collimator, which served as mandrel wrap and provided about a 3 dB (or ∼50%) reduction in the illumination fluctuations at the specimen when the optical fiber is moving owing to movement of the animal. Using a ceramic mating sleeve (ADAF1, Thorlabs), we connected the patch cord to an optical fiber (CFM14L05, Thorlabs) that we implanted in the mouse brain (cf. **Mouse Preparation**). The total mean power delivered to the brain was 25–200 µW. Fluorescence emissions from the brain passed through the dual-edge dichroic mirror and were split into red and green channels using a single-edge dichroic mirror (FF562-Di01, Semrock). Light in each channel passed through a bandpass filter (FF01-520/28 or FF01-630/92, Semrock) and was focused onto an avalanche photodiode (APD) (APD440A2, Thorlabs) by an aspheric lens (A240TM-A, f=8.00 mm, NA=0.5, Thorlabs).

##### Phase-sensitive detection module

The phase-sensitive detection module unmixes emission signals that are excited by different lasers but captured on the same detector. With this approach, one need not rely solely on emission filters to separate fluorescence from different fluors but can also exploit the fluors’ different absorption spectra. To demodulate the fluorescence signals from the photodetector outputs, we used a pair of low noise, digital lock-in amplifiers (LIA) (MFLI-MD, Zurich Instruments) (**Figure 1A**), one for each illumination path. Each LIA provides one analog modulation signal and can demodulate up to 4 signals.

We amplitude-modulated the 488 nm and 561 nm lasers with analogue sinusoidal oscillations at 75 kHz and 50 kHz respectively, with 0 V to 4 V peak-to-valley amplitudes. We used one LIA to demodulate signals from the photodiode measuring 488 nm laser power fluctuations and from the APD measuring green fluorescence signals. We used the other LIA to demodulate signals from the photodiode measuring 561 nm laser power fluctuations and from the APD detecting the red fluorescence. Signals were demodulated using a linear-phase finite impulse response (FIR) filter, which is specifically designed for digital signal processing applications and allows low-pass filtering of signals with a frequency range that depends on the demodulation frequency. The low-pass filter used for demodulation was set to have a 0.1 ms time-constant and 48 dB/octave roll-off, enabling a measurement bandwidth of 0–500 Hz.

For concurrent fiber-optic TEMPO recordings from two cell types (**Figures 3,S2,S3**), we routed the reference modulation signal from the first LIA controlling the 488-nm-laser to the second LIA, in order to demodulate the signal from the APD capturing red fluorescence and thereby to extract the time-varying level of cyOFP fluorescence.

For concurrent measurements from two cell-types and two brain regions (**Figure S3**), we duplicated the fluorescence collection module. This approach yielded 4 independent APD signals, 2 per brain area. The LIA configuration was similar to that used for studies with a single GEVI, except that we no longer measured laser power fluctuations, thereby enabling each of the two LIAs to demodulate all 3 fluorescence signals collected by a single optical fiber.

By combining the set of 2 emission filters and 2 dichroic mirrors with the frequency modulation and demodulation scheme used in our phase-sensitive detection module, we obtained the following optical bleedthrough and cross-talk numbers for the uSMAART systems. Less than 0.001% of the collected fluorescence from the red fluor (either mRuby2 or Varnam2) enters the green fluorescence detection channel. Approximately 8.8% of the fluorescence collected from the green fluor ASAP3 (or ∼6.4% for Ace-mNeon1) enters the red fluorescence detection channel. However, owing to the frequency modulation, the red and green signals can be unmixed from each other, because they are excited by lasers modulated at distinct temporal frequencies. Therefore, virtually no green fluorescence (either GFP, Ace-mNeon1 or ASAP3) contributes to the demodulated signal from the red fluorescence detection channel, except for the added green fluorescence photon shot noise, which has a white frequency spectrum and cannot be removed through demodulation.

Nevertheless, in a dual-GEVI experiment, because cyOFP is excited with the same laser source as the green fluor, the bleedthrough of green GEVI fluorescence into the red detection channel does affect the demodulated cyOFP signal. Further, 1.1% of the fluorescence collected from cyOFP bleeds into the green fluorescence detection channel. In addition, because cyOFP has a broad absorption spectrum, a certain amount of cyOFP fluorescence can be excited by the green laser used to excite Varnam. Consequently, when the green and blue laser intensities are set to equivalent levels at the specimen, ∼18.5% of the total cyOFP fluorescence collected in the red fluorescence channel cannot be removed through frequency demodulation and hence will appear as crosstalk in the demodulated Varnam2 signal. Conversely, ∼10.3% of the collected fluorescence from Varnam2 will appear as crosstalk in the demodulated cyOFP signal, because the blue laser excites both fluors. For these reasons, in dual-GEVI experiments, data analyses included a decrosstalking step (see below, **Data processing for fiber-optic TEMPO recordings**)

#### Electrophysiological recordings

We performed electrical recordings concurrently with fiber-optic TEMPO (**Figures 1–3**, **S2,S3,S7**) and TEMPO imaging studies (**Figure 6**). We amplified electrophysiological signals with a digital head-stage amplifier (RHD 16-channel or 32-channel, Part #C3334 or #C3324, Intantech) attached to our custom-made optrode (see below). We acquired digital data at a 2-kHz-sampling rate using a USB data acquisition board (RHD USB interface board, #C3100, Intantech).

#### Synchronization of optical and electrical recordings

To synchronize the optical and electrical recording instruments, we generated a synchronizing digital signal (hereafter called ‘syncTTL’), which comprised a square-wave with a 50% duty ratio and a 1-s-long ON-state but with intervals of random duration between the square pulses, to enable unambiguous temporal alignments. The syncTTL signal was externally and independently generated by a data acquisition system (USB-6003, National Instruments) and recorded concurrently by both the electrical and optical data acquisition systems.

To temporally align the optical and electrical signals, we first preprocessed the LFP signal as follows: (1) we removed any residual 60-Hz narrow-band noise, representing contamination from the electrical power line, by using an 8^th^-order Butterworth notch-filter; we designed the filter using the *designfilt()* function in MATLAB, with the option ‘bandstopiir’ and the low and high cut-off frequencies set at 59.5 Hz and 60.5 Hz, respectively. We then applied the filter using the zero-phase digital filtering function *filtfilt()*; (2) We bandpass-filtered the resulting LFP traces with a 3^rd^-order Butterworth filter with low and high 3 dB cutoff frequencies of 0.1 Hz and 500 Hz; (3) we interpolated the LFP traces and the squared syncTTL signal so the sampling rates of the resulting traces matched the sampling frequency of the optical data. We used the MATLAB function *interp1()* with the ‘spline’ method to interpolate the continuously valued LFP trace. To interpolate the binary valued syncTTL signal, we used the *interp1()* function with the ‘nearest’ method. Finally, we aligned the optical and electrical traces using the temporal registration provided by the concurrently recorded syncTTL signal.

#### Optical and optrode implants

For all uSMAART studies with a single GEVI, we implanted an optrode into the brain to allow joint optical and LFP measurements (**Figures 1, 2, S2**), except for our recordings in the visual cortex of anesthetized mice (**Figure 2A–D**), for which we only used an optical fiber. For TEMPO microscopy studies of hippocampus, we used a distinct but similar optrode for dual optical and electrical measurements (**Figure 6**). For uSMAART studies of visual cortex with two GEVIs (**Figure 3, S2**), we implanted an optical cannula without an ipsilateral LFP electrode, except for **Figure 2G-I**. However, for studies of hippocampus with two GEVIs, we implanted a linear silicon probe in the contralateral hippocampus. We manufactured all optical and optrode implants under a stereomicroscope (Leica MZ7-5) equipped with a soldering station.

For TEMPO mesoscope studies in the hippocampus (**Figure 6**), we manually fabricated each cannula implant using a borosilicate cover glass (170 ± 5 µm thickness, Schott) that was laser cut to a 3-mm-diameter circle (Fluence Technology). We glued the cover glass onto a 3-mm-outer-diameter, 1.5-mm-long stainless-steel ring (grade 316, gauge 11TW, MicroGroup) using adhesive cured with ultraviolet light (Loctite #3105). To record optical and electrical signals simultaneously, we glued two tungsten wires (0.002”, 99.95% CS, with a single polyimide insulation, M215580, California Fine Wire) to the optical implant, ∼50 µm beyond the glass window, using UV-light-cured adhesive (Loctite #3905). We used coated stainless-steel wire (0.005” bare, 0.008” coated, 100 feet, A-M systems) for the ground electrode, and we soldered both the LFP and ground electrodes into a gold-plated pin head (0.031”, WPI). During the implant surgery, we positioned the ground electrode in the cerebellum.

Optrode implants for uSMAART comprised an optical fiber (400 µm core diameter, 0.5 NA, Thorlabs) (**Figures 1–3,S2,S3**). Optrode implants for hippocampal imaging comprised a circular borosilicate cover glass (3 mm diameter, 170 mm ± 5 mm thickness, Schott) that was glued to the end of a stainless-steel ring (3 mm OD, 1.5 mm long) with UV-curable adhesive (Loctite #3105) (**Figure 6**). To achieve dual-modality, optical/LFP recordings (**Figures 1–3,6**), we affixed a pair of ∼1-cm-long, ∼50-µm-diameter tungsten wires coated with single polyimide insulation (M215580, California Fine Wire) to the optical fiber (**Figures 1–3**) or cannula (**Figure 6**) at two different axial positions, spaced in increments of ∼50 µm, with the first tungsten wire placed flush with the optical fiber or window, and with the tip of the other wire extended into the brain tissue under investigation. This set of electrode placements was chosen to maximize the chances that at least one of the electrodes would provide high quality LFP signals, given that our surgical placement of the implant was performed without fine visual feedback regarding the exact location of the ∼100-µm thick pyramidal cell layer of CA1. We glued these tungsten wires onto the optical cannula with ultraviolet-light-cured adhesive, which constrained the electrodes to be laterally offset by 1.5 mm from the center of the imaging window. Next, we soldered a gold-plated pin-head (NC0102069, WPI) onto the end of each tungsten wire projecting away from the imaging window. The reference electrode comprised a ∼5-mm-long, ∼127-µm-diameter, uncoated, stainless steel wire (#791600, A-M Systems) soldered to a gold-plated pinhead (NC0102069, WPI). Separately, we fabricated a 4-channel-connector using 4 gold-plated sockets (NC1456862, WPI) and a connector board (EIB8, Neuralynx).

#### Mouse preparation for fiber-optic TEMPO measurements

Mice (aged 8–16 weeks at start) underwent two surgical procedures under isoflurane anesthesia (1.5%–2% in O_2_). In the first procedure, we injected viruses to express the GEVI and reference fluor. In the second procedure, performed about a week after viral injection, we implanted the optrode.

Coordinates for virus injections in CA1 (**Figures 1–3, 6, S1–S3, S7**) were –2.0, 1.7, –1.1 (in mm from Bregma; AP, ML, DV). Coordinates for virus injections in V1 (**Figures 2-3, S2, S3, S7**) were –3.5, 2.5, –1.2. Coordinates for injections of retrograde virus in the lateral septum (**Figures 6**) were 0.4, 0.4, –3.3. For single cell-type studies, we injected 0.5 µL per mouse of a solution containing 5E9 genome copies (GC) of the GEVI virus and 1E9 GC of the reference virus. For dual cell-type studies, we injected 0.5 µL per mouse of a solution containing 10E9 GC of the Cre-dependent GEVI virus, 5E9 GC of the CaMKII-dependent GEVI virus and 1E9 GC of the reference virus.

For all mice, we prepared the skull by manually removing the conjunctive tissue. We cleaned the skull by applying H_2_O_2_, quickly followed by rinsing with Ringer’s solution (#50980245, ThermoFisher). For studies in the primary visual cortex V1, we drilled a 0.5-mm-diameter opening in the skull above V1 using 0.5-mm-diameter micro drill burr (#19007-05, Fine Science Tools). For studies in the dorsal CA1, we drilled a 2-mm-diameter opening in the skull above CA1 using 0.5-mm-diameter micro drill burr (#19007-05, Fine Science Tools). We removed cortical tissue above CA1 using suction applied via a 30-gauge blunt needle under ice cold Ringer’s solution. To improve optical access to hippocampal tissue, we gently removed two of the three layers of the overlying alveus. We discarded mice in which CA1 was damaged, as a damaged hippocampus is prone to undergo local epileptic seizures. We inserted the implant and sealed the gap between the skull and the implant using adhesive cured with ultraviolet light (Loctite 33105). In mice used for uSMAART studies under head-restraint (**Figures 3, S2, S3**), we glued a custom-designed stainless steel head bar on the mouse skull and secured the implant to the skull using blue-light cured resin (Flow-it ALC, Pentron). To mitigate post-operative pain, we administered carprofen (5 mg kg^−1^) about 30 min prior to the end of surgery. Mice recovered for >2 weeks before imaging experiments began.

#### Fiber-optic TEMPO recording sessions

We performed all recordings during the mouse’s light cycle. In studies with ketamine-xylazine-anesthetized animals, mice were either unrestrained (**Figure 2**) or head-restrained (**Figure S3**). Recordings began ∼20 min after anesthesia administration, when the mice became immobile, and lasted 2–10 min. In studies of mice allowed to behave freely or those with kainate-induced epilepsy, the animals were placed in a circular arena with clear acrylic walls (31 cm diameter, 31 cm height, **Figure 1**) or within their house cage (**Figures 2, 3**); the recordings lasted <60 min. In some experiments (**Figures 2, 3**), we took video recordings of the mouse’s behavior in the open field arena or home cage using infrared illumination and an overhead camera (DMK23 U40, Imaging Source) that acquired images at a 20 Hz frame rate. To control video acquisition, we used the IC-Capture software package (Imaging Source).

#### Visual stimulus presentation

We placed an LCD monitor (85-cm-diagonal) in the monocular visual field of the mouse at a distance of 20 cm, contralateral to the fiber optic implant (**Figures 3, S2, S3**) or cranial window (**Figures 5, 7, S6, S7**). We used custom software developed with the Psychtoolbox in MATLAB (MathWorks) to display drifting gratings (sine waves of 1 Hz temporal frequency, 0.033 cycle per degree of spatial frequency and 100% contrast, oriented at 315° to horizontal). We placed a blue bandpass optical filter (#5084 Damson Violet - 48’ X 25’ Roll, Rosco E-Colour) onto the video monitor to ensure that its illumination was outside the range of wavelengths detected by the TEMPO apparatus. Presentations of the drifting grating lasted 1.5 s and were preceded and followed by presentations of a gray isoluminant screen. Inter-trial intervals (ITIs) were randomized between 2–5 s.

#### Behavioral studies of rest-to-run transitions

We built a custom mouse running wheel using a 3D-printed PLA plastic wheel (8 cm wide, 13 cm diameter). We covered the wheel’s surface with self-adherent wrap (Coban, 3M) and tracked the angular displacement using a rotary encoder (optical AB Phase Quadrature Encoder 600P/R, Amazon). We recorded the wheel displacements via a data acquisition device (BNC-2090A, National Instruments) controlled via LabView 2019 software (National Instruments). To elicit a rest/run behavioral transitions, we delivered an airpuff to the mouse’s back. The airpuff intensity was ∼30 psi and was administered via a 20-gauge blunt needle placed 1–2 cm above the mouse’s back. We triggered the air puff after the mouse stayed immobile for at least 5 s. Each experimental session lasted for about 5 min. All mice were habituated on the running wheel for at least 2 days, for at least 15 min per day, before TEMPO recordings began (**Figures 6, S2**).

#### TEMPO mesoscope

The mesoscope system includes an LED illumination module, an objective lens with an electronically controlled focusing mechanism, a fluorescence filter module, a tube lens system that allows differential focusing between the two fluorescence channels, a pair of sCMOS cameras, data acquisition and system control electronics, as well as a custom mechanical design (**Figure 4**).

We developed the TEMPO mesoscope to monitor neural voltage dynamics across wide fields-of-view with high spatiotemporal resolution and detection sensitivity. (Our system has a nominal field-of-view of 8 mm, although, in practice, we used glass cranial windows of 7–8 mm in diameter). The imaging system we designed and built has several distinctive advantages, including a 0.47 numerical aperture (NA) objective lens, which enables good fluorescence collection across the large field-of-view and provides high efficiency (∼85%) transmission of visible wavelengths. The system’s high-speed imaging capabilities allow recordings at 100 Hz across the full field-of-view and even faster recordings over a reduced field-of-view of width, *FOV*_*y*_, at a frame rate of 100 *Hz* × 8 *mm*/*FOV*_*y*_. For example, to study gamma activity we typically acquired images at 130–300 Hz; the images taken at ∼130 Hz were of sufficient spatial size to cover the full area of visible brain tissue through a ∼7-mm-diameter cranial window (*e.g.*, **Figure 4H**), whereas we used ∼300 Hz frame acquisition to capture in detail the propagation of individual gamma waves (*e.g.*, **Figure 4O**). The optical apparatus has a resolution of approximately 6 μm and two fluorescence detection channels that use a custom-made, polished dichroic filter (λ/4 peak-to-valley wavefront errors, Alluxa) to separate green and red fluorescence. This combination of features is unavailable in commercial or previously published microscopes, yet was vital for the unprecedented recordings of this paper.

Past work on large-scale voltage imaging focused on slow (<10 Hz) voltage activity and used imaging systems that supported the detection of voltage activity in this frequency range.^135,136^ During the optical design process, we sought a design that maximized the SNR, imaging frame-rate and field-of-view for studies of voltage dynamics up to the high-gamma range. We also sought recordings up to 60 min, without photodamage to tissue or undue photobleaching. To obtain SNR values sufficient to detect the tiny fluorescence changes associated with fast voltage dynamics (*i.e.*, <0.1% ΔF/F), we sought photon shot-noise-limited measurements such that the mean flux of signal photons would be much greater than accompanying photon shot noise. In practice, the goal of maximizing the photon flux implies that the cameras are operating close to saturation.

Given this general approach, we examined the SNR for detecting a voltage wave of spatial wavelength Λ_γ_ and temporal frequency *f*_γ_. For simplicity, we considered a wave propagating in one spatial dimension, *A*(*x*, *t*) = *A*(φ(*x*, *t*)), where the phase depends on space and time, φ(*x*, *t*) = 2π(*x*/Λ_γ_ − *t*/*f*_γ_). We considered the SNR level needed to detect the wave in a single trial (*i.e.*, without trial averaging) and quantify its propagation properties. Assuming there is a spatially uniform noise of standard deviation, σ_ɛ_, originating mainly from photon shot noise, the ability to distinguish changes in the wave’s amplitude in space and time via two measurements is related to the local phases of the wave and to the desired SNR:

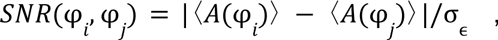

where φ*_i_* and φ*_j_* are the phases of the wave for the two measurements in pixels *i* and *j*. Similarly, the ability to detect the wave relates to the SNR with which one can distinguish between the polar-opposite phases:

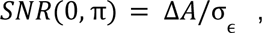

where Δ*A* = |〈*A*(0)〉 − 〈*A*(π)〉|. For waves at the edge of detectability, it is beneficial to work at the Nyquist limit of the wave and to boost the SNR by averaging fluorescence signals across as many pixels as possible within a correlation length, which can be estimated as half a wavelength, Λ_γ_ /2.

(Averaging over pixels beyond this correlation length would lead to canceling out of the wave amplitudes). The number of photoelectrons acquired from this width, Λ_γ_/2, of tissue, within *n*_*f*_ image frame acquisitions relates to the corresponding number of camera pixels and their individual well capacity, *W*_*C*_, assuming the camera is operating close to saturation. Given a system magnification, *M*, and pixel pitch, *l*_*px*_, the maximum number of photoelectrons that can be acquired is thus:

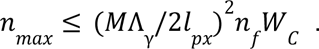

This sets the shot noise in the photoelectron count as 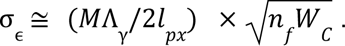. For a wave of maximum fluorescence amplitude 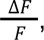 the maximum SNR thus is bounded as follows:

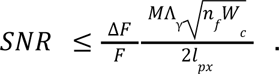

Optimization of this expression leads to high magnifications and the use of many pixels of large well capacity, which is opposite to the approach taken in most prior voltage-imaging studies in which the dorsal neocortex was imaged at low magnification in order to obtain fast imaging frame rates.^136^ Further, with our sCMOS cameras, one can use a high sampling frequency, *f,* to maximize *n*_*f*_, at the expense of the vertical field-of-view, *FOV*_*y*_. Similarly, the magnification limits the field-of-view, suggesting that major gains in SNR are attainable in exchange for working with limited fields-of-view and large pixel counts of large well capacity. (**Figure S4H-J** has plots showing how the SNR scales with magnification and pixel well capacity at a constant illumination power and field-of-view).

Given the above considerations, we decided to build a microscope whose imaging field can span our 7-8 mm cranial window preparation. Given this field-of-view size, we sought to optimize the other properties noted above that influence detection sensitivity, and we chose fast and high-quantum efficiency sCMOS cameras with sensor chips 13–15 mm across. We conducted optical design studies with ZEMAX software. We modeled the objective and tube lenses in an infinity-corrected configuration and aimed to optimize the resolution and transmission across the field-of-view, which led to the minimization of vignetting and aberrations while keeping photon transmission high.

During optical design, we examined the properties of both custom lenses and commercially available optical components. After an iterative optimization process that considered both the properties of the objective lens and its performance in conjunction with the tube lens, we selected two existing lenses for high-end photography. The Leica Noctilux-M 50mm f/0.95 ASPH (NA=0.47) serves as our system’s objective lens, and the Canon EF 85mm f/1.2L II USM serves as the tube lens in each spectral channel. This pair of lenses provides a net optical magnification of 1.85x, a resolution of about 6 µm, and visible light transmission of ∼85% at 532 nm in the absence of any filters.

To accompany the objective and tube lenses, we designed an illumination module to provide high-power, high-stability incoherent illumination across the absorption spectrum of our green and red fluorescent GEVIs. Illumination from two LEDs (UHP-T-475-SR and UHP-T-545-SR both with low-noise and detached-fan options, Prizmatix) was combined using a dichroic beam combiner (T505LPXR, Chroma) installed in a beam combiner module (Prizmatix). The light then passed through a dual-bandpass clean-up filter (FF01-482_563, Semrock). A dual-band custom dichroic (488/568-Di, Alluxa) reflected illumination from both LEDs and transmitted the fluorescence. The LEDs had a specified emission stability of <0.05%, and in dual cell-type experiments we directly monitored the illumination power using photodiodes.

A motorized stage (Sutter MP-285) allowed us to perform fine axial focus adjustments of the objective lens. We created a custom mechanical assembly to mate the lens with the focusing stage and to allow a high mechanical accuracy of insertion and alignment with the rest of the optical axis. An additional differential focusing mechanism relied on the internal focusing mechanism of the Canon 85 mm f/1.2 tube lenses, controlled by an Ultrasonic Motor (USM) Autofocus (AF) controller (Astromechanics) via the serial port. Each USM focus controller had a EOS-C-mount adapter that allowed alignment between the lens and the camera. Alignment of the tube and objective lenses for the green channel was achieved using a CNC-machined filter cube holder (**Figure 4**); residual angular misalignment between the two video streams was corrected computationally. Fine alignment of the red-channel tube lens/camera module was implemented using a 5-axis compact kinematic stage (Newport 9081-M), placed at the base of the camera.

The microscope’s emission pathway was designed to separate the illumination from the fluorescence emissions in both spectral channels, while preserving the optical resolution in both transmission and reflection. In the emission path, we used a custom-made 550 nm short-pass dichroic mirror (Alluxa), placed on a super-flat substrate, such that the wavefront error in reflection was empirically 150 nm or less across the entire coated filter. We used emission filters specific to each color channel, a 520/41 nm for Ace-mNeon (Alluxa) and ASAP3 and 609/62 nm (FF01-609/62-25, Semrock) for VARNAM2. With these filters, only 0.1% of the fluorescence collected from the red fluor (either mRuby or Varnam) enters the green fluorescence detection channel. Whereas ∼7% of the fluorescence collected from green fluorescent ASAP3 (∼9.5% for Ace-mNeon1) enters the red fluorescence detection channel. In experiments using cyOFP, ∼2% of its collected fluorescence enters the green fluorescence detection channel. During data processing, we computationally corrected for the bleedthrough of the green emissions into the red fluorescence detection channel (see section below, *Processing of TEMPO mesoscope video data*).

We utilized scientific grade sCMOS cameras (Hamamatsu ORCA Fusion Digital CMOS camera; Hamamatsu Photonics K.K., C14440-20UP) as detectors. We controlled two cameras through the CoaXPress interface using a PCIe frame grabber and streamed the data through the same interface.

After having designed the optics, we designed the mechanical assembly using Inventor (Autodesk) computer-aided-design (CAD) software. The main components were custom-designed and machined on a 5-axis CNC mill (Optima Precision) using aluminum alloys 6060 and 7075 and then anodized black to reduce stray light from reflections.

Electronics for controlling the microscope and data acquisition are detailed in **Figure S4**. The cameras ran in external trigger mode, using the configuration of **Figure S4C**. For studies using a single GEVI, we used the External Start Trigger mode to initiate frame acquisition on both cameras. The cameras acquired frames in the overlap rolling shutter mode based on the internal pixel clock, which we verified gave temporal synchronization between the two cameras across recordings of up to 30 min. (We did not test longer time spans). This operation mode provided stable exposure durations as well as the fast data acquisition rates that our work required.

Figure S4 provides a schematic of the camera operations, including the trigger output signals used for diagnostics and control, such as VSYNC or Global Exposure signals. As **Figure S4** shows, we controlled the system from a PC workstation utilizing PCIe and serial port interfaces. A DAQ board (myRIO-1900, National Instruments) equipped with a custom-programmed field programmable gate array (FPGA) (Xilinx Z-7010) controlled all analog and digital signals. The DAQ board providing the interface to the FPGA was controlled by the PC operating the mesoscope apparatus. Live data from each camera streamed to a RAID0 array of NVMe SSD (Samsung 970 Pro) hard drives on a NVMe RAID Controller with a PCIe interface (Highpoint Technologies Inc).

We stored data files as native Digital Camera Image files (DCIMG, Hamamatsu). We used Samsung NVMe 970 Pro M.2 storage with triple-cell base units and no buffering, which was crucial for achieving sustained write-in using consumer grade hard drives. For long-term storage of raw data, we used a QNAP NAS TS-1685, which hosts a RAID6 array of 140 TB of storage, with 2TB of NVMe (Samsung 860) cache, connected to the acquisition computer via a direct 10 Gbps optical link.

In practice, we found that a suboptimal streaming configuration, even when using hard-drive interfaces that nominally exceed the requisite write-in speed, may lead to detrimental frame dropping. To prevent frame-dropping, we established a diagnostic routine based on the analysis of the time stamps natively written in the DCIMG files and decoded with custom software provided to us by Hamamatsu. The precision of the time stamps was in the microseconds range, allowing us to detect frames whose exposure interval was notably larger than the nominal period between image frames. With this approach, we verified several hard-drive configurations that provided a stable data streaming architecture for >1 yr. We observed that after ∼1 year of use, cells in NVMe memory can degrade, leading to a recurrence of frame dropping. This issue can be resolved by replacing the hard drives.

#### TEMPO imaging studies in two neuron-types

To track the emissions of 3 different fluorophores, we created a protocol based on the interleaving of illumination from the two different LEDs and our use of two spectrally distinct GEVIs plus a long Stokes-shift reference channel (cyOFP) (**Figure 7**). We operated the two LEDs in pulsed mode, interleaving them in time, while the two cameras recorded continuously as we encoded information about which fluorophore(s) were being excited and imaged (**Figure S4**). Successful implementation of this protocol necessitated synchronizing the two cameras, operating in rolling shutter mode, with the pulsed light sources. To prevent spatially varying artifacts caused by uneven illumination of camera rows, the synchronization had to be precise down to the readout time (10 μs) of a single image row.

To implement the illumination interleaving scheme, we programmed the FPGA (40 MHz clock) to create trigger pulses for the LEDs, and we synchronized them to the global-exposure trigger from both cameras that marks the time period when all rows of the camera are simultaneously acquiring photons (**Figure S4**). Synchronization to the global exposure allowed us to avoid spatial artifacts that might have arisen due to temporal mismatches between two cameras operating in rolling shutter mode. The FPGA created digital pulses that externally gated the LEDs. The FPGA also detected, counted, and, depending on the parity of the count number, rerouted global exposure signals to the LEDs (**Figure S4**). We used the triggering protocol shown in **Figure S4E** for dual GEVI studies, in which exposure times were equalized across different rows of the camera yet remained longer than the nominal, fastest possible acquisition time given the number of rows being read out. For instance, the acquisition of 2048 rows at 100 Hz led to negligible, 10 μs temporal overlap between the exposures of the first and last rows. An increase of the exposure time to 12.5 ms decreased the frame rate to 80 Hz yet allowed us to use 2.5-ms-pulses of light that were precisely synchronized to the time period when all camera rows were exposed simultaneously.

The protocol outlined above was effective in cases when the fluorophores were expressed in a balanced manner, as shown in the results presented in **Figure 7**. However, when expression of cyOFP was stronger than that of the GEVIs, we adjusted the exposure time for the camera in the red fluorescence camera to reduce the amount of red fluorescence excited by the longer-duration blue pulse, which was also used to excite the green GEVI. To capture signals from 3 fluorophores with varying expression patterns and intensities, cameras with different exposure times are needed in principle. However, the use of multiple illumination sources can result in extra intensity fluctuations, and running two cameras in frame-by-frame trigger mode can introduce exposure and illumination jitters that degrade the accuracy of the data. Therefore, for studies with 2 GEVIs, we implemented a protocol using the External Edge Trigger mode and trigger signals generated by the FPGA clock, which allowed the two cameras to operate with different exposure durations.

#### Software for TEMPO imaging

We controlled the TEMPO mesoscope using custom software written in LabView (2019) (National Instruments). The high-level architecture of our software has 3 main components (**Figure S4F**). The main software, Recorder, was the primary controller of most of the hardware components and served as the primary user interface. To create recorder interfaces to the camera, LEDs and mechanical actuators, we respectively used a software development kit (SDK) from Hamamatsu, and serial port communication protocols from Prizmatix and Sutter. Recorder exchanges the software state and metadata with subsequent FPGA programs, split into Host and Target programs. The diagram in **Figure S4F** shows all software submodules and the relationships between them, as well as with the external devices, lines or file containers.

The submodules of Recorder, running on the PC, handle several functions. The camera setting manager interfaces with the sCMOS cameras, while the DCAM streamer interfaces with a set of proprietary DLL libraries from Hamamatsu. The software includes a file system manager and a metadata manager to store and organize data that we designed and implemented to support our experiments and data analysis pipeline. A shared setting server allows multiple users to access the same settings. Image frames from the cameras are buffered for real-time monitoring, and the software runs simple analytics to check data quality. The Recorder software also enables active control of the LEDs and focusing stage, which are synchronized to the recordings. The LEDs are turned on only during the recording and are pre-launched 0–10 s before recording starts, to remove the fastest photobleaching components from the fluorophore signals.

The FPGA host software includes a shared settings server that links camera and FPGA settings. A key feature of this software is its ability to take FIFO input from the FPGA target, which buffers and streams data from various sources, including VSYNC signals, behavioral stimulation signals, position and velocity data from the rotary encoder, behavioral properties, real-time averaged signals from LEDs matched to the frame duration, and time stamps with microsecond-precision. The VSYNC is a synchronization signal used in camera control that indicates the start of the image sensor’s exposure cycle.

The FPGA target software constitutes the actual code running on the FPGA. It has several modules that generate pulses to control the cameras and LEDs, edge detectors, counters and quadrature encoders. There is also a pulse selector for dual-GEVI studies and a real-time averaging module that provides a convenient readout of LED power, resampled to the frame-rate. The FPGA target software communicates with the host software via a FIFO, containing the data specified above.

#### Mouse preparation for mesoscope TEMPO measurements

Mice (aged 4–16 weeks at start) underwent two surgical procedures under isoflurane anesthesia (1.5%–2% in O_2_). The first procedure involved virus injection, to express the fluorescent proteins. In the second procedure, we implanted either a glass window (7–8 mm diameter) in the neocortex covering ∼20 brain regions (**Figures 4,5,7,S7**) or a 3 mm-diameter optrode device in the hippocampus (**Figure 6,S7**, see description above). For cortical imaging, we injected the PHP.eB virus retro-orbitally (RO) in ∼5 week-old mice. For TEMPO imaging experiments of one cell-type (**Figures 4,5,S7**), we injected 100 µL of a Ringer solution containing 1.5E11 GC of AAV2/PHP.eB-EF1α-DIO-ASAP3 virus and 1E11 GC of AAV2/PHP.B-CAG-mRuby2 virus. For imaging studies of two neuron-types (**Figure 7A–N**), we injected 100 µL of Ringer solution containing 1.5E11 GC of AAV2/PHP.eB-EF1α-DIO-ASAP3 virus, 2.5E11 GC of AAV2/PHP.eB-CaMKII-Varnam2 virus, and 1E10 GC of AAV2/PHP.eB-CAG-cyOFP virus. For studies of two neuron-types with reversed GEVI assignments (**Figure 7O,P**) we injected 100 µL of Ringer solution containing 3E11 GC of AAV2lPHP.eB-CaMKII-ASAP3 virus, 1.5E11 GC of AAV2/PHP.eB-EF1a-DIO-Varnam2 virus, and 1E10 GC of AAV2/PHP.eB-CAG-cyOFP virus. For controls in which neither fluor was a GEVI (**Figure S7A**), we injected 100 µL of a Ringer solution containing 1.5E11 of AAV2/PHP.eB-CAG-GFP virus and 2E12 GC of AAV2/PHP.B-CAG-mRuby2 virus. For hippocampal imaging studies (**Figure 6**), we locally injected viruses into dorsal CA1 at coordinates relative to Bregma of (AP, ML, DV): -1.8, -2.5, -1.3. We injected 500 nL of a viral solution containing 5E9 GC of AAV2/PHP.eB-EF1α-DIO-ASAP3 virus and 1E9 GC of AAV2/PHP.B-CAG-Ruby3 virus. See above for a description of the implant for hippocampal imaging.

For neocortical studies, we implanted a glass window (7–8 mm diameter) on the right hemisphere of ∼12-week-old mice, ipsilateral to the retro-orbital injection. For 7-mm-diameter windows we used #64-0723 coverslips from Warner Instruments. To create larger windows, we manually cut them from #22-266-940 cover glasses from Fisher Scientific. The surgical procedure was similar to that of our previously published work.^137^ We anesthetized mice with isoflurane (4% for induction and 1%–2% during surgery) on a stereotaxic frame (Model 963, David Kopf instruments). To mitigate post-operative pain, we injected carprofen (5–10 mg/kg) subcutaneously prior to any surgical incision. We applied eye ointment to the mouse’s eyes to maintain their moisture. A heating pad (FHC, 40-90-8D) maintained the mouse’s body temperature at 37°C.

We removed the skin atop the cranium and mechanically removed all soft tissues from the skull surface with a scalpel. Next, we used a 0.7-mm-diameter micro drill burr (#19007-07, Fine Science Tools) to perform a craniotomy along AP 3.0 to AP -5.0 (referenced to the mouse Bregma), approximately matching the diameter of the implanted glass window size. During this most critical step, it was important to not drill past the desired diameter so that the skull and window edges fit properly. Moreover, it was important to minimize damage to the dura to prevent excessive bleeding. We applied mammalian Ringer’s solution (#11763-10, Electron Microscopy Science) throughout the drilling to rinse off debris and minimize tissue heating. After completing the drilling, we disconnected the resulting circular-shaped bone piece from the surrounding skull. Once the bone piece was disconnected from the skull, we centered a plastic pipette tip atop the detached skull bone. We then replaced the bone piece with the glass coverslip and pressed the window ∼100 µm downward to flatten the cortical brain tissue beneath the window. We dried the edges of the window implant with surgical eye spears (#1556455, Butler Schein Animal Health) and glued the window to the skull with UV-light-cured adhesive (#4305, Loctite), while protecting the mouse’s eyes with aluminum foil to prevent UV-light damage. To allow head-fixation during brain imaging, we installed a stainless steel annular headplate (12 mm inner diameter), concentric with the glass window, and we filled the gap between the headplate and the skull with dental cement (#10000787, Fisher Scientific).

We transferred the mouse to a recovery cage and placed it on a heating pad until it awoke. We then returned the mouse to its home cage and provided food and water on the cage floor without use of a food hopper. We subcutaneously administered the mice carprofen (5 mg/kg) for the first 3 days after surgery to reduce post-surgical discomfort. For the first 7 days after surgery, we checked all mice daily for signs of distress. Mice recovered for at least 2 weeks before imaging experiments began.

#### Recording sessions for TEMPO microscopy

We performed all recordings during the mouse’s light cycle. Before every recording session, we first removed cage dust deposited onto the mouse cranial window using a cotton tip sprayed with ethanol and immediately blew away the excess ethanol using an air duster tank. Next, to align the mouse cranial window orthogonal to the microscope optical axis, we used two goniometers to control the mouse head-stage in two rotational angles (tilt and yaw) We used a ∼1 mW laser pointer (CPS532-C2, Thorlabs) that was directed vertically onto the window and aligned the reflection off the cranial window back to the laser output. For experiments using ketamine-xylazine anesthesia (**Figure 4, S6**), recordings began ∼20 min after anesthesia administration, when mice became immobile, and recording durations were between 2–10 min. Experiments involving head-restrained visual simulation (**Figures 5, 7, S6, S7**) were similar to the experiments performed with fiber optic TEMPO (**Figures 3, S2, S7**), and recordings lasted up to 10 min. Experiments involving head-restrained locomotion (**Figures 6, S7**) were similar to the experiments performed with fiber-optic TEMPO (**Figures S2, S3**), and recordings lasted up to 10 min. Recording depth for cortical imaging was ∼200-300 µm (**Figures 4, 5, 7, S6, S7**). The recording depth for hippocampal imaging studies was ∼50-100 µm (**Figures 6, S7**). Across all experiments, we used up to 150 mW of blue LED and 110 mW of green LED illumination across the 7–8 mm wide field-of-view.

#### Histology and fluorescence microscopy of brain slices

At the end of *in vivo* experimentation, we deeply anesthetized mice with isoflurane. We transcardially perfused the mice with phosphate buffered saline (PBS) (pH 7.4), followed by 4% paraformaldehyde (PFA) in PBS. We fixed brains in PFA at 4°C for at least 24 h and prepared tissue sections (∼100 µm) using a vibrating microtome (VT1000S, Leica). We washed the sections with PBS several times and incubated them in 150 mM glycine in PBS for 15 min to quench fluorescence induced by PFA. We then washed sections 3 times in PBS, and mounted sections with Fluoromount-G (Southern Biotech).

To capture the native fluorescence of the GEVIs (ASAP3, Ace-mNeon1, Varnam1 or Varnam2) and reference fluorescent proteins (GFP, mRuby2, mRuby3 or cyOFP), we used a commercial confocal system (Leica SP8) with a 0.75 NA, 20x immersion objective and hybrid photomultiplier tube. We used continuous wave laser excitation, of 488 nm or 561 nm wavelength, along with green fluorescence (525/50) and red fluorescence (600/50) filters (**Figures 1,3,7,S2,S6**).

#### Data processing for fiber-optic TEMPO recordings

For all fiber optic TEMPO recordings **(Figures 1–3, S1–S3,S7**), we analyzed the data using a custom-written uSMAART analysis package (*to be made publicly available at publication*) that runs in MATLAB (Mathworks). In brief, we first loaded the raw, unfiltered data into MATLAB. Next, we applied a notch bandstop filter to remove the artifacts induced by the dynamic diffuser from the decoherence stage (see above, **Instrumentation for fiber-optic TEMPO measurements**). We designed this bandstop filter using the matlab function, *designfilt()*, and created an infinite impulse response, 8th-order Butterworth filter with center cut-off frequencies of 300 ± 1 Hz or 600 ± 1 Hz to remove the fundamental and its first harmonic, respectively. We applied this filter to the voltage and reference traces using the zero-phase-delay matlab implementation, *filtfilt()*.

To remove the photobleaching dynamics, we then detrended the fluorescence signals by normalizing the trace by the time-varying baseline fluorescence, *F*_0_, as estimated using a temporal low-pass filter (4th-order Butterworth filter, 0.5 Hz cut-off frequency). As is common in fluorescence studies, photobleaching typically exhibited multi-exponential kinetics, which we characterized through parametric fits. For example, we found that the characteristic time interval, *t*_1/2_, over which ASAP3 fluorescence decays to half of its initial value is *t*_1/2_ = 2.5 h, using an illumination intensity of ∼2.8 mW/mm^2^. We calculated this value by fitting the time-course of ASAP3 photobleaching with a tri-exponential decay, with 3 time-constants (*t*_fast_ = 26 s; *t*_mid_ = 300 s; *t*_slow_ ≈ 5.5 h), which provided a substantially better fit than a bi-exponential function. For Varnam2 and mRuby2, we determined that *t*_1/2_ is 6.4 h for an illumination intensity of 0.25 mW/mm^2^. by using a bi-exponential fit (*t*_fast_ = 291 s; *t*_slow_ ≈ 9.7 h). These *t*_1/2_ values are much longer than those reported for voltage imaging studies at single action potential resolution (*e.g*., *t*_1/2_ <10 min for ASAP3^58^, Ace-mNeon2 and Varnam2^1^), since TEMPO uses ∼20–200 times weaker illumination intensity levels than imaging studies of neural spiking.

Finally, to remove the contributions of non-voltage dependent artifacts and noise sources to the fluorescence voltage trace, we unmixed the voltage and reference channel traces using our custom-designed convolutional unmixing algorithm (below).

In dual cell-type TEMPO studies (**Figures 3,S2**,**S3,S7**), prior to the convolutional unmixing step, we included a decrosstalking step to correct for fluorescence crosstalk between the demodulated cyOFP and Varnam2 time traces. The rationale for this step is that fluorescence signals collected from cyOFP will, by design, include some crosstalk from the voltage-dependent Varnam2 indicator. Because the blue laser excites both of these fluors, ∼10.3% of the fluorescence collected from Varnam2 appears as crosstalk in the demodulated cyOFP signal, as calculated under the condition that the blue and green laser emissions are equal in intensity at the specimen. The demodulated cyOFP signal also includes some crosstalk from ASAP3 emissions, because ∼8.8% of the collected ASAP3 photons bleed into the red fluorescence detection channel. However, we observed empirically that crosstalk from Varnam2 was the dominant contaminant of the cyOFP signal, because (i) Varnam2 fluorescence was generally of greater amplitude than ASAP3 fluorescence, and (ii) in real experimental situations, the blue laser was typically set at a ∼10-fold higher intensity level than that of the green laser (see values above) to compensate for the weaker expression of ASAP3.

To perform the decrosstalking, we regressed the demodulated cyOFP trace against the demodulated Varnam2 trace, using bandpass-filtered versions of both traces centered on the relevant frequency band (3rd-order Butterworth filter with cut-off frequencies set at 3-7 Hz for visual cortical studies and 5-9 Hz for hippocampal studies), which contains well-characterized voltage dynamics and is generally free of artifacts. We weighted the unfiltered Varnam2 trace by the value of the regression coefficient and subtracted the resultant from the unfiltered cyOFP trace, which was then used as one of the inputs to the convolutional filter. This fluorescence decrosstalking step ensures that the subsequent convolutional filtering algorithm does not unmix or alter the voltage signals within the Varnam2 trace.

#### Frequency-dependent unmixing of voltage signals

We developed a custom algorithm for unmixing hemodynamics and other artifacts captured in the reference fluorescence channel from the voltage signals captured in the GEVI fluorescence channel (**Figure S5**). This convolutional filtering algorithm outperforms a frequency-independent regression^138^ and presumes that both artifacts and voltage signals have characteristic frequency domain representations, and so it performs a frequency-dependent regression between the two channels. The frequency-dependent amplitudes and phases used for this regression constitute a convolutional filter. Although this appears to be the first instance of convolutional filtering applied to voltage imaging, the concept of signal estimation through filtering was introduced decades ago by Kolmogorov^139^ and Wiener^140^. Today, Wiener filtering is widely used in digital signal processing, control, image processing, and other fields of engineering.^83^

Here, we describe the algorithm for the case in which there is one GEVI. In studies with two GEVIs, we unmixed the reference channel content from each voltage channel individually. We treat the case in which one detection channel collects fluorescence from a GEVI (*e.g.*, ASAP3 or Varnam2) and the other collects emissions from a reference fluor (*e.g.*, mRuby2 or cyOFP) (**Figure S5A**). The data generally have 5 notable characteristics: (1) the GEVI channel contains a mixture of neural voltage-dependent and -independent signals, whereas the reference channel only contains signals of the latter type, *i.e.*, that are independent of neural membrane potentials and thus are artifacts for the purposes of voltage imaging; (2) the biological artifacts (*e.g.*, hemodynamics and brain motion) and instrumentation noise (*e.g.*, illumination fluctuations) generally have intrinsic temporal frequency signatures; (3) oscillatory hemodynamic signals are generally the artifacts of the greatest magnitude and have characteristic frequencies concentrated around that of the heartbeat and its harmonics (**Figure S5B**); (4) hemodynamic oscillations are present in the same frequency bands and are generally coherent between the two channels, but do not always oscillate with the same phase in the two channels (**Figure S5B**); (5) the spatial distribution of hemodynamic artifacts is non-uniform across the field-of-view (see *e.g.*, **Figure S5I**).

We implemented our unmixing algorithm using the assumption that the signals in the GEVI channel, *G*(*t*), are a sum of the voltage signals, *V*(*t*), plus a voltage-independent component, *H*(*t*):

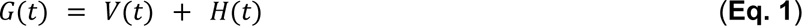

Based on above observations (1–5), we formulated the problem of estimating the non-voltage signals, *H*(*t*), as a Wiener filter estimation problem in which we seek a filter, *F*(*t*) such that the convolution of the reference channel trace, *R*(*t*), with the filter, *F*(*t*), yields *H*(*t*):

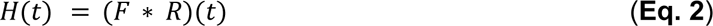

Our physiological intuition for why this formulation makes sense is that *H*(*t*) predominantly arises from changes in the optical properties of the tissue, due to time-variations in oxygenated (HbO) and deoxygenated (Hb) hemoglobin content. These two forms of hemoglobin will have distinct influences on the GEVI and reference fluorescence signals, but these influences are not independent. A simplified way of depicting the dynamics relating the concentrations of Hb and HbO is to represent them as a linear dynamical system, which implies a linear dynamical relationship between *R*(*t*) and *H*(*t*) that can be represented as a convolution in the time-domain. With this framework, the main steps of our algorithm are (**Figure S5C,D**):

1. We split the traces from the GEVI, *G*_*k*_(*t*), and reference channels, *R*_*k*_(*t*), into *N* temporal segments of duration *τ* (typically 0.5–2 s) indexed by the variable *k* = 0, 1, … *N*, and with fractional overlap 1 − γ ∈ [0, 1] (typically 0.75)

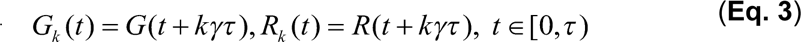 We compute the corresponding Fourier transforms *g*_*k*_(ω) and *r*_*k*_(ω):

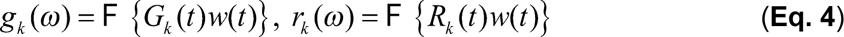

where F{·} denotes a Fourier transform and *w*(*t*) is a windowing function (*e.g.*, a Hann window).
2. We estimate the Wiener filter in the frequency domain, *f*(ω), by concurrently fitting a complex coefficient *f*_*k*_(ω) for every frequency ω to approximate the GEVI channel signal *g*_*k*_(ω) with the reference channel signal *r*_*k*_(ω) for all *N* segments:

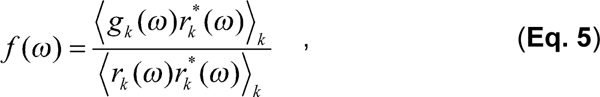 where *r*\* is the complex conjugate of *r* and ⟨…⟩_*k*_ denotes averaging over the *N* segments. We limit the upper bound of the filter amplitude relative to the amplitude of the linear regression estimated at the heartbeat frequency ω_0_, with α > 0 (typically 1.1, see **Figure S5M**)

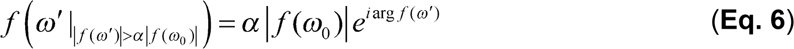

If ω_0_ is the fundamental harmonic of the heartbeat frequency, the amplitude _|_*f*(ω)_|_ can be interpreted as a linear regression coefficient between the channels filtered at the heartbeat frequency. We set the spectral amplitude limit for the filter to α_|_*f*(ω)_|_, thus explicitly prohibiting unbounded amplification.
3. We apply an inverse Fourier transform to find the time-domain representation of the filter *F*(*t*):

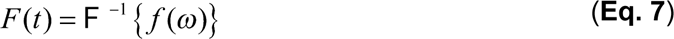 We convolve *F*(*t*) and *R*(*t*) in the time-domain to estimate the non-neuronal contribution *H*(*t*), (**Eq. 2**), and subtract *H*(*t*) from the GEVI signal *G*(*t*) to recover the unmixed voltage signal *V*(*t*):

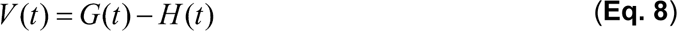

The above algorithm can be interpreted as a rank-constrained linear regression in the frequency domain. We fit a complex-valued scaling coefficient *f_k_*(*ω*) for every frequency *ω* to simultaneously approximate the voltage channel signal *g_k_*(*ω*) with the reference channel signal *r_k_*(*ω*) for all segments, *k*. The rank of the regression (*i.e.*, the number of free parameters or the number of points in the filter) is equivalent to the filter length, *τ*, and thus inversely proportional to the spectral resolution.

We determined the appropriate segment length *τ* (usually 1.0–1.5 s) and optimal relative spectral limit, *α* (1.0–1.3) by splitting the data in half into train and test segments and minimizing the remaining signal in the test movie segment after unmixing with respect to the parameters *τ* and *α*. The unmixing procedure and the results of the paper are robust with regard to minor changes in the values of these parameters (**Figure S5M**). Larger segment lengths allow for greater spectral resolution of the filter, whereas smaller segment lengths result in more robust estimation in low SNR recordings.

In some recordings, the noise floor in the reference channel was higher than that in the GEVI channel (see *e.g.*, **Figure S5H**). In such recordings, unmixing would lead to masking of high-frequency signals underneath a noise floor inherited from the reference channel. Therefore, we performed a pre-filtering step that consisted of estimating *H*^*avg*^(*t*) using a spatially averaged reference movie, *R*^*avg*^(*t*), and then removing *H*^*avg*^(*t*) at the single pixel level (**Eq. 9**):

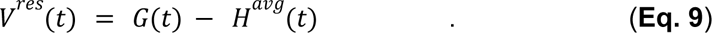

Then, we performed a second convolutional filtering at the single pixel level between signals in the reference channel, *R*(*t*), and those in the voltage channel, *V^res^*(*t*). The pre-filtering step is reasonable because, as non-neural activity generally does not travel spatially, it boosts the *SNR* of *R^avg^*(*t*). The suitable scale of spatial averaging depends on the relative noise levels between the two channels and was usually set to be ∼0.5–1.5 mm. Although more spatial averaging leads to improved SNR values (*e.g.*, compare the single-pixel and spatially-averaged traces in **Figure S5H,I**), reduced spatial averaging allows the spatial variations of the hemodynamics to be captured at higher accuracy. Overall, compared to a simple (*i.e.*, frequency independent) regression^138^, our unmixing method better removed broad- and narrow-band artifacts and did not transfer noise from the reference to the GEVI channel, which was critical for detecting high-frequency activity with high sensitivity (**Figure S5E–J**).

#### Processing of TEMPO mesoscope video data

We preprocessed the data using a custom pipeline written in MATLAB, (*Voltage Imaging Analysis*, *to be made public upon publication*). This pipeline comprised the following main steps:

i. *<u>Loading and conversion:</u>* We first loaded the raw, 16-bit *.dcimg* video files from the GEVI and reference channels into the computer RAM in small batches (∼10% of the available RAM memory). We spatially downsampled each movie by applying 8 × 8 binning using the Matlab function *imresize3()*, with the ‘box’ interpolation method that performs averaging over all 64 pixels. We saved the 3D matrices as single floating point numbers (32-bit) in HDF5 file format to a separate RAID0 SSD array. The HDF5 files also contained the recording settings (*e.g.*, imaging frame-rate).
ii. *<u>Motion correction:</u>*Both GEVI and reference channel movies underwent rigid motion-correction using phase correlation image registration^141^ within the NoRMCorre^142^ software package and were then cropped to remove parts of the images lacking fluorescently labeled brain tissue. We used the rigid registration mode of NoRMcorre, instead of non-rigid registration, as our image data primarily exhibited rigid displacements, and we found the former mode of operation to be more robust and faster than the latter. The NoRMcorre parameter shift_method was set to ‘cubic’, instead of the default ‘fft’.
iii. *<u>Registration</u>*: We next registered the GEVI and reference motion-corrected movies to correct for any static spatial displacements between them. We did this by estimating a rigid transformation, which corrects for translation and rotation, between representative image frames of spatially bandpass-filtered versions of the green and red fluorescence movies. The image from the green channel served as the reference because of its superior optical contrast showing the vasculature pattern. We estimated the affine transformation between the green and red images using 5 different approaches: (1) an intensity-based method (MATLAB function *imregcorr()*), (2) a phase correlation-based method (MATLAB function *imregcorr()*) and (3) a mutual information correlation-based method (MATLAB function *imregtform()*). The last two approaches were implemented twice, using either the ‘mono-modal’ or ‘multimodal’ option, yielding a total of 5 methods. The ‘mono-modal’ option assumes similar brightness and contrast between the two images, whereas the ‘multi-modal’ option does not.^143,144^ All 5 approaches performed differently for different movies. Therefore, we evaluated their respective performance using the structural similarity index measure (SSIM; MATLAB function *ssim()*), computed between the fixed green image and transformed red image. We used the image registration approach that yielded the highest SSIM value and then applied it to all individual movie frames of the red channel.
iv. *<u>Decrosstalking</u>*: To remove any residual crosstalk between the GEVI and reference channels, we first estimated the fraction of fluorescence bleedthrough from the optical filters’ properties. In practice, ∼7% of ASAP3 (∼9.5% for Ace-mNeon1) fluorescence signals bled into the reference channel. We scaled the green fluorescence movie (usually the GEVI channel) by the estimated coefficient and subtracted the resultant from the red fluorescence movie (usually the reference channel). We did not correct from bleedthrough of the red fluor into the green fluorescence detection channel, since the fluorescence filters were expressly chosen to keep this value negligible (∼0.1% for bleedthrough of mRuby or Varnam into the green channel).
v. *<u>Detrending.</u>* To account for fluorescence photobleaching, at each time point in the fluorescence video, *F*(*t*), we normalized each pixel by *F*_0_(*t*), the time-varying value of the pixel’s baseline fluorescence. The resultant is the detrended movie, *F*(*t*)/*F*_0_(*t*). For **Figures 4–6, S6**, the detrending involved normalizing the trace by the time-varying baseline fluorescence, *F*_0_, which we estimated using a first-order exponential fit (see typical values, in **Data processing for fiber-optic TEMPO recordings**, above). We then applied a temporal high-pass filter to the data [type-1 FIR filter (symmetric, even order, linear phase; >10^5^ stopband attenuation, <10^-2^ passband ripple) designed using the *design()* function of the MATLAB DSP System Toolbox; 0.5 Hz cutoff frequency for anesthetized mice, 1.5 Hz for awake mice]. For **Figure 7**, the detrending involved normalizing by the time-varying baseline fluorescence *F*_0_, as estimated using a temporal low-pass filter (4th-order Butterworth filter, 0.5 Hz cut-off frequency).
vi. *<u>Unmixing.</u>* To remove broad- and narrow-band hemodynamic artifacts as well as residual brain motion, we used the custom convolutional unmixing approach described above.

For dual cell-type recordings (**Figure 7**), we incorporated an additional step in which we upsampled the videos in the temporal domain by twofold and temporally shifted them by half of the nominal frame duration, *i.e.*, by 1 frame after interpolation. This allowed us to eliminate any phase shifts between signals obtained from cameras using interleaved light sources, thereby simplifying the unmixing of the fluorescence signals.

#### Fast denoising of TEMPO movies

For the analyses of **Figure 6**, after applying the frequency-dependent convolutional unmixing algorithm (above), we improved the signal-to-noise ratios of the videos by applying a custom denoising algorithm based on a singular value decomposition (SVD) of the movie data. Because the runtime of SVD scales cubically with the size of the movie, we first broke the unmixed movie into temporal segments, each of 2500 frames in duration. We denoised each segment separately and then re-concatenated them to obtain a denoised version of the full movie.

To denoise each movie segment, we reshaped it into a two-dimensional matrix, **M** ∈ R*^p^* ^✕^ *^d^*, where *p* is the number of video frames in the segment (*i.e.*, 2500) and *d* is the total number of pixels in the field of view. Using SVD, we decomposed **M** as a product, **M** = **U**・**C**, where **U** ∈ ℝ*^p^* ^✕^ *^k^* is a set of *k* low-rank components, and **C** ∈ ℝ*^k^* ^✕^ *^d^* are weighting coefficients. The components **U** are assumed to be semi-unitary, without loss of generality, and were obtained by computing the SVD of **M**. The number, *k*, of low-rank components retained in **U** was five; we found that this choice removed the grainy appearance of noise in the videos of wave dynamics without altering the underlying depiction of wave propagation. We calculated the coefficients as **C = U***^T^*・**M**. For each row of the coefficient matrix, after reshaping it back into a two-dimensional image, we denoised it with the MATLAB routine *denoiseImage()*. This routine calls the MATLAB function *denoisingNetwork()*, which uses a deep convolutional neural network for denoising.^145^ The resultant is a set of denoised coefficients, **Ĉ**, from which we computed the denoised movie segment, **U**・**Ĉ**. We then reshaped the movie segment back to its original dimensions. After denoising all the movie segments, we assembled the denoised version of the full movie and proceeded to the analyses of hippocampal waves.

#### Computations of coherence between optical and electrical traces

To assess the coherence between the optical and electrical traces (**Figures 1–3, S1–S3, S7**), we applied a Fourier-based approach and used the MATLAB function *mscohere() to* compute the coherence magnitude between the two time series, and we used the MATLAB function *cpsd()* to compute the coherence phase between the two time series. We applied both functions to 1-s-long data segments, chosen such that temporally successive segments from the traces overlapped by 0.8 s. In imaging studies, we computed the coherence between the LFP and the TEMPO voltage signal at each pixel in the field-of-view (**Figure 6E,F**). The Fourier approach for assessing coherence levels has been commonly used in neurophysiology.^146–150^ However, we also checked on a subset of our data that this method yielded qualitatively similar results as a wavelet-based approach to assessing coherence. This alternative method relied on the MATLAB function *wcoherence()*, which provides a time-frequency representation of the coherence between a pair of non-stationary time series without using computational time-windows of fixed duration.

In **Figure 1H,I**, to identify the resting and running epochs of freely behaving mice for which we did not have behavioral video tracking, we used the LFP time series to isolate periods of high versus low theta power, which is a well-accepted proxy.^8,63,151,152^ We validated this data segmentation approach on mice in which we had both the LFP and behavioral video tracking.

#### Behavioral video tracking and pose estimation for freely moving mice

To track the mouse’s two-dimensional (*x-y*) position in the raw .avi behavioral movies (**Figures 1, 3**), we used the deep learning-based animal tracking algorithm, DeepLabCut (version 2.2.1).^153^ We labeled the mouse’s head and the base of its tail in 187 image frames that were randomly chosen from 5 video recordings. We split this dataset into portions used for training and testing in a 95/5 ratio. We used a ResNet-50-based neural network^154,155^ and 500,000 training iterations with default parameters. We used a p-cutoff of 0.6 to condition the *x-y* coordinates, which led to a training error of 1.24 pixels and a test error of 3.46 pixels (image size was ∼300 × 300 pixels). We then used this network to analyze behavioral videos taken under similar conditions. Because the estimated position of the base of the mouse’s tail was more accurate than the estimated head position, we used the displacements of the tail base across image frames as the basis for computing the mouse’s velocity.

#### Characterizations of cross-frequency coupling

To compute cross-frequency coupling between two oscillations of different frequencies (**Figures 2, S3**), we first bandpass-filtered the raw signal at the low-frequency bandwidth of interest. From this filtered version of the trace, we estimated the analytic signal using the Hilbert transform (MATLAB function, *hilbert()*) and computed the phase using the function *angle()*. Next, we estimated the wavelet spectrogram of the raw signal using the MATLAB function *cwt()* and computed the average across the high-frequency bandwidth of interest. Finally, we identified every phase reset of the low-frequency oscillation and aligned the high-frequency signal accordingly.

In studies of cross-frequency coupling (**Figure 2C,D,G–J** and **S3J–L**), the convention used for the phase of the carrier oscillations (delta or theta) was that zero degrees refers to the trough of the carrier oscillation, *i.e*., the greatest hyperpolarization in the TEMPO signal (**Figure 2C,D** and **S3J–L**) or the greatest hyperpolarization in the extracellular electric field recording (**Figure 2G–J**).

To detect periods of epileptic spikes (**Figure 2M**), we high-pass-filtered the raw LFP signal (3 dB cutoff of 25 Hz; 6th-order Butterworth) and detected spikes using the MATLAB function *findpeaks()*, with the input arguments *minpeakwidth*, *maxpeakwidth*, and *minpeakprominence* set at 2 ms, 20 ms, and 1000 µV, respectively.

#### Continuous and event-related spectrograms

To compute continuous spectrograms (**Figure 3**), we applied the MATLAB function *spectrogram()* to the LFP trace. To compute event-related spectrograms (**Figures 3,5-7,S1,S2,S6**), we first computed the wavelet spectrogram of the entire time-series using the MATLAB function *cwt()*. Then, we isolated the time points of each stimulus presentation and determined the mean spectrogram, averaged across the set of all stimulus presentations.

#### Identification of hippocampal layers using electrophysiological landmarks

To identify the boundaries between the different layers of CA1 hippocampus in our 32-site silicon probe LFP recordings, we examined the voltage traces recorded from each site during intervals in which there were either sharp-wave ripple (SWR) events or theta oscillations (**Figure S3A,B**). For each of the 32 sites, we arranged plots of the LFP traces in order of the depth into CA1 tissue, temporally bandpass-filtered (3rd-order Butterworth) across either the ripple (120–180 Hz) or theta (5–9 Hz) frequency range. We identified the 4 canonical hippocampal layers using two electrophysiological landmarks: (1) the center of *stratum pyramidale* corresponds to the depth at which ripple oscillations have the greatest amplitude; (2) the anatomical boundary between *stratum radiatum* and *stratum lacunosum moleculare* corresponds to the depth at which theta oscillations change in both amplitude and phase.

#### Detection of hippocampal ripples from silicon probe recordings

To study ripple events (**Figures 3Q, S3F**), we applied the following detection algorithm using the data acquired on the recording site of our 32-site probe with the greatest ripple amplitude (**Figure S3A**). Specifically, to detect ripple events, we used a previously described method that involves bandpass filtering the raw LFP signals at the ripple band frequency (120–200 Hz; 3rd-order Butterworth filter) and then calculating the normalized squared signal (NSS).^156–158^ To compute the NSS, we first squared the filtered signal, then applied a moving maximum (MATLAB function *movmax()*) with a 20-ms-window, and then performed a smoothing using a 20-ms-window (MATLAB function *smooth()*). We scaled the resulting trace between 0 and 1 using the *rescale()* MATLAB function. To isolate ripple events, we then identified local maxima using the MATLAB function *findpeaks()* with the following option parameters, ‘MinPeakProminence’, ‘MinPeakDistance’, ‘MinPeakWidth’ and ‘MaxPeakWidth’ set to, respectively, 10%, 10 ms, 10 ms and 200 ms. We chose these values based on characteristics of hippocampal ripples in the published literature.^156–158^ All extracted ripple events were isolated within a 300-ms-window. Finally, we visually curated all automatically identified ripple events so as to exclude non-stereotypical ripples (*e.g.*, events corrupted by high-frequency noise or events that were recorded on all 32 recording channels, instead of being localized to the hippocampal pyramidal layer).

#### Seed-pixel spatial correlation maps

To characterize wave propagation, we selected a seed pixel located in the brain region of interest and computed the cross-correlation between the voltage signal at the seed pixel and every other pixel. We used the Matlab function function *xcorr()* to compute the cross-correlation map, *X*_*i*,*j*_(*a*, *b*), where *i* and *j* denote spatial indices and the pair, (*a, b*), denotes the temporal indices of a pair of video frames. In this way, we obtained cross-correlation and delay maps characterizing the spatiotemporal propagation of the wave present in the seed pixel.

#### Calculations of velocity distributions in visual cortex

To characterize traveling waves in **Figures 4** and **5**, we used the cross-correlation function to determine the level of similarity between pairs of time traces and the delay between them in a spatially resolved manner. For each pair of points in space, corresponding to pixels *i* and *j*, we computed the time-dependent correlation coefficient, 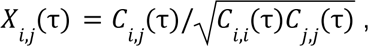 where *C*_*i*,*j*_(τ) = ∫ *V*_*i*_ (*t*)*V*_*j*_ (*t* − τ)*dt*. We then estimated the speed, *v*_*i*,*j*_, at which a wave traveled between two points, (*x_i_*, *y_i_*), (*x_j_*, *y_j_*), using τ*_max_* = *argmax*_τ_ {*X*_*i*,*j*_} and 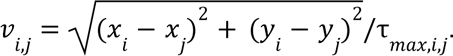 We applied a threshold *X*_*i*,*j*_> *X*_*min*_ to restrict speed calculations to pairs of points at which the voltage waveforms were similar. We chose *X*_*min*_ values between 0.6–0.9, balancing the spatial range of the wave and the signal-to-noise ratio of the spatially resolved waveforms. To estimate wavefront gradients, we parametrically fit the spatial map of wavefront delays, τ(*x*, *y*), using least-squares fitting to the equation τ(*x*, *y*) = *Ax* + *By* + *C*. We then used the fit coefficients to calculate the local wavefront velocities, 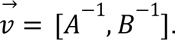 To improve the quality of the fit, we often performed additional spatial binning. We obtained the flow maps of **Figures 4** and **5** by normalizing all velocity vectors to a uniform length. We computed the distributions of wave speed and traveling direction based on the velocity vector distributions.

#### Calculations of velocity distributions in hippocampus

To characterize the speeds and directions of traveling voltage waves in **Figure 6**, we performed these four steps: (1) Wave detection; (2) Space-Time projection; (3) Wave crest identification; (4) Linear Regression. We began these analyses using movie data that had already undergone convolutional unmixing (see **Figure S5** and above section, “*Frequency-dependent unmixing of voltage signal*s”) and denoising (see “*Fast denoising of TEMPO movies”*).

To detect wave occurrences, we (1) created a trace representing the time-dependent, net fluorescence averaged across the entire field-of-view; we identified wave events as intervals in which the net fluorescence exceeded 3 s.d. of the mean fluorescence signal (time-averaged across the entire trace). Next, we (2) projected the movie onto two orthogonal spatial vectors, **x** and **y**, onto its row and column dimensions, yielding a pair of one-dimensional movies (see *e.g.*, **Figure 6G**). Then, for each wave event, we (3) determined the time bin at which each pixel in the pair of one-dimensional movies attained its maximum fluorescence signal. Finally, using robust linear regression with the MATLAB function *robustfit()*, we regressed the spatial location of each pixel in the one-dimensional movies against the time at which the pixel achieved its signal maximum (*i.e.*, with the spatial location as the independent variable). These regressions yielded a pair of wavenumber values, α_*x*_ and α_*y*_, with physical units of s/mm. We determined the wave’s vector speed and direction by converting the vector wavenumber from a Cartesian to a polar coordinate representation. (To avoid infinite values for estimated speeds, we ensured that both Cartesian wavenumber values were significantly different from zero (p-value <0.01, as determined using the above robust regression).

To validate this method of speed determination, we applied it to simulated movies of waves with known characteristics; we assessed the accuracy of velocity determination across a range of speeds, propagation directions, and signal-to-noise ratios (SNRs). These tests provided confidence in the wave velocities estimated for empirical datasets (**Figure 6**). Overall, in comparison to the method of velocity determination described in the preceding Methods section, the 4-step method described here was faster and more robust when applied to low SNR movies, but was also unsuitable for waves that did not travel in a unidirectional manner.

#### Model to estimate the physiological time delay between two cell-types

During our joint studies of two neuron-types at once, we found that, owing to the voltage-dependence of voltage-indicator activation and deactivation time-constants, the different voltage-indicators used here (Ace2, pAce, and Varnam2) report oscillatory dynamics with slightly different phase delays in different neuron-types. In other words, the oscillatory phase-shift induced by temporal low-pass filtering varies slightly between the different indicators and neuron-types. Hence, to estimate the actual physiological time delay between the visually evoked 3–7 Hz oscillations of cortical PV interneurons and pyramidal cells in **Figure 7N,P**, we conducted a series of 7 ancillary experiments to characterize the phase shifts arising from indicator kinetics (**Figure S7D–F**).

Each ancillary experiment involved a different pair of voltage indicator assignments to cortical PV and pyramidal cells and yielded an apparent time delay between the visually evoked 3-7 Hz oscillations in the two neuron-types. We computed these time delays using the MATLAB cross-correlation function *xcorr()*, by applying it to every bout of oscillations measured in the green and red fluorescence channels (bandpass filtered between 3–7 Hz). The time at which the value of the cross-correlation function was maximized corresponds to the apparent time delay for a specific bout of 3-7 Hz activity, as studied with a specific pair of voltage indicators. For each mouse used to study a given indicator pair, we evoked 100 trials of oscillatory activity.

We then performed a *L*_1_-regularized linear regression to determine the values of the time delays induced by indicator kinetics for the 6 different assignments of our 3 indicators to the 2 neuron-types, as well as the actual time delay between the oscillations of PV and pyramidal cells. To do this, we sampled the apparent time delays for 100 different oscillatory bouts, for each of the 7 experiments. Using these 700 measurements, we computed mean and s.d. values for each of the 7 different time delays. The loss function for the regression was the sum, over all 7 experiments, of the squared difference between the mean measured time delay and the predicted value based on the estimate for the actual delay plus the difference of the two estimated indicator-induced delays for each experiment, with each term in the sum weighted inversely by the square of the experiment’s computed s.d. value. The regularization term was the Euclidean norm of the 6-dimensional vector comprised of all 6 indicator-induced delays. Fitting results were not sensitive to the value of the *L*_1_-regularization parameter, which we set to 0.01. Using this approach, and the constraint that indicator-induced delays must have positive values, we fit the regression model for 100 different samplings of measured time delays; from the results, we computed mean and s.d. values for each of the 7 different delay times across the 100 different samplings, as shown in **Figure S7E**.

To generate **Figure 7N,P** showing the distributions of estimated physiological time delays between PV interneurons and pyramidal excitatory neurons, we applied the linear regression model described above to the data taken in individual mice plus that from the 6 experiments in **Figure S7D-E** with distinct GEVI labeling assignments. In other words, for each imaging session studied in **Figure 7N,P** we determined the histogram of estimated physiological time delays through parametric fits to the measured values of the time-delays observed in each individual mouse, plus its 6 complementary experiments. We repeated the fitting procedure for each of the 4 recordings in **Figure 7N,P**.

#### Statistics

We performed statistical tests using standard MATLAB functions (MathWorks). For two-sample comparisons of a single variable, we used a non-parametric test, either the Wilcoxon rank sum or the Wilcoxon signed-rank test (Matlab functions, *ranksum()* and *signrank()*). All tests were two-tailed, unless stated otherwise.

---

**Figure S1:**
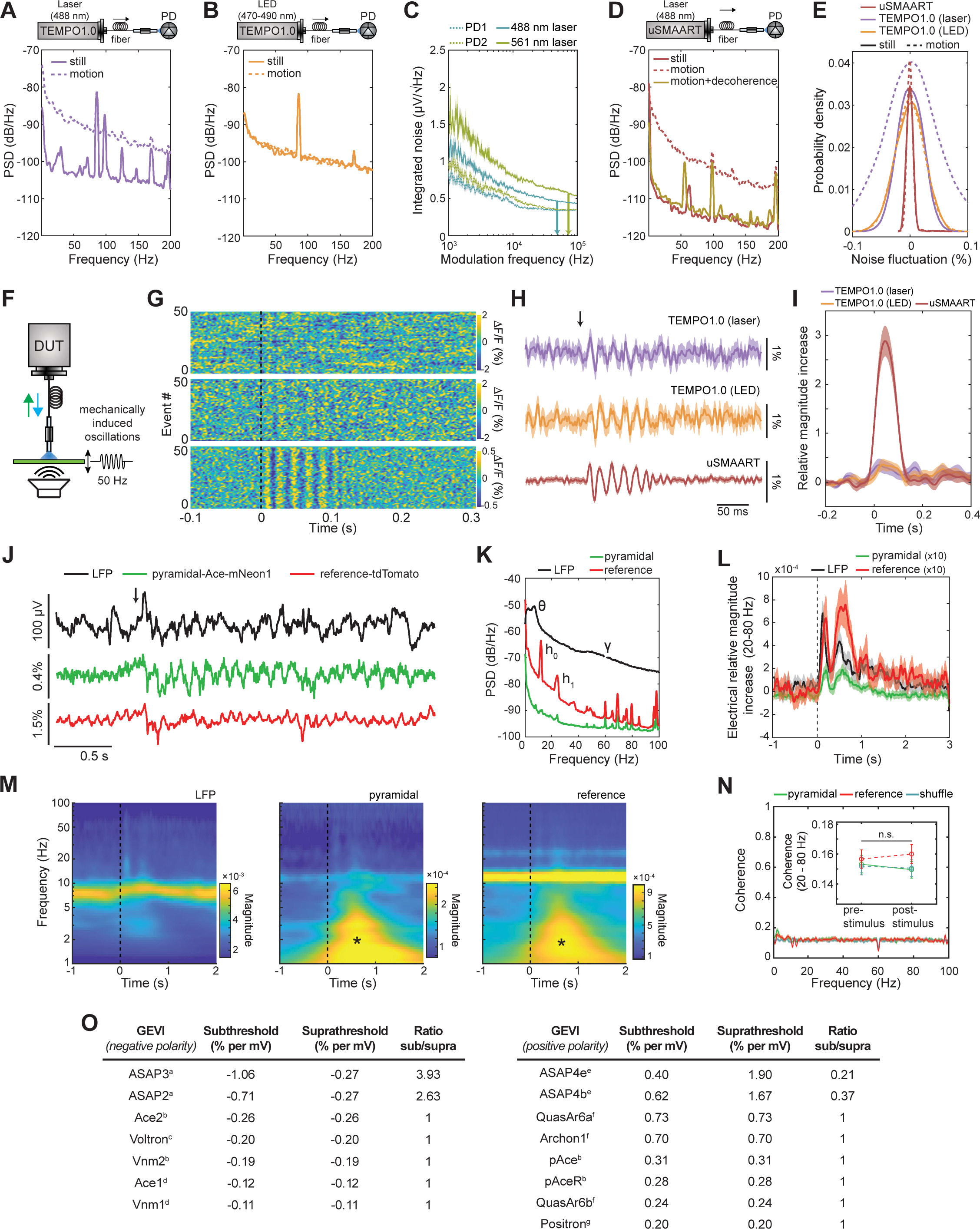
Benchmarking of uSMAART performance and tabulations of GEVI kinetics. (**A–E**) Coherent illumination from a solid-state laser had lower intensity noise fluctuations than the incoherent illumination of a light-emitting diode (LED) but led to motion-induced optical mode-hopping within the multi-mode optical fiber used for photometry. Passing the laser light through a decoherence module before coupling it into the optical fiber made the light intensity at the fiber output insensitive to movement of the fiber. (**A**) Power spectrum of illumination intensity noise, as measured for a 488-nm-laser (Obis; Coherent) that is commonly used for photometry, at the output of the multi-mode optical fiber, with (dashed curve) and without (solid curve) continuous shaking of the fiber. To perform a lock-in measurement, we sinusoidally modulated the laser output intensity at 3.5 kHz and used a digital lock-in amplifier to demodulate the signals from the photodetector to produce the plots shown (3.5 kHz demodulation frequency; 1 ms time constant). Mean illumination intensities at the output of the optical fiber were uniformly set to 50 μW in panels (**A, B**, **D, E**). (**B**) Power spectrum of illumination intensity noise, as measured for a commonly used LED (Thorlabs; ∼470–490 nm spectral bandwidth), at the output of the multi-mode optical fiber, with (dashed curve) or without (solid curve) continuous shaking of the fiber. Although shaking the fiber did not induce additional intensity fluctuations, the baseline intensity noise was substantially higher than that of the 488-nm-laser (compare to panel (**A**)). Illumination intensity modulation and signal demodulation were performed as in (**A**). (**C**) We compared the levels of noise power in our photodetector signals when the laser sources were either turned off (dashed curves) or on (solid curves), as a function of the modulation frequency used for the lock-in measurements. Notably, higher modulation frequencies led to reduced noise fluctuations. Thus, for all subsequent fiber-optic measurements, we used modulation frequencies of either 50 or 75 kHz (marked with arrows on the graph). (**D**) Power spectrum of illumination intensity noise, as measured at the output of the optical fiber for the 488-nm-laser source and using a modulation frequency of 50 kHz, without fiber movement (red solid curve), with continuous shaking of the fiber (red dashed curve), or with continuous shaking of the optical fiber plus the use of an optical decoherence module to prevent motion-induced mode-hopping (orange solid curve). Use of the decoherence module made the noise spectrum insensitive to fiber movement. Moreover, use of the 50 kHz modulation frequency reduced noise power levels by about 10 dB across the entire measurement bandwidth (0–200 Hz), in comparison to when the modulation frequency was 3.5 kHz (**A**). (**E**) Probability distributions of illumination intensity noise amplitudes for the optical configurations used in (**A**, **B**, **D**). (**F–I**) Sensitivity benchmarking with a fluorescent slide confirmed the enhanced ability to detect gamma-band (50 Hz) signals using uSMAART. (**F**) Schematic of an apparatus to generate artificial gamma band oscillations. We affixed a fluorescent slide to the membrane of a voice coil actuator and drove the voice coil with a 50 Hz sinusoidal voltage wave (100 ms in duration). We positioned the tip of the multi-mode optical fiber from the photometry system (either TEMPO 1.0 or uSMAART) nearby the slide such that it detected ∼150 pW of fluorescence photons, which is typical of *in vivo* TEMPO recordings from sparse neuron-types using 50 µW of illumination power. (**G**) Color raster plots showing relative changes in fluorescence evoked by the 50-Hz-wave and captured by either a TEMPO 1.0 system using a laser source (top plot), a TEMPO 1.0 system using an LED source (middle plot), or the uSMAART system of **Figure 1A** (bottom plot). Data from 50 different trials are shown in each plot, unfiltered. Dashed vertical line marks the onset of the 100-ms-duration sinusoidal (50 Hz) wave. (**H**) Traces of the mean time-dependent fluorescence during the 50-Hz-waves, averaged over 50 wave presentations for each of the 3 different fiber photometry systems. Shading: s.e.m. (**I**) Mean time-dependent signal amplitudes in the gamma band (45–55 Hz), determined via a wavelet transform of the traces of panel (**H**). uSMAART reported the artificial gamma bursts with ∼10-fold greater signal-to-noise ratios (SNR) than prior TEMPO instruments [gamma SNR increases for uSMAART *vs.* TEMPO 1.0 (laser): 2.9 ± 0.3 *vs.* 0.3 ± 0.1, p<10^-11^; uSMAART *vs.* TEMPO 1.0 (LED): 2.9 ± 0.3 *vs.* 0.3 ± 0.1, p<10^-11^ rank sum test]. Shading: s.e.m. (**J–N**) Re-analysis of stimulus-evoked fiber-optic TEMPO dataset from Marshall *et al*. 2016.^102^ (**J**) Example set of concurrently acquired LFP, voltage and reference fluorescence traces taken in a head-fixed mouse with our prior fiber-optic TEMPO system.^102^ The voltage indicator, Ace-mNeon1, was virally expressed in somatosensory cortical pyramidal cells via the *CaMKIIa* promoter, and the reference fluor was red fluorescent tdTomato. Black arrow marks the approximate start of vigorous whisker stimulation. The sustained post-stimulus oscillation at ∼4 Hz could be due to motion-induced mode-hopping noise in the optical fiber, although similar features are also visible in the LFP trace. (**K**) Power spectral density of all three signals in (**J**). The reference channel contains the heartbeat rhythm (h_0_) and its harmonics (*e.g.* (h_1_), as well as high-frequency instrument noise in the 40–100 Hz range, some of which is also apparent in the GEVI channel. (**L**) Wavelet spectrograms for all three signals in (**J, K**), averaged over stimulus presentations (*t*=0 is the time of stimulus onset). A theta-band oscillation is evident in the LFP recording (5-10 Hz), and the heartbeat (10–15 Hz) appears in the reference channel. Asterisks mark the onset of instrument noise at ∼1-5 Hz after stimulus onset in both the pyramidal cell and reference channels. (**M**) Average relative signal magnitudes in the gamma range for all 3 signals shown in (**J**). All three traces have similar time courses, but as shown in (**N**) the GEVI signal increase is not phase coherent with the LFP increase. (**N**) Plot of the mean coherence between the LFP trace and either the reference, pyramidal cell or temporally shuffled pyramidal signals, showing the lack of coherence between LFP and pyramidal voltage signals across the full 0.5-100 Hz frequency range. *Inset*: Gamma coherence between LFP and pyramidal cell voltage during pre- and post- stimulus presentation periods (LFP/pyramidal: p=0.78, LFP/shuffled pyramidal: p=0.68; LFP/reference: p=0.93; mean ± s.e.m., n=246 trials from n=3 mice). (**O**) Rates of fluorescence change per millivolt of membrane voltage change for negative (*left*) and positive (*right*) polarity GEVIs, as estimated for the subthreshold (–100 mV to –40 mV) and suprathreshold (–40 mV to +30 mV) voltage ranges. Superscripts denote values that were extracted from the following publications: (a) Villette *et al.*, *Cell*, 2019^58^; (b) Kannan *et al.*, *Science*, 2022^1^; (c) Abdelfattah *et al*., *Science* 2019^3^; (d) Kannan *et al*., *Nature Methods*, 2018^8^; (e) Evans *et al*., BioRxiv 2021^159^. (f) Tian *et al*., *Nature Methods*, 2023^160^; (g) Abdelfattah *et al.*, *Nature Communications*, 2020^161^.

**Figure S2:**
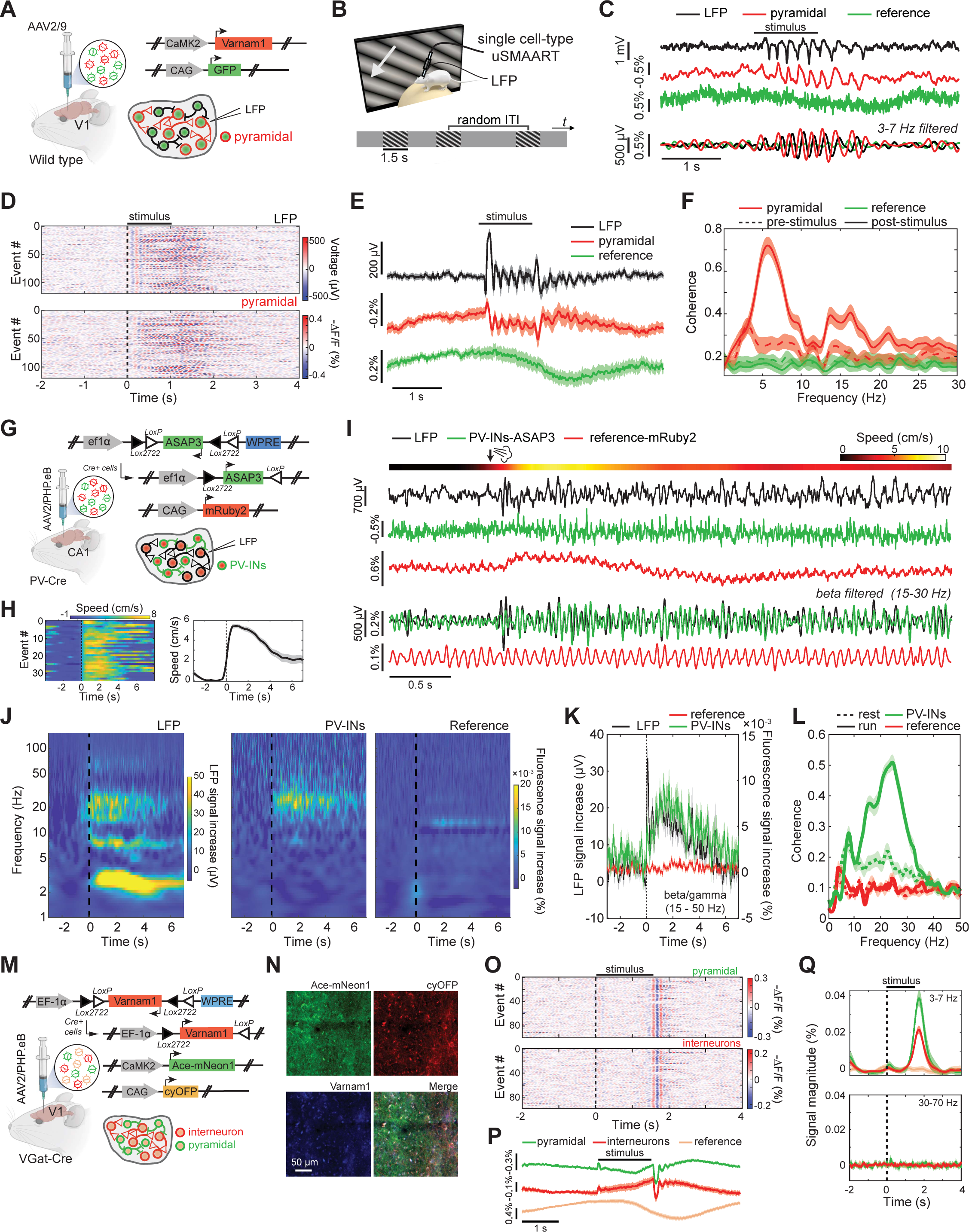
uSMAART captures high-frequency voltage dynamics in the primary visual cortex and hippocampus of head-fixed, awake mice. (**A–F**) uSMAART studies of visually evoked high-frequency voltage dynamics in pyramidal neurons in the primary visual cortex of awake mice. (**A**) Genetic constructs used to virally express the voltage indicator Varnam1 in glutamatergic neurons and the reference fluor GFP in a variety of neuronal cells. (**B**) We provided visual stimuli using the same protocols as in **Figure 3B**. (**C**) Example time traces of LFP, pyramidal cell membrane voltage, and reference fluorescence signals during visual stimulation, along with bandpass-filtered (3–7 Hz) versions of the same traces. (**D**) Plot of the coherences between the Varnam1 and reference fluorescence signals in the pre- (dashed curves) and post-stimulus (solid lines) periods (2-s-epochs). Pyramidal cell membrane voltage signals but not the reference signals exhibited visually evoked increases in coherence with the LFP up to 30 Hz (p<0.05, rank sum test). Shaded area: 95% C.I. (**E**) Raster plots of LFP (top) and pyramidal cell voltage signals (bottom) showing that visual stimuli consistently evoked 3-7 Hz oscillations. Each row shows data from one of 115 different visual stimulation trials in the same mouse. (**F**) Mean time-dependent fluorescence traces, obtained by averaging each of the 3 signals across all 115 trials of (**E**). Shaded area: 95% C.I. (**G–L**) uSMAART studies of how rest-to-run transitions impact beta- and gamma-band activity in hippocampal PV interneurons. We delivered airpuffs to the mouse’s back to induce running. (**G**) Genetic constructs used to virally express the voltage indicator ASAP3 in PV interneurons of hippocampal area CA1 along with the reference fluor mRuby2. (**H**) *Left*: Raster plot showing time courses of locomotor speed by a head-fixed mouse on a running wheel. *Right*: Plot of the mean time-dependent locomotor speed, averaged over 36 rest-to-run transitions. Vertical dashed lines mark the time of airpuff delivery. (**I**) *Top*: Color plot showing the change in locomotor speed induced by an individual airpuff (black arrow). *Bottom:* Time traces of concurrently recorded LFP and fluorescence signals. The top 3 traces are unfiltered, whereas the bottom three are bandpass-filtered (15–30 Hz). Note that the reference signal captures a slow hemodynamic response after the airpuff as well as the heartbeat artifact, which both appear to be absent from the LFP and PV cell membrane voltage traces. (**J**) Mean, time-dependent wavelet spectrograms, showing the locomotor-evoked increase in beta frequency (15–30 Hz) power in both the LFP (*left*) and PV cell (*middle*) recordings but not in the reference signals (*right*). Vertical dashed lines mark the time of airpuff delivery. (**K**) Mean time-dependent magnitudes of the LFP and the two fluorescence signals at high-frequencies (15–50 Hz), across 50 trials in one mouse. Vertical dashed line marks the time of airpuff delivery. Shaded area: s.e.m. (**L**) Plots of the mean coherence between the LFP and the two fluorescence signals during resting (solid curves) and running (dashed curves) periods, across 36 rest-to-run transitions in one mouse. Shaded area: s.e.m. (**M–Q**) Studies of the joint dynamics of excitatory and inhibitory neuron-types in the primary visual cortex of awake mice. (**M**) Local injection of three AAVs of the PHP.eB serotype into the visual cortex of VGat-Cre mice allowed expression of Cre-dependent Varnam1 in GABAergic interneurons, Ace-mNeon1 in pyramidal neurons, and cyOFP, a reference fluorophore. (**N**) Confocal images of a brain slice from a mouse expressing the fluorescent proteins shown in (**M**). (**O**) Raster plots showing the population membrane voltage dynamics of pyramidal cells (*top*) and interneurons (*bottom*) in the same mouse on 96 different trials of visual stimulation. The visual stimuli consistently evoked voltage oscillations in both neural populations. Vertical dashed lines in (**O,Q**) mark the time of visual stimulus onset. (**P**) Mean time-dependent fluorescence traces, obtained by averaging the signals from all three fluors across all 96 trials of (**O**). Shaded area: 95% C.I. (**Q**) Mean time-dependent fluorescence signal magnitudes in the 3-7 Hz (*upper panel*) and gamma (30–70 Hz; *lower panel*) frequency bands for all three fluors. Shaded area: 95% C.I.

**Figure S3:**
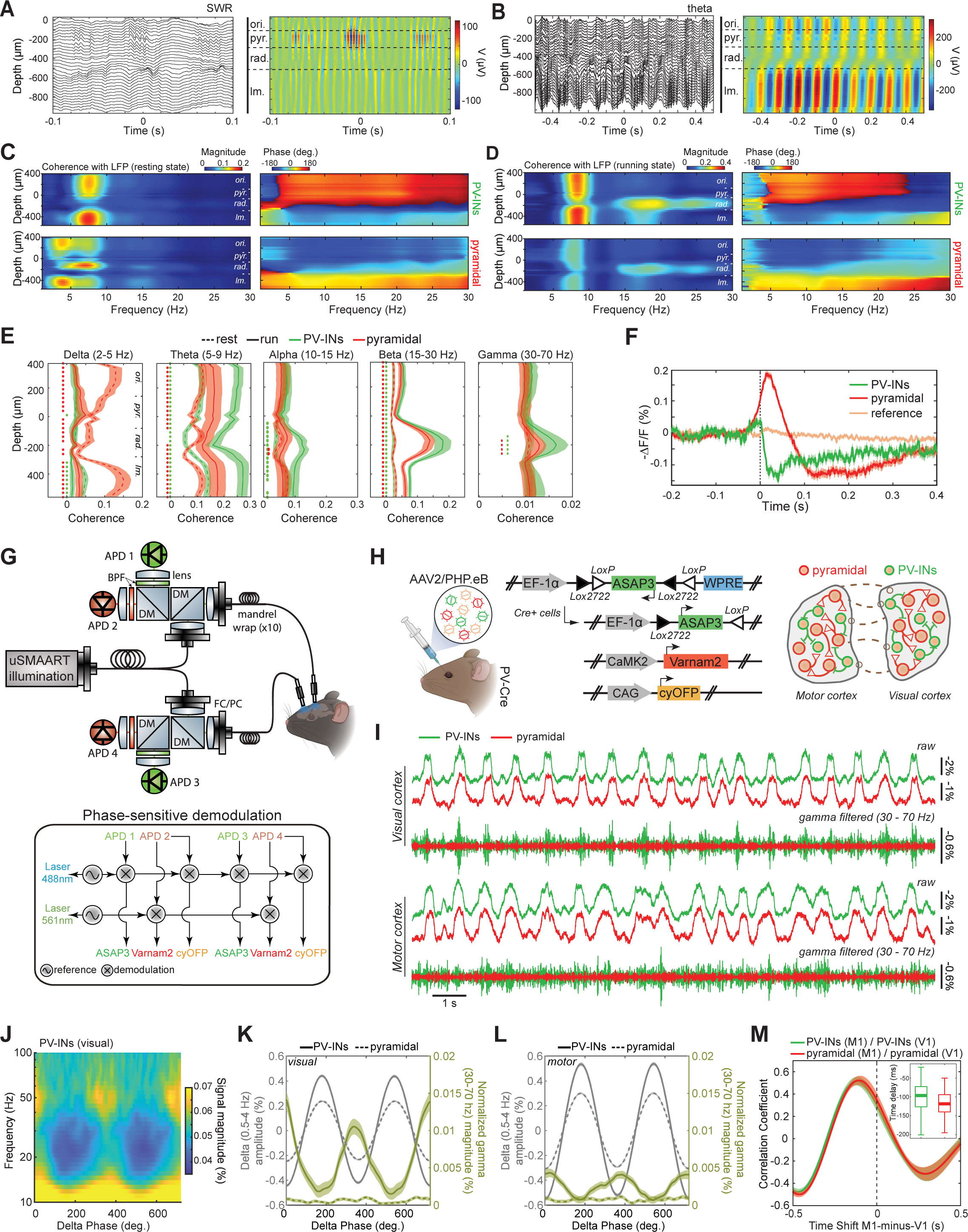
uSMAART studies of the dynamic interplay between multiple neuron-types. (**A, B**) To identify the boundaries between the different layers of CA1 hippocampus in our 32-site silicon probe recordings, we examined the voltage traces from each of the recording sites during intervals in which there were either sharp-wave ripple (SWR) events or theta oscillations. The left panels of (**A**) and (**B**) show illustrative, raw unfiltered LFP traces for each of the 32 recording sites, plotted with the tissue depth of each recording noted on the *y*-axis, as measured from the most dorsal recording site. An example SWR event is shown in (**A**), and a bout of theta oscillations is shown in (**B**). The right panels have color plots of the same data shown at left but temporally bandpass-filtered across either the ripple (120–180 Hz; **A**) or theta frequency (5–9 Hz; **B**) range. We identified the 4 canonical hippocampal layers using two electrophysiological landmarks: (1) the center of *stratum pyramidale* corresponds to the depth at which ripple oscillations have the greatest amplitude; (2) the anatomical boundary between *stratum radiatum* and *stratum lacunosum moleculare* corresponds to the depth at which theta oscillations change in both amplitude and phase. (**C, D**) Plots of the magnitude (*right* column) and phase (*left* column) coherence between PV and pyramidal cell fluorescence voltage signals and the LFP across the 32 recording sites of the silicon probe (plotted as a function of depth relative to the most dorsal recording site) during resting (**C**) and running (**D**) periods. (**E**) Depth-dependent average coherence magnitude between each of the 32 recording sites of the silicon probe and PV and pyramidal cell TEMPO signals, across 5 different frequency bands, as extracted from the data of (**C, D**). During rest, PV and pyramidal cells exhibited delta frequency rhythms that had greater coherence with the LFP than during locomotion, especially in *stratum oriens* and *stratum radiatum*. During locomotion but not rest, PV and pyramidal cells exhibited beta and gamma frequency rhythms that were coherent with the LFP, especially in *stratum pyramidale*. Green and red dots indicate statistically significant differences (p<0.01, two-sample rank sum test) between the coherence values associated with rest and locomotion. Averages were performed across n=26 and n=41 epochs (each 30 s in duration) of rest and locomotion, respectively. Shaded areas: 95% C.I. (**F**) Mean time-dependent traces of the fluorescence voltage signals for CA1 hippocampal PV interneurons and pyramidal cells during ripples, as in **Figure 3Q** but for another mouse. Shading: s.e.m. (**G–M**) uSMAART allows concurrent optical voltage recordings of two neuron-types in each of two different brain areas. (**G**) Diagrams of the optical pathway (*top*) and electronic signal demodulation scheme (*bottom*) used to monitor two neuron-types in each of two different areas. (**H**) Retro-orbital injection of three PHP.eB viruses into PV-Cre mice enabled expression of Cre-dependent ASAP3 in PV interneurons, Varnam2 in pyramidal neurons and cyOFP in all neuron-types. (**I**) Example fluorescence voltage traces for PV and pyramidal cells in visual V1 and motor M1 cortical areas of a ketamine-xylazine-anesthetized mouse. For each brain area, traces in the top set are unfiltered, whereas traces in the bottom set are bandpass-filtered (30-70 Hz). Note that in V1, it is apparent that high-frequency oscillations coincide with up-state depolarization events. (**J**) Wavelet spectrogram of the activity of the PV interneurons in V1, averaged across cycles of the delta oscillation (using 0.9 Hz as the frequency of peak delta power). Bursts of gamma frequency (30–70 Hz) power arose at depolarized voltages of the delta rhythm. In **J–L**, the convention used for the phase of the delta oscillation is that 0-deg. refers to the trough of the oscillation, *i.e*., the greatest hyperpolarization in the TEMPO signal. (**K, L**) Average signal amplitudes filtered at the delta frequency (*black curves,* left axis) and magnitude of gamma frequency (*olive curves,* right axis) estimated for each cell-type in V1 (**K**) or M1 (**L**). Note that ASAP3-labeled PV interneurons (*solid curves*), but not Varnam2-labeled pyramidal neurons (*dashed curves*), displayed delta-gamma CFC in both areas, with CFC in V1 being stronger than in M1. The observed difference between the two neuron-types was likely due to the distinct signaling ranges of the two GEVIs used. Shading: s.e.m. (**M**) Plots of the mean time-dependent cross-correlation coefficient between the two areas V1 and M1 quantified using the voltage dynamics of either excitatory or inhibitory neuron-types in each brain area. For PV cells, anesthesia-induced delta rhythms in M1 exhibited a –103 ± 41 ms (mean ± s.d.) phase lead over the delta rhythms in V1, whereas in pyramidal cells the phase lead was –117 ± 35 ms ms. Shading: s.e.m. over 60 samples, each 10 s in duration. *Inset*: Box-and-whisker plot of the differences in the times of the peak correlation coefficient between V1 and M1, for PV (green) and pyramidal (red) cells. Horizontal lines denote medians, boxes cover the middle two quartiles, and whiskers extend to the 10th and 90th percentiles.

**Figure S4:**
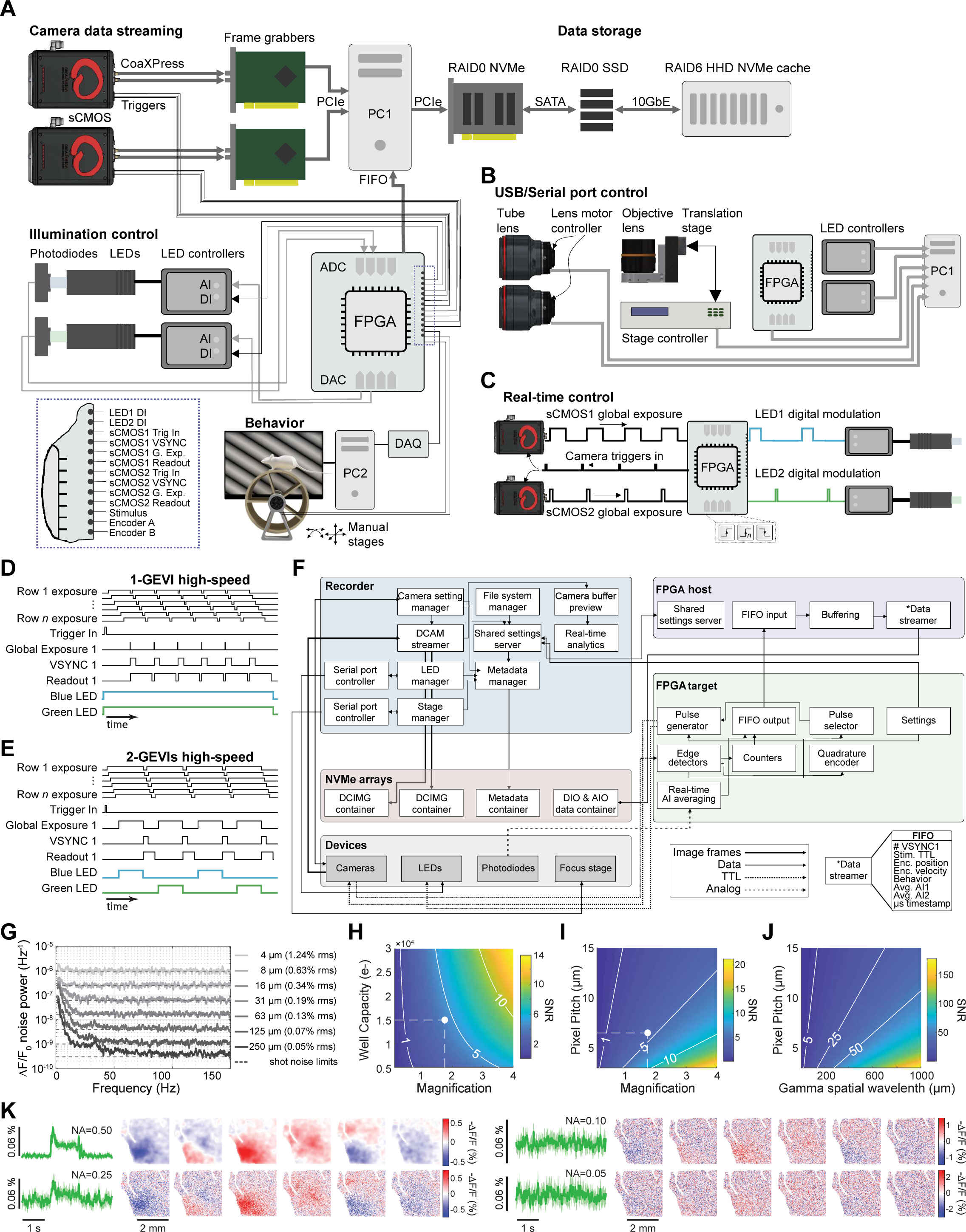
Electronics, controllers and software for the TEMPO mesoscope. (**A**) Each of the two sCMOS cameras (Orca Fusion; Hamamatsu) streams image frame data at 0.8 GB/s to their respective frame grabbers installed on a single personal computer (PC) via the CoaXPress interface. A RAID0 hardware array of Non-Volatile Memory Express (NVMe) hard drives on PCIe stores incoming frames in real-time. For each frame, the FPGA transmits structured data back to the computer using a FIFO queue framework (see (**F**)). After each imaging session, we stored and processed the data on a RAID0 SSD array on the same PC. We backed up the raw data on a RAID6 HDD array connected to the PC via a 10 GbE optical link. We used a field-programmable gate array (FPGA) board to control the cameras’ input and output triggers as well as the LEDs pulsing protocol (see (**D-E**)). *Inset*: expanded view of all digital signals received or sent by the FPGA board. For dual cell-type mesoscope TEMPO measurements, we monitored the LEDs power with photodiodes, whose signals were sent to the analog input ports of the FPGA. For visual stimulation protocol (**Figure 5, 7**), we used another PC with its own DAQ system (see **STAR Methods**) and recorded the stimulation timing on the FPGA board. We monitored the mouse’s position using a quadrature rotary encoder whose pair of encoders A and B were routed to the FPGA. (**B**) The TEMPO mesoscope uses a combination of USB and serial port communication to control its components from a single PC workstation, which facilitates streamlined operation and synchronization of the entire system. The focusing mechanism in each tube lens (see **Figure 4A**) is controlled via a serial port, as is the driver for the stage adjusting the axial position of the objective lens. The FPGA board communicates with the main PC via USB. The LED controllers use a serial port. (**C**) To achieve uniform illumination and prevent spatially dependent flickering, it is crucial to precisely trigger the LED pulses within the time period when all camera rows are simultaneously illuminated. For dual cell-type TEMPO mesoscope recordings, real-time TTL pulses controlled the LEDs based on the global exposure of the cameras operating in rolling shutter mode. The FPGA board generated a camera trigger and created an interleaved pulse sequence that matched the camera exposure precisely. To ensure uniform illumination, the FPGA board triggered either the green or blue LED, using every other pulse from the global exposure signal provided by the camera. (**D**) Triggers and digital pulse sequences for experiments with one GEVI. Both cameras are triggered simultaneously with one TTL pulse and operate in rolling shutter mode, with a frame exposure duration that maximizes the frame acquisition rate for the chosen field-of-view size and duty ratio of the frame imaging time, *i.e.* avoiding the sensor reset time. To synchronize the recording on the sub-frame level, we monitor different types of triggers that have different meanings. The camera operates in the rolling shutter mode, in which all camera rows are exposed simultaneously for a very short time. To control this precisely, the camera outputs the global exposure trigger signal. The camera outputs are also monitored via the VSYNC and readout triggers. The VSYNC signal indicates the end of a frame and the start of a new one and is vital for achieving accurate synchronization of the data. The two cameras themselves are not directly synchronized to each other, but we routinely use an oscilloscope and logic analyzers to test that no misalignment occurs during the duration of typical recordings (<10 min). (**E**) Schematic of the timing protocol used for dual neuron-type recordings. To capture sufficient fluorescence light in this modality, it is crucial to extend the exposure time of both cameras beyond their default values while staying within the global exposure period. However, the trade-off is that longer exposure times lead to a slower frame rate, and we are limited by the energy of the illumination pulse. Thus, careful optimization is needed to find the optimal exposure time that balances the needs for sufficient photon capture and for having a reasonable frame rate. This implies that the frame rate is slower than that allowed in principle by the field-of-view sampled on the camera chips. Using global exposures of both cameras as depicted in (**C**), we created interleaved pulses of blue and green LED illumination. The interleaved pulses allowed us to separate the signals from each GEVI using a long-Stokes shift reference fluorophore (cyOFP). (**F**) The high-level architecture of software and data flow of the system comprises 3 main programs: the recorder graphical user interface, FPGA host software, and FPGA target software. The Recorder program acquires, processes and stores data in real-time. The FPGA host software communicates with the FPGA target software on the FPGA board, which controls the devices involved in data transfer and real-time control, such as the cameras, LEDs, photodiodes, and focus stage. We stored data in real-time on NVMe arrays, which allowed us to perform continuous recording for nearly 1 h. (**G**) To study the effect of spatial averaging on high-frequency noise, we used a control mouse implanted with a 7-mm window that expressed GFP and mRuby2 in all cell-types. We imaged this mouse at 300 Hz (**Figure 4A**) and unmixed the signals in the green and red channel movies using convolutional unmixing (**Figure S5**). Plotted are the power spectral densities of the single pixel noise when computed using different levels of spatial averaging. Normally, we spatially averaged voltage and reference signals over 31 µm (8 × 8 camera pixels) of brain tissue, as we found this to be an effective compromise that yielded reasonable computational speed, sensitivity to high-frequency phenomenon, and spatial accuracy of the unmixing procedure. (**H–K**) We quantified the ability of the TEMPO mesoscope to capture traveling gamma oscillations as a function of different system parameters, such as the camera well capacity, the camera pixel size and the system magnification. We also considered the spatial wavelength of the high-frequency voltage dynamics as an input parameter. (**H-J**) Theoretical estimates of the wave detection signal-to-noise ratio (SNR), highlighting the influence of different parameters of the optical system and the sCMOS camera chip. The plots show the interactions between three different pairs of parameters: the optical magnification and the camera’s pixel well capacity (**H**); the magnification and the camera pixel pitch (**I**); the wavelength of the traveling gamma wave and the pixel pitch (**J**). With our system’s magnification of 1.75×, well capacity of 15,000 electrons, and NA of 0.47, the resulting single-pixel level SNR was about 4.28 (marked by the white points in each panel). This value is sufficient to quantify the dynamics of a single traveling gamma wave, as shown in **Figure 5**. Jointly, these panels suggest that, to detect high-frequency propagating voltage waves, it is best to use a modest magnification over the brain areas of interest together with an sCMOS camera with a small pixel size and large well capacity. (**K**) Detection of propagating gamma events (**Figure 5**) strongly depends on the voltage signal SNR, which in turn depends on the numerical aperture (NA) of the microscope objective lens. The NAs considered in this set of plots are 0.5 (the actual NA value of our lens), and then 0.25, 0.1, and 0.05, which respectively led to declines in SNR by factors of 2, 5, and 10. The effects of having different SNRs were mimicked by computationally adding noise to the panels of the top row, reducing effective SNR levels.

**Figure S5:**
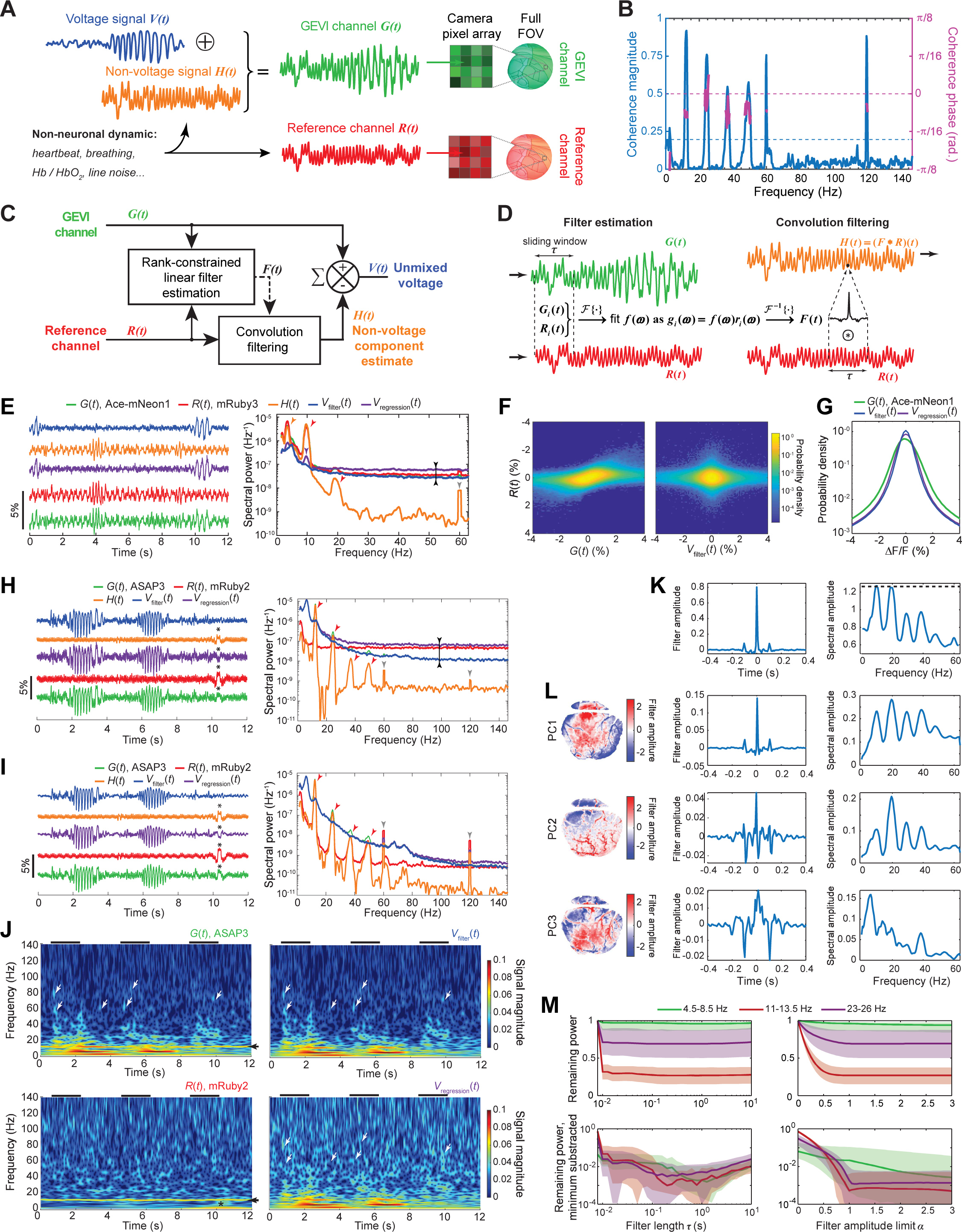
Convolutional unmixing removes biological and instrumentation artifacts from neural voltage signals in a frequency-dependent manner. (**A**) A model of signal content used for the estimation and unmixing of artifacts from the GEVI fluorescence channel. Hemodynamics and other artifacts are present in both fluorescence channels, but neural voltage signals are present only in the fluorescence voltage channel. The diagram refers to studies with a single GEVI; when using two GEVIs, we unmixed the reference channel content from each voltage channel independently. (**B**) Plots of the frequency-dependent, normalized amplitude (blue curve) and phase (magenta curve) of the coherence between the voltage and reference fluorescence channels, before unmixing and averaged over brain area V1 in an example mouse expressing ASAP3 in PV interneurons and mRuby2 in all neuron-types (same mouse as (**H–J,M**)). Phase values are shown only for frequencies at which the coherence amplitude is >0.2 (horizontal dashed blue line). The two fluorescence channels display high coherence peaks at the fundamental (∼12 Hz) and harmonic frequencies of the heartbeat, as well as the fundamental (60 Hz) and second-harmonic (120 Hz) frequencies of the electric power line noise. Although these stereotypical oscillations are highly coherent between the GEVI and reference channels, the phases are variable and differ from zero (dashed purple line), and a simple linear regression approach is unable to capture non-uniform phase lags. (**C**) Block diagram of the unmixing strategy. A rank-constrained linear filter, *F*(*t*), is estimated using both fluorescence channels and captures the manner in which artifacts seen in the reference channel coherently invade the voltage channel in different frequency bands (*e.g.* as in (**B**)). *F*(*t*) is convolved with the reference channel, *R*(*t*), to attain an estimate, *H*(*t*), of the non-voltage signals present in the GEVI channel. *H*(*t*) is subtracted from the GEVI channel trace, *G*(*t*), to attain an estimate of the true voltage signals, *V*(*t*). Plain arrows represent data flow. Dashed arrow denotes transfer of the *F*(*t*) function. (**D**) Schematic of the filter estimation and convolution steps in (**C**) (see also **STAR Methods**). We used the Fourier transform *F*{.} to obtain the frequency-domain representations, *r*(ω) and *g*(ω), of *R*(*t*) and *G*(*t*), respectively, over sliding time windows of duration, τ (typically 0.5–2 s, see (**M**)). We then used linear regression to compute for each frequency, ω, the value of the filter coefficient, *f*(ω), that best describes the amplitude with which the reference signal, *r*(ω), is present in the voltage channel, *g*(ω), across the set of all time windows. After performing an inverse Fourier transform of *f*(ω) to obtain the filter’s time-domain representation, *F*(*t*), we estimated *H*(*t*) by convolving *F*(*t*) with *R*(*t*) and then subtracted *H*(*t*) from *G*(*t*) to obtain *V*(*t*). (**E**) *Left*: An example set of raw, *G*(*t*) and *R*(*t*), and unmixed, *V*_filter_(*t*) and *V*_regression_(*t*), fluorescence Δ*F/F* traces for the Ace-mNeon1 voltage indicator and mRuby3 reference fluor, obtained from an Ai218 transgenic mouse crossed with a PV-Cre mouse (**Figure S6**). Traces are colored to match the convention used in (**A–D**) and *R*(*t*) was scaled (by 1.8-fold) to match the power at the heartbeat frequency (that is the dominant shared contribution, ∼10 Hz) between *G*(*t*) and *R*(*t*). Hemodynamics are visible in *R*(*t*) (red trace) throughout the recording segment and contaminate the raw voltage trace (green) but are reduced after unmixing using either standard linear regression (purple trace) or our filter-based approach (blue trace). Notable voltage signals appear just after ∼10 s and remain after unmixing. *Right*: Power spectral density plots for all 5 traces shown in the left graph, computed across the full recording duration. After unmixing using our filter-based method, the voltage signals (blue curve) lack several spectral peaks that arise from heartbeat-related artifacts (∼9 Hz and ∼18 Hz harmonic; red arrowheads), breathing-related artifacts (∼4 Hz; orange arrowhead) and electric line noise (60 Hz; light gray arrowhead). Unmixing with a standard linear regression (purple curve) does not fully remove the heartbeat and breathing artifacts, since a standard regression does not account for amplitude and phase differences in the heatbeat content between the two fluorescence channels (see (**B**)). Moreover, the linear regression introduces high-frequency (>10 Hz) noise from the reference channel into the voltage channel, as seen via the increase in baseline high-frequency spectral power after unmixing (marked with opposing black arrowheads). By comparison, after the filter-based unmixing approach, the power at high-frequencies remains unchanged after unmixing. (**F**) Heat maps showing the joint probability distribution of fluorescence signals in the voltage and reference channels before (*left*) and after (*right*) filter-based unmixing across the full recording session (9 min) of (**E**). After unmixing, the two channels are less correlated. (G) Probability distributions of signal fluctuations for the traces used in (**F**), confirming that filter-based unmixing reduces the variance level in the voltage trace, *V*_filter_(*t*). Note that unmixed traces *V*_filter_(*t*) and *V*_regression_ (*t*) have smaller variances as compared to the raw trace *G*(*t*). (**H, I**) Plots in the same format as those in (**E**), for studies using the ASAP3 voltage indicator, as computed for a single image pixel in area V1, (**H**), or averaged over the entirety of V1, (**I**). Colored and black arrowheads mark the same spectral features noted in (**E**). Asterisks mark an artifact at ∼10.5 s that is likely to result from brain motion. (**J**) Wavelet spectrograms of the traces *G*(*t*) and *R*(*t*) from (**I**) (*left column*), and of the estimated traces of the voltage signal obtained by standard regression, *V*_regression_(*t*) (*lower right*) or convolutional filtering, *V*_filter_(*t*), (*upper right*), estimated for a single image pixel in area V1. Black horizontal bars above the plots mark periods of visual stimulation. Both unmixing approaches (right column) removed the heartbeat artifact and its harmonic (∼11 and 22 Hz; marked with black arrowheads), but only convolution filtering removed a low-frequency artifact (at time ∼10.5 s and frequency <5 Hz, marked with a black asterisk as in (**H-I**)), which is likely to result from brain motion. Notably, because linear regression adds high-frequency noise to the estimated voltage trace, only convolution filtering preserves the visibility of visually evoked high-frequency voltage transients (marked with white arrowheads) that were absent in the reference channel (*bottom left*). (**K**) Plots of the filter, *F*(*t*), in the time (*left*) and frequency (*right*) domains, averaged across the whole imaging window. The horizontal dashed line in the right plot represents the unmixing coefficient as determined by linear regression estimated on the fundamental frequency of the heartbeat artifacts. (**L**) To characterize how *F*(*t*) varied across the cortical surface, we performed a principal component analysis (PCA) for the example mouse used in (**E**). Plots show the spatial variations (*left column*), time-dependence (*middle column*) and frequency-domain power density (*right column*) for the first three principal components (PCs) of *F*(*t*). The first PC is mainly uniform across the field-of-view, except for the periphery. The second and third PCs capture a gradient of variation across the field-of-view and distincts sets of blood vessels. The PCs have distinct signatures in the frequency domain, highlighting their distinct hemodynamic properties. (**M**) Plots of the relative signal powers that remain after unmixing as a function of the hyperparameters τ (filter length; *left column*) and α (filter amplitude limit; *right column*) in a given frequency band. The dependencies are plotted for 3 frequency ranges: low-frequency visually evoked voltage signals (4.5–8.5 Hz), and the fundamental (11–13.5 Hz), and first harmonic (23–26 frequencies Hz) of the heartbeat. For individual pixels in the GEVI channel *G*(*t*), we computed the power in each of the frequency bands as a function of both hyperparameters, either by normalizing by their respective initial values (*top row*) or by normalizing and then subtracting each pixel’s minimum value (*bottom row*). The solid curves show median values across all pixels for every value of the hyperparameter, and the shaded areas indicate the 0.05–0.95 interquartile range. In each case, we first estimated the convolutional filter *F*(*t*) on a training dataset and then applied it to perform unmixing on the testing dataset (50/50 split between train and test). Note that, in the bottom right plot, the heartbeat-related powers decline rapidly until α∼1 and stay constant for higher values of α. However, the power in the voltage frequency band flattens for α∼1 but then decreases for higher values, suggesting over-subtraction of the GEVI signals due to excessive amplification of the filter’s coefficients at this specific frequency band due to cross-channel coherence. Typical values of τ are τ ∼ 0. 5–2 *s*, depending on the dataset. We used τ = 1 *s* for the plots in the right column. A typical value for α is 1.2, meaning that no coefficient values of the convolution filter *f*(ω) will be more than 20% greater than the linear regression coefficient (*e.g.*, 1.25 in (**K**)) for the fundamental hemodynamic frequency. The dataset used is the same as that in (**H-I**).

**Figure S6:**
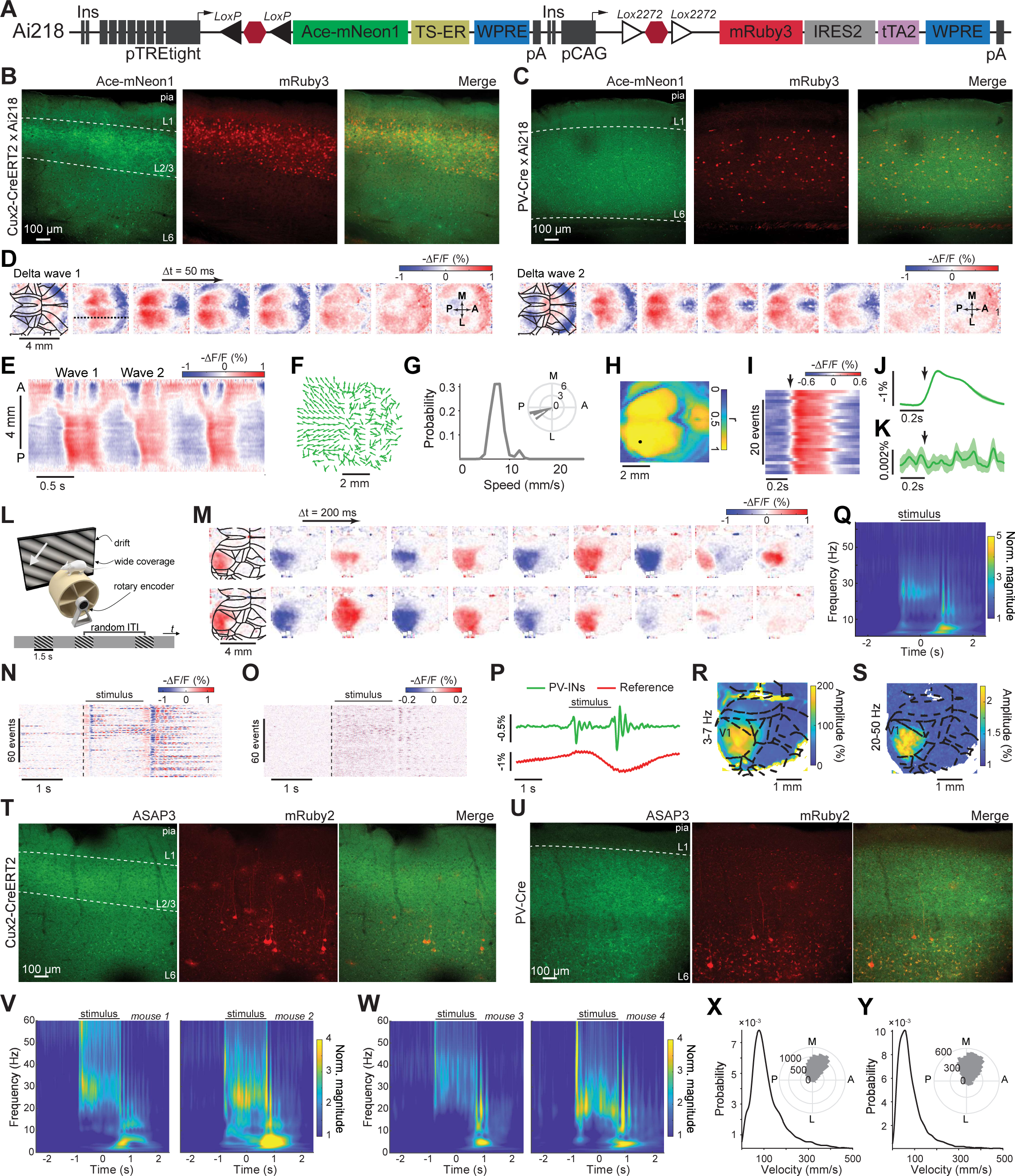
Construction and validation of mouse transgenic and viral methods for fluorescence labeling that were expressly designed for TEMPO studies. (**A**) To co-express the FRET-opsin voltage sensor Ace-mNeon1 and the reference fluor mRuby3 in specific cell-types, we generated a Cre-dependent Ai218 reporter mouse line expressly designed for TEMPO studies (see **STAR Methods**). The genetic construct is inserted in the TIGRE2.0 mouse locus. To express Ace-mNeon1 and mRuby3 in targeted neuron-types, we crossed the Ai218 reporter mice with cell-type-specific Cre-driver lines. (**B, C**) Confocal fluorescence images of a mouse brain slice showing the cortical expression patterns of Ace-mNeon1 and mRuby3 in mice that are genetic crosses of Ai218 with either a Cux2-Cre^ERT2^ mouse line, (**B**), to label layer 2/3 pyramidal cells, or a with PV-Cre line, (**C**), to label PV interneurons. Approximate neocortical layers are marked to aid visualization (L1: layer 1, L2/3: layer 2/3 and L6: layer 6). (**D**) Two examples of traveling neocortical voltage waves in the delta frequency band, shown in sequences of Ace-mNeon1 images (50 ms between successive images) taken at 130 Hz in a ketamine-xylazine-anesthetized Ai218 × Cux2-Cre^ERT2^ mouse. Images underwent unmixing (**Figure S5**) to remove hemodynamic changes captured in the mRuby3 channel but were not otherwise filtered. Brain area boundaries are superposed on the first image in each sequence. In each case, a voltage depolarization (denoted by red hues) sweeps across the cortex in the anterior to posterior (A-P) direction. For display purposes only, the images shown were spatially low-pass filtered using a Gaussian filter (156 μm FWHM). (**E**) Color plot (top) showing the anterior to posterior propagation of the two traveling waves in (**D**). To create the plot, at each time point (*x*-axis) and for each A-P coordinate (*y*-axis), we averaged the measured fluorescence ΔF/F values along the medio-lateral direction. (**F**) Flow maps showing the local propagation directions of voltage depolarization for the individual delta wave #1 observed in (**D, E**). Flow vectors are all normalized to have the same length. (**G**) Distributions of delta wave propagation speed across all delta events seen in the Ai218 × Cux2-Cre^ERT2^ mouse, computed near the center of area V1 (marked by a black dot in (**H**)), where wave propagation was consistently anterior to posterior. *Inset*: Polar histogram showing the distribution of wave propagation direction for the same mouse at the same location, revealing an approximate alignment of wave propagation with the A-P axis. (**H**) Maps of peak correlation coefficients, *r*, for a Ai218 × PV-Cre mouse computed for each spatial point by calculating the temporal correlation function between the local fluorescence trace and that at the center of V1 (black dot) and then finding this function’s maximum value. (**I–K**) Raster plots of down-to-up transition events in the ΔF/F voltage signal in area V1 (**I**), along with plots of the mean time-dependent activity obtained by averaging over 29 transition events in the delta frequency (0.5-4 Hz) (**J**) or gamma frequency (30–60 Hz) bands (**K**). At the peaks of the delta waves, we observed no activity in the gamma frequency band, unlike in our studies of anesthetized mice using the ASAP3 indicator. (However, see (**N**) below). The dark arrows mark the onset of the down-to-up state transition. Shaded area: 95% CI. (**L**) We used the same visual stimulation approach as in **Figure 3B**. (**M**) Two example sequences of fluorescence images from the same Ai218 × PV-Cre mouse, showcasing the spatiotemporal dynamics of visually evoked 3-7 Hz oscillations in PV interneurons in area V1. Successive images shown in the sequences were taken 200 ms apart). Brain area boundaries, based on the Allen Brain Atlas (**Figure 4A** inset), are superposed onto the first image in each sequence. For display purposes only, the images shown were spatially low-pass filtered using a Gaussian filter (156 μm FWHM). (**N**) Raster color plots showing fluorescence voltage signals, averaged over V1, revealing 3-7 Hz oscillations locked to stimulus-offset in V1 (80 stimulus trials shown). (**O**) Raster plots of gamma-band filtered fluorescence voltage activity in V1, showing that gamma-band activity was evoked during stimulus presentation in V1 (80 stimulus trials shown). (**P**) Mean time-dependent traces of PV interneuron voltage activity (green trace) and reference fluor signals (red trace), obtained by averaging the unfiltered fluorescence signals of panel (**N**) over all 80 trials. Shaded area: s.e.m. (**Q**) Stimulus-triggered average wavelet spectrogram of the PV cell fluorescence voltage traces from (**N-P**), revealing the high-frequency voltage activity during visual stimulus presentation, plus the 3-7 Hz oscillations that arose at stimulus offset. (**R, S**) Spatial maps showing the magnitudes of visually evoked 3-7 Hz (**R**) and gamma-band (**S**) activity at their peaks within the averaged trial. The plots illustrate that the visually evoked voltage signal was confined within the visual system. (**T, U**) Fluorescence confocal imaging of brain slices from a Cux2-Cre^ERT2^ mouse expressing mRuby2 in all cell-types and ASAP3 in layer 2/3 pyramidal neurons (**T**), and a PV-Cre mouse expressing mRuby2 in all cell-types and ASAP3 in PV interneurons (**U**). Images were acquired from the sensory neocortex atop the dorsal hippocampus. The mouse preparations are the same as for **Figures 4, 5**. L1: layer 1, L2/3: layer 2/3, L6: layer 6. (**V, W**) Event-related wavelet spectrograms for the two PV-Cre mice (**V**) and Cux2-Cre^ERT2^ mice (**W**) used in **Figure 5G-H**. (**X, Y**) Histograms showing distributions of 3-7 Hz wave propagation speed (main plots) and direction (inset plots) in area V1 for a PV-Cre mouse (Mouse 1 in **Figure 5G-H**), determined from either the raw data (**X**), or by computing wave speeds using gamma-band filtered (35–100 Hz) versions of the same movie data (**Y**; n = 50 wave events). The similar speeds and directions evidenced in the two sets of histograms suggest that the high- and low-frequency content are both part of the same phenomenon, *i.e.* with the gamma frequency activity constituting harmonics of a 3-7 Hz carrier wave. Importantly, the speeds and directions of the gamma waves analyzed here are notably distinct from those of the isolated gamma waves shown in **Figure 5**.

**Figure S7:**
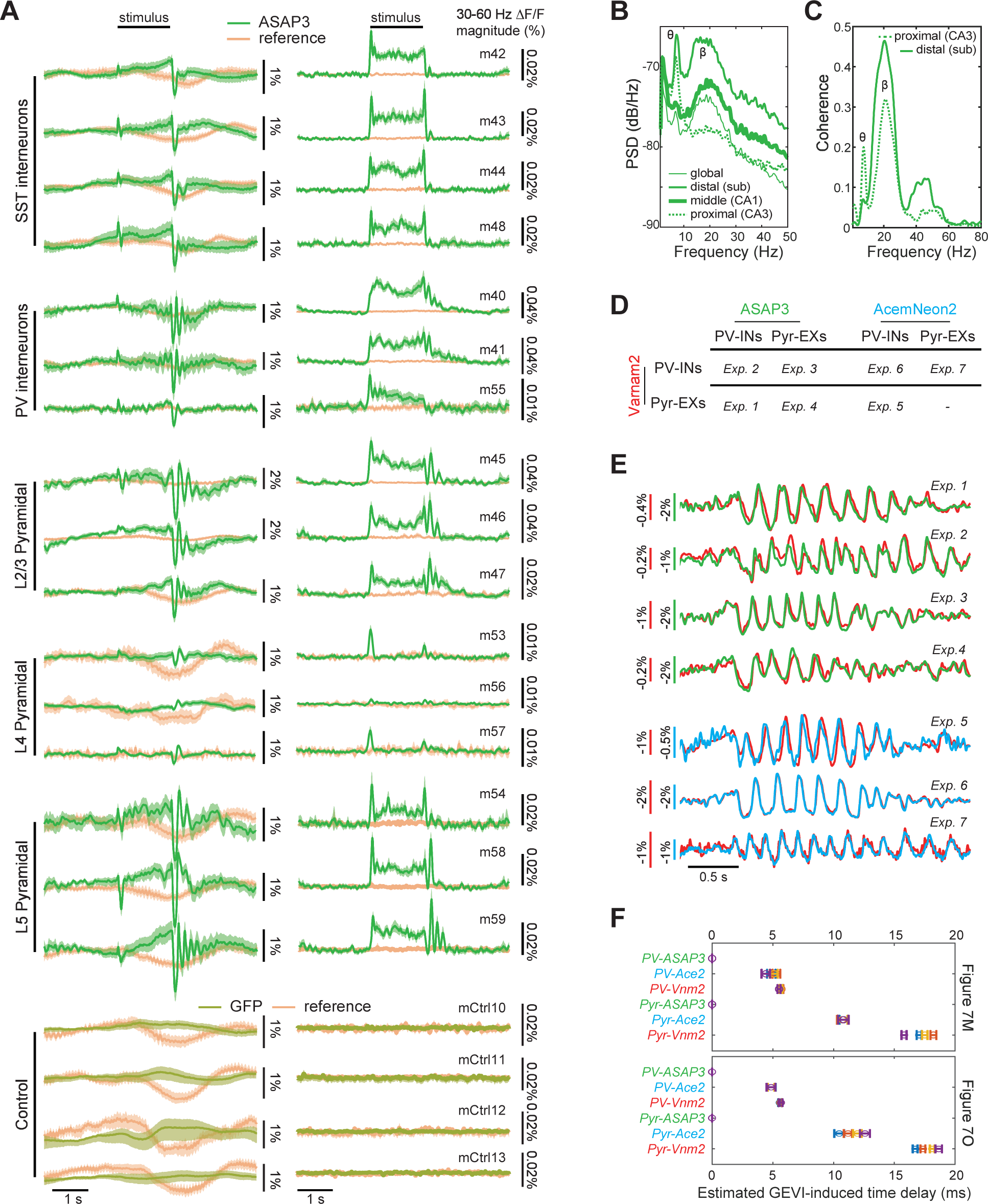
Voltage signal variations with neuron class and indicator-type. (**A**) Similar to **Figure 5G-H**, plotted for mice in which ASAP3 labeled cortical SST-interneurons (4 mice), PV-interneurons (3 mice), layer 2/3 pyramidal neurons (3 mice), layer 4 pyramidal neurons (3 mice), and layer 5 pyramidal neurons (3 mice). Mean fluorescence traces are shown in the left plots; mean fluorescence signal magnitudes in the gamma-band (30–60 Hz) are shown in the right plots. Visually evoked gamma- and 3-7-Hz-band activity are respectively visible, to varying extents across the different cell-types, during stimulus presentation (black horizontal bars) and following stimulus offset. Note, for example, the near absence of oscillatory activity except near the times of stimulus onset and offset. Traces for the mRuby2 reference fluor are included in all plots. The bottom 4 plots show results from control mice, in which the static green fluorescent marker, GFP, was used instead of the green ASAP3 voltage indicator. Shading: 95% C.I. (**B**) Plots of the power spectral density of PV-cell fluorescence voltage signals, acquired as in **Figure 6B–D** by TEMPO imaging in hippocampus, at 3 different parts of the 2.1-mm-diameter field-of-view (*proximal*, *middle*, *distal*) along the CA1-CA3 axis or across the full field-of-view (*global*). (**C**) Plot of the coherence between the LFP, acquired in the same mouse as in **B** at the CA1 side of the field-of-view, and PV cell voltages at proximal and distal positions in the visible field in area CA1. (**D, E**) Table, (**D**), summarizing the voltage-indicator labeling configurations used across 7 different experiments using fiber-optic TEMPO to study relative time delays in the visually evoked 3-7 Hz oscillations of PV-interneurons and/or pyramidal neurons in neocortical area V1 (*c.f*. **Figure 7M,O**). Representative voltage traces from each experiment, (**E**), show subtle differences in the measured temporal shifts across the sets of paired traces that depended upon the assignments of 3 different voltage-indicators (Ace-mNeon2, ASAP3, Varnam2) to the 2 neuron classes. Traces are colored according to the color scheme shown in (**D)**. (**F**) We determined the temporal shifts in the fluorescence reports of oscillatory activity that arise from voltage-indicator kinetics by performing parametric fits to the values of the time-delays found in the data from **Figure 7M–P** and in the 7 experiments of **D-E** (see **STAR Methods**). The plots show the mean ± s.d. temporal shifts induced by each of the 6 possible assignments of the 3 indicators to the 2 neuron classes. The top and bottom plots show time shift values determined using the data from the 4 imaging sessions of **Figures 7M** and **7O**, respectively, which had reversed labeling assignments. The colors of each data point match those used to label the 8 corresponding imaging sessions (n=7 mice) in each of **Figure 7M,O**. Notably, the estimated time-shifts arising from indicator kinetics are nearly invariant across the top and bottom plots, arguing that the values obtained do not vary much across mice. We estimated s.d. values across 100 different samplings of the data, each containing 100 different oscillation events.

